# Hierarchical imaging and computational analysis of three-dimensional vascular network architecture in the entire postnatal and adult mouse brain

**DOI:** 10.1101/2020.10.19.344903

**Authors:** Thomas Wälchli, Jeroen Bisschop, Arttu Miettinen, Alexandra Ulmann-Schuler, Christoph Hintermüller, Eric P. Meyer, Thomas Krucker, Regula Wälchli, Philippe Monnier, Peter Carmeliet, Johannes Vogel, Marco Stampanoni

## Abstract

The formation of new blood vessels and the establishment of vascular networks are crucial during brain development, in the adult healthy brain, as well as in various diseases of the central nervous system (CNS). Here, we describe a method that enables hierarchical imaging and computational analysis of vascular networks in postnatal- and adult mouse brains. Resin-based vascular corrosion casting, scanning electron microscopy, synchrotron radiation and desktop µCT imaging, and computational network analysis are used. Combining these methods enables detailed visualization and quantification of the three-dimensional (3D) brain vasculature. Network features such as vascular volume fraction, branch point density, vessel diameter, - length, -tortuosity, and -directionality as well as extravascular distance can be obtained at any developmental stage from the early postnatal to the adult brain. Our method allows characterizing brain vascular networks separately for capillaries and non-capillaries.

The entire protocol, from mouse perfusion to vessel network analysis, takes approximately 10 days.

**Online summary:** This protocol uses vascular corrosion casting, hierarchical synchrotron radiation µCT imaging, and computational image analysis to assess the three-dimensional vascular network architecture.

## INTRODUCTION

The growth of tissues and organs during embryonic and postnatal development, tissue homeostasis in adulthood, as well as tumor growth, require adequate vascularization via the establishment of complex three-dimensional (3D) vascular networks and subsequent tissue perfusion and oxygenation^1–3^. In these physiological and pathological conditions, the formation of new blood vessels can occur via different mechanisms^1,2,4–6^. Vasculogenesis is defined as the *de novo* formation of blood vessels occurring throughout life, being most active during embryonic and early postnatal development, via differentiation of bone-marrow – respectively blood islets – derived angioblasts (endothelial progenitor cells) into endothelial cells^1,2^. Sprouting angiogenesis describes the formation of new blood vessels from pre-existing ones^2,3^, relying on a number of spatiotemporally tightly regulated cellular mechanisms involving cell adhesion and -spreading, sprout initiation and -formation, tip cell migration, stalk cell proliferation, tube formation, and vessel anastomosis^1,2^. Another mechanism of physiological vessel formation that also occurs throughout the entire life span is intussusceptive angiogenesis, the process during which pre-existing vessels split and give rise to daughter vessels without a corresponding increase in the number of endothelial cells^1,7,8^. Besides these three modes of physiological blood vessel formation, which can also occur in pathological settings, three additional mechanisms modifying blood vessels have been described in pathological settings (mainly tumors)^4^. These include vascular co-option, describing the organization of tumor cells into perivascular cuffs around existing microvessels of the surrounding healthy tissue to allow the oxygenation of the early tumor mass^4,9,10^; vascular mimicry, defined as the integration of tumor cells into the blood vessel wall mimicking endothelial cells^11,12^; and glioma stem-to-endothelial or glioma stem-to-pericyte transdifferentiation^13–16^. The relative importance of these modes of vessel formation and modification in the formation of a well-established 3D vascular network depends on the developmental stage and the pathology in the respective organs^1,2,5^ and are still not completely understood^1^.

### Angiogenesis and vascular network formation in the postnatal and adult mouse brain

The human brain crucially depends on its vasculature for proper functioning, as it – despite accounting for only 2% of the body mass – consumes 20% of the cardiac output and 20% of the oxygen and glucose of the entire body, corresponding to a 10-fold higher rate of energy metabolism compared to an equivalent tissue volume in the rest of the body^17,18^. The brain vasculature is constituted of a complex 3D vascular network of 644 kilometers (400 miles)^19,20^, anatomically organized in a way that each individual neuron is not more than 15μm away from the closest microcapillary, thereby assuring proper oxygen delivery^21^. In the mouse brain, the absolute numbers are obviously lower but show similar relative values^22,23^. Cerebral angiogenesis in mice and humans (and in all mammals) is highly active during embryonic and postnatal brain development and almost quiescent in the adult healthy brain^2,3,24^, but is reactivated in vascular-dependent central nervous system (CNS) pathologies^6,25,26^. Thus, addressing how the brain vasculature network shapes from the developmental to the adult stage is crucial for a better understanding of underlying vascular dynamics also in pathological conditions^6,27^.

In the developing mouse brain, the functional, perfused vascular network of the CNS parenchyma is established following multiple distinct steps: it starts at around embryonic day 7.5 – 8.5 (E7.5 – E8.5) with the formation of the perineural vascular plexus (PNVP, responsible for the vascularization of the future meninges)^28–30^ via vasculogenesis in the ventral neural tube following a ventral-to-dorsal gradient^6,31–33^. In a subsequent step, at around E9.5, vascular endothelial sprouts emanating from the PNVP invade the CNS parenchyma in a caudal to cranial direction, vascularizing the CNS tissue via sprouting angiogenesis, thereby forming the intraneural vascular plexus (INVP)^3,6,27,31,34–37^. The INVP is expanded and refined via subsequent vessel sprouting, branching and anastomosis^6,31,36,37^ while these vascular sprouts migrate from the PNVP radially inwards towards the ventricles^3,27,36^. These important angiogenic processes of blood vessel formation and remodeling start during embryonic CNS development^31,36^ and continue at the postnatal stage of brain development^17,27,30,38–40^. The expansion and remodeling of the intricate 3D CNS vascular network ^38,39^ is required in order to adapt to local metabolic needs and neural activity^27,38,39,41^. In order to stabilize the newly formed blood vessels, perivascular cell types of the neurovascular unit (NVU) such as vascular smooth muscle cells, pericytes, neurons and astrocytes, are recruited to the vessel wall^3,25,37,42–44^, thereby establishing the mature, functional CNS vasculature^3,6,25,45^. The establishment of the CNS vascular network via sprouting angiogenesis requires a subset of specialized endothelial cell types: endothelial tip cells at the leading edge of the vascular sprout extend finger-like filopodia to sense environmental cues thereby guiding the blood vessel sprout towards gradients of pro-angiogenic growth factors such as vascular endothelial growth factor (VEGF), basic fibroblast growth factor (bFGF)^1–3,46,47^, platelet-derived growth factor (PDGF)^48^, Ephrins^49^ and Angiopoietins (Angs)^50^, and away from gradients of anti-angiogenic factors such as Thrombospondin^51,52^, Endostatin^53^, Angioarrestin^54^, Netrin-1^55^, Semaphorins 3A-B-E-F-G^56^ or Nogo-A^39,40^. Behind the leading tip cells, endothelial stalk cells elongate the growing vascular sprout as they proliferate and form the vascular lumen^1–3,57–59^. Neighboring sprouts then fuse and form vascular anastomosis resulting in perfusion of the newly formed vessels^1–3,58,60–62^. While the new blood vessel further matures and reaches a functional stage, its endothelial cells acquire a quiescent state, called endothelial phalanx cells^1,2,63^. Perfusion is a prerequisite for the stabilization and maintenance of a functional vascular network, acting most likely via flow-induced shear-stress related signaling pathways, such as the Sphingosine 1-Phosphate Receptor 1 (S1PR1) signaling^64–67^ and the VEFR-2^68^ and flow-induced dose-dependent expression of the transcription factor Krüppel-like factor 2 (KLF2)^69,70^, which regulates the expression of various antioxidative and anti-inflammatory mediators to stabilize the vessel wall. Accordingly, the absence of perfusion of newly formed vascular sprouts leads to vessel regression, fine-tuned mainly by non-canonical Wnt signaling^71–73^. Whereas sprouting angiogenesis and vascular remodeling remain highly dynamic throughout the first postnatal month until around postnatal day 25 (P25) – with a peak in endothelial cell proliferation at around postnatal day 10 (P10)^24^ – most of these redundant newly formed vessel sprouts undergo strategic pruning (most likely influenced by regional metabolic needs related to neural activity) and only a small fraction ends up forming mature perfused capillaries^24^. After the first month, the number of vessel segments remains largely constant and the vascular network stabilizes, followed by a progressively decreased turnover between formed and eliminated vessels during physiological aging^24^.

### Reactivation of developmentally active angiogenic signaling cascades in vascular-dependent CNS pathologies and the concept of the neurovascular link (NVL)

Cerebral angiogenesis is highly active during embryonic and postnatal brain development and almost quiescent in the adult healthy brain^2,3,24^. In a variety of angiogenesis-dependent CNS pathologies such as brain tumors, brain arteriovenous malformations, or stroke, developmental signaling programs driving angiogenic growth during development are reactivated^1,2,6^, thereby activating endothelial- and perivascular cells of the NVU^3,6,74^. However, which (developmental) molecular signaling cascades are reactivated in these angiogenesis-dependent CNS pathologies and how they regulate angiogenesis and vascular network formation on a cellular and subcellular level is not well understood. Thus, more knowledge is needed to better define the mechanisms of angiogenesis and vascular network formation during brain development, in the adult healthy brain as well as in brain pathologies.

Directly related to this reactivation of developmentally active signaling cascades is the concept of the neurovascular link (NVL), defining the cellular and molecular basis of the parallel organization and alignment of arteries and nerves in the brain, which is of crucial importance to understand brain vascular development, both at the embryonic and at the postnatal stage^6,25^. At the (sub)cellular level, the growth cones of newly formed axons resemble the endothelial tip cells of sprouting blood vessels^3^. Microenvironmental stimulating and inhibiting signals result in extension and retraction of these structures thereby inducing directed movements of growing nerves and blood vessels^6,25,43^. At the functional level, numerous common molecular cues that guide both endothelial tip cells and axonal growth cones via these specialized cellular and subcellular structures have been discovered^3,6,75^. As angiogenesis is highly active during postnatal brain development^38,39,76^, it provides an adequate model system to address the process of sprouting angiogenesis, which is the predominant mode of vessel formation in the brain^6,31^. Molecularly, sprouting angiogenesis and endothelial tip cells are regulated via numerous signaling axes. For instance, endothelial tip cells are – similar to axonal growth cones – guided by the interplay of attractive and repulsive cues such as the classical axonal guidance molecules Netrins, Semaphorins, Slits and Ephrins, as well as the neural growth inhibitor Nogo-A^3,4,6,25,39,40^. We have previously shown that in mice lacking functional Nogo-A, additional endothelial tip cells and vascular sprouts and can result in an increased number of perfused blood vessels that integrate into the vascular network^27,39^.

It is further known that the transition from sprouting endothelial tip cells to functional, perfused blood vessels that integrate into vascular networks is mainly regulated by the VEGF-VEGFR– delta-like ligand 4 (Dll4)-Jagged1-Notch pathway^1,2,62,77–82^, thought to be a central pattern generator of vascular sprout- and endothelial tip cell formation. The spatiotemporally tightly regulated balance between endothelial tip- and stalk cell numbers and the balance between tip migration and stalk proliferation affect branching frequency and plexus density^78^. It was also shown that disturbance of these molecular processes may result in non-productive angiogenesis with the formation of aberrant blood vessels^80–84^. Importantly, the remodeling of the final perfused vascular network exerting its crucial functions depends on many different factors that include tissue-specific metabolic and mechanical feedbacks from the microenvironment as well as blood-flow related signaling pathways, respectively^73^. Nowadays, these processes of vascular remodeling from the developing to the adult, mature stage are incompletely understood^62^.

For instance, most of the numerously generated vascular sprouts in the first postnatal, highly angiogenic month in the mouse brain, undergo pruning and only a small fraction results in functional, perfused capillaries^24^. Thereafter the brain vascular network remains relatively stable. Nevertheless, despite the vascular stability at the adult stage, some vessel formation and elimination continue throughout life^24^.

Another interesting aspect is that while we observed a significantly increased vessel density in young postnatal mice lacking function Nogo-A, this effect was almost completely gone in the adult Nogo-A^−/−^ mice^39^, raising questions regarding compensatory mechanisms by other molecules in the Nogo-A^−/−^ mice but also regarding the plasticity of the vasculature in the young postnatal versus the adult stage.

These findings underline the need for a method allowing to address and quantify differences of 3D vascular networks obtained from mice at any age between the postnatal developing-and the adult stage. At present, to the best of our knowledge, there is no established protocol to quantitatively chart the 3D architecture of the brain vascular network of mice at any postnatal age. However, such a protocol is crucial in order to investigate the mechanical and metabolic feedback that is finally responsible for maturation and stability of functional, perfused, 3D vascular networks^73^. The protocol presented here fills this gap and allows direct and quantitative comparison of the brain vascular network tree between different postnatal and / or adult developmental stages as well as to distinguish capillaries and non-capillary blood vessels based on various parameters such as vascular volume fraction, vascular branch point density and - degree, vessel segment length, -diameter, -volume, -tortuosity, -directionality, and extravascular distance.

### Development of the protocol

Vascular network formation has previously been addressed using vascular corrosion casting techniques in mice at the embryonic, postnatal and adult stages of development. In mice at the embryonic stage (8.5-13 days of gestation), vascular corrosion casting has been applied to investigate embryonic aortic development^85–87^ and the fetal circulation in relation to the placenta^88,89^. In adult mice, vascular corrosion casting techniques are widely used in all major organs, shown for example to be particularly useful to study cochlear vasculature in age-related hearing loss^90–93^, the microvasculature in the urine bladder^94^, and to investigate alveolar anatomy during both physiological development^95,96^ and in several diseases^97,98^.

Regarding the brain vasculature, vascular corrosion casting is a well-established technique to study cerebrovascular anatomy in physiological settings in mice^40,99–107^. Angiogenesis and vascular network morphology have previously been described using vascular corrosion casting in the adult healthy mouse brain^27,39^, for instance using brain-specific VEGF_165_-overexpressing mice (age: 16±2 weeks = P112) to study the effects of this pro-angiogenic stimulus on the adult brain vascular network^108^. Moreover, various vascular-dependent CNS pathologies have been investigated using the technique, examples are mouse models of Alzheimer’s disease using APP23 transgenic mice (age: ranging from 3 to 27 months of age = P90 to P810)^109,110^, brain arteriovenous malformations utilizing an *Alk-1*^2f/2f^ (exons 4-6 flanked by loxP-sites) VEGF stimulated mouse models^111,112^, microvascular networks in brain tumors in rats and primates using gliosarcoma xenotransplantation^113^, a post-traumatic brain injury model in rats^114^ after transient focal cerebral ischemia in rats^115^ and postmortem in human traumatic brain injury patients^116^.

Recently, we have adapted the vascular corrosion casting technique enabling to describe the effects of the well-known axonal growth-inhibitory protein Nogo-A – which we have previously identified as a negative regulator of CNS angiogenesis and endothelial tip cell filopodia^39^ – on the 3D vascular network in the postnatal mouse brain^40^.

Here, we demonstrate an optimized adaptation and extension of these previously described methods, allowing hierarchical imaging and computational analysis of the vascular network tree during any postnatal brain developmental stage in the mouse brain. Using vascular corrosion casting, hierarchical, desktop- and SRµCT-based imaging and image analysis we provide a refined, quantitative 3D analysis generating absolute parameters, characterizing the entire 3D mouse brain vessel network at a µm-resolution.

### Applications of the protocol

The presented protocol comprises vascular corrosion casting, hierarchical imaging and computational analysis of individual vessel segments of the 3D vascular network in postnatal and adult mouse brains. The method distinguishes between vessels of different diameters, for instance between capillaries (inner diameter of < 7 µm) and non-capillaries (inner diameter ≥ 7 µm). The protocol allows to study the influence of any angiogenic molecule including pharmacological compounds and blocking antibodies, and to test their pro- or antiangiogenic effects on vascularization and various specific parameters of the vascular network in postnatal as well as adult wild-type or genetically modified mice^39,117^. Importantly, the method provides a detailed quantitative description of the 3D vasculature allowing precise comparison of different experimental settings or genetic modifications. Quantitative information gained include vascular volume fraction, vascular branch point density and -degree, vessel segment length, -diameter, -volume, -tortuosity, -directionality, and extravascular distance, on both the level of global vascular network morphometry as well as local vascular network topology. This is quite different from the classical use of the corrosion cast technique for which in general only gross visual comparisons have been made and described. Importantly, the method could be used with laboratory/desktop μCT scanners only, which significantly increases the applicability of the protocol as compared to combined desktop μCT plus SRμCT setups. We have advanced the method which now can also be applied to the analysis of the 3D vascular network in the entire mouse brain by creating artefact-free, high-resolution images of the complete and intact vascular corrosion cast, allowing to directly address the vascular network morphology of any given region of interest within the mouse brain. This improves precision and is a significant simplification in contrast to the two-step ROI-based approach during which the ROIs first have to be placed on a lower resolution overview μCT scan with subsequent high-resolution SRμCT or desktop μCT scanning of selected ROIs. Finally, given the crucial role of angiogenesis and vascular networks during both tissue development as well as in various pathologies in- and outside the CNS^1,2,6,118,119^, and referring to previous vascular corrosion casting-based analyses of Alzheimer’s disease, brain tumors, and other pathologies^110,111,113,114^, the improved method presented here can – after validation in other tissues or pathological conditions – in principle be applied to virtually any physiological or pathological setting.

### Experimental design

Here, we present a technique for the visualization and computational analysis of the 3D vascular network in the postnatal and adult mouse brain. Figure 1 illustrates the different steps of the procedure with the corresponding timescale. Briefly, the protocol starts with transcardial perfusion of artificial cerebrospinal fluid (ACSF) containing Heparin, followed by 4% paraformaldehyde in phosphate-buffered saline (PBS) and the resin PU4ii (vasQtec, Zürich, Switzerland) at the desired day of postnatal development (in our case P10, corresponding to the peak of endothelial cell proliferation in the mouse brain according to Harb et al., *J Cereb Blood Flow Metab, 2013*^24^) or adulthood (in our case P60). This is followed by resin curing (overnight), soft tissue maceration (for 24h), decalcification of the surrounding bone structures (overnight), dissection of the cerebral vascular corrosion casts from the remaining extracranial vessels, followed by subsequent washing, freeze-drying (lyophilization), contrasting (osmium tetroxide), optional low-resolution desktop-µCT scanning for region-of-interest selection, high-resolution SR-µCT or desktop-µCT scanning, and computational analysis.

**Figure 1.**
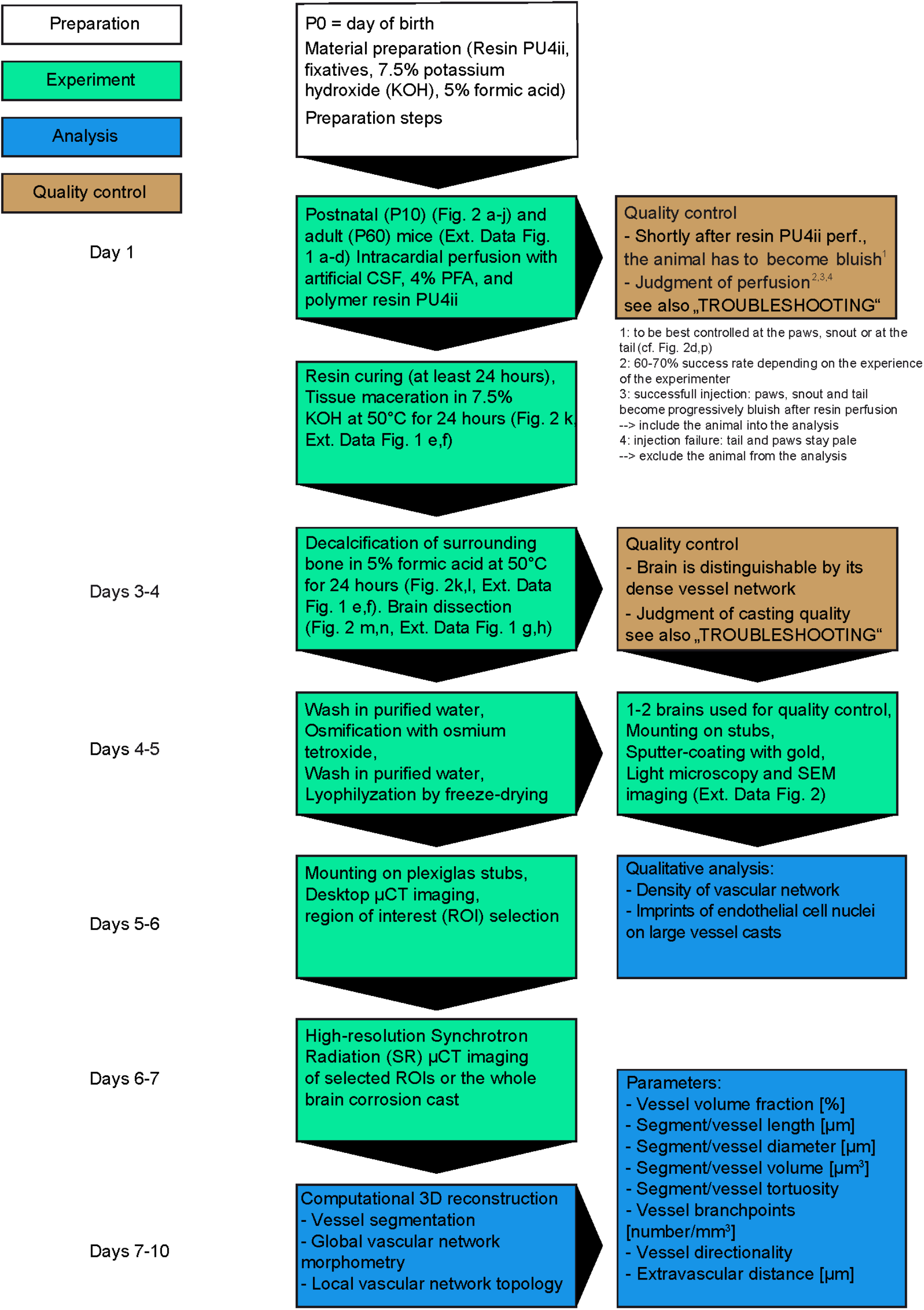
Flow chart, summary, and time frame of the protocol. The different sections of the protocol are indicated in color-coded boxes. White boxes indicate preparation steps, green boxes describe experimental steps, blue boxes indicate analysis steps and brown boxes describe quality control steps. Black arrows indicate the order in which the different steps of the protocol are performed. A time scale in days is shown on the left.

In more detail, after the animal is properly anesthetized, the chest is opened in order to gain access to the left ventricle (Figure 2a-f, Extended Data Figure 1a,b). To this end, the skin is lifted with the forceps and then cut perpendicular to expose the sternum (Figure 2c,d,g,h, and Extended Data Figure 1b). The sternum is then lifted and lateral cuts through the rib cage are made exposing the chest cavity. The rib cage flap is pulled back and fixed using a pin/needle and the pericardium is carefully removed using rounded forceps. The heart is lifted slightly using forceps and then a winged perfusion needle is inserted into the left ventricle through the apex and the point of insertion is sealed with instant glue (Figure 2e,i, and Extended Data Figure 1c). Note the insertion at a flat angle of about 30° (Figure 2e,i). Care is taken during this procedure not to insert the needle too deep into the heart in order to avoid damage of the atrioventricular valve. After proper placing and securing of the needle (using the wings of the needle and pins/needles) in the left ventricle, the right atrium is opened to allow the blood and perfused fluid to leave the circulation (Figure 2b,f,j, Extended Data Figure 1d, and Supplementary Video 1). Perfusion of 5 ml (for P10 animals) and 20 ml (for adult animals) of artificial cerebrospinal fluid (ACSF) containing 25,000 U/L Heparin through the left ventricle, is followed by perfusion with 4% paraformaldehyde (solved in PBS) and the resin PU4ii solution. All perfusion solutions are perfused at a pressure of 90-100 mmHg for P10 mice (4 ml/min) and 100-120 mmHg for P60 mice. Successful perfusion is indicated by sufficient outflow of the solutions from the opening in the right atrium and by the bluish appearance of the paws, snout and tail immediately after starting the resin perfusion (Figure 2f,j, Extended Data Figure 1d and Supplementary Video 1). After 24 hours of resin curing, soft tissue is macerated in 300-400ml 7.5% potassium hydroxide (KOH) for at least 24 hours, followed by incubation in 300-400 ml 5% formic acid for another 24 hours to decalcify bony structures. The brain vascular corrosion cast is dissected out of the skull and freed from extracranial vessels (Figure 2k-n, Extended Data Figure 1e-h, and Supplementary Video 1).

**Figure 2.**
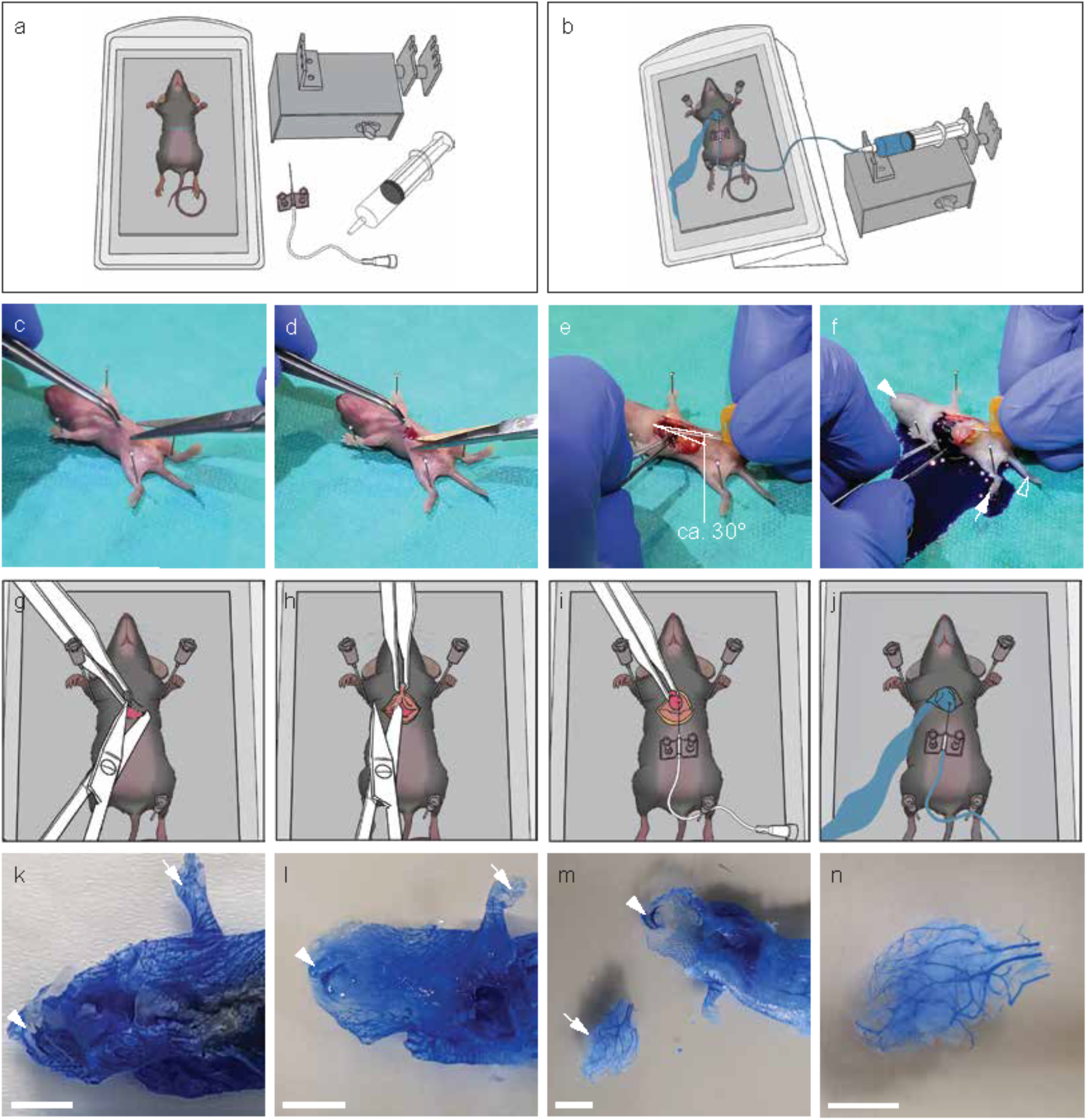
Intracardial resin perfusion and brain dissection for postnatal day 10 (P10) animals. **a-b** Schematic drawing illustrating the experimental setup before (**a)** and after **(b)** resin perfusion. Surgical opening (**c,d**), angle of 30 degrees in the sagittal plane (**e**) and site of needle insertion (**e**) for intracardial perfusion with 10–20 ml artificial cerebrospinal fluid containing 25,000 U/L Heparin via the left ventricle, followed by 4% paraformaldehyde in phosphate buffered saline, and by **(f)** the polymer resin PU4ii (vasQtec, Zürich, Switzerland), were all infused at the same rate (4 ml/min and 90-100 mmHg) in a P10 mouse pup (see also Supplementary Video 1). Successful perfusion is indicated by a bluish aspect of the animal, which can be well observed at the paws (white arrow), snout (white arrowhead) or tail (white outlined arrowhead). **g-j** Schematic drawings depicting the individual steps visible in **c-f**. **k** Resin cast of a P10 animal after soft tissue maceration, before (**k**) and after (**l**) bony tissue decalcification (paw, white arrow and snout, white arrowhead). **m** The cerebral vasculature is sharply dissected from the extracranial vessels resulting in the isolated brain vascular corrosion cast of the P10 mouse brain (white arrow, **m**) (**n**). Scale bars, 5 mm (**k-n**).

The principle of vascular corrosion casting is schematically shown in Figure 3. The step-by-step procedure of intracardial polymer resin perfusion, tissue maceration, bone decalcification, and brain dissection, is illustrated in Figure 2, in Extended Data Figure 1 and in Supplementary Video 1. As an additional sidestep, not part of the main protocol pipeline, a combination of resin perfusion and immunofluorescence staining and imaging was performed (Extended Data Figure 10). For this purpose, resin curing is immediately followed by removal of bony structures (without tissue maceration), in order to preserve soft tissue required for immunofluorescence staining. After cutting, the soft tissue slices are then incubated with biological markers of interest such as Fluoro Myelin Red (Molecular Probe Inc) for myelin and DAPI. Additional to the inherent fluorescence of PU4ii, specific fluorescent dyes can be used (Extended Data Figure 10).

**Figure 3.**
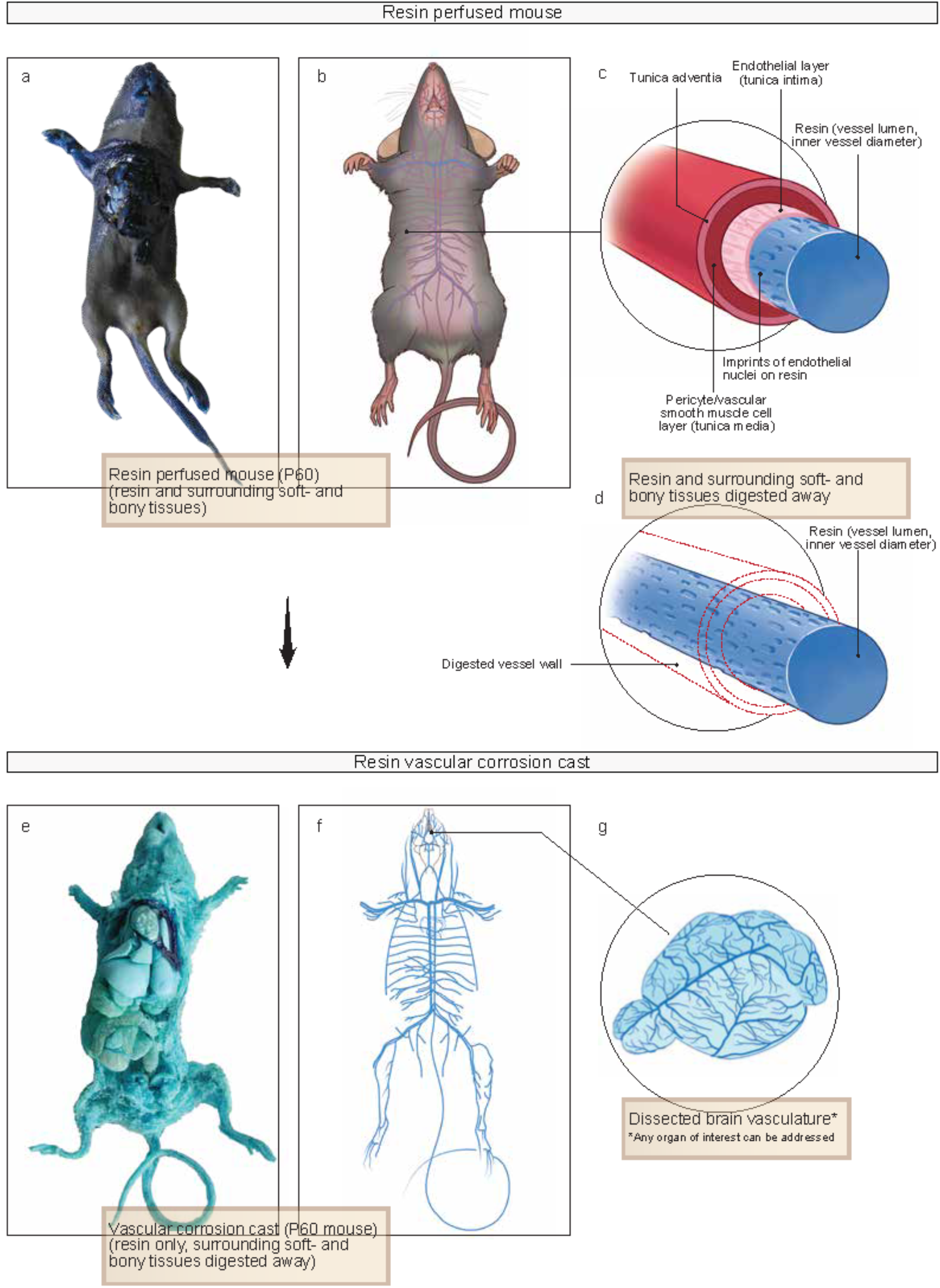
The principles of (brain) vascular corrosion casting. **a-g** Photograph **(a)** and schematic drawing (**b**) of a whole adult (P60) mouse after perfusion of resin PU4ii. Insets showing a schematic illustration of the principle of vascular corrosion casting (**c,d**), in which the vascular corrosion casts represents the inner volume of perfused blood vessels in blue; defined by the inner vessel diameter, before **(c)** and after **(d)** maceration of soft tissues. Maceration of the surrounding CNS tissue including the blood vessel wall and decalcification of bony structures, exposes the entire vascular corrosion cast (**e,f**), with a schematic drawing of the casted brain vasculature in more detail **(g)**.

For quality control, random samples of vascular corrosion casts are mounted on stubs and sputter-coated with gold^109,120^ for scanning electron microscopy (SEM, Hitachi S4000) (Extended Data Figure 2). Optionally, the quality control could be performed with scanning helium ion microscopy that does not require sputter coating. In this case, the same casts that will be μCT imaged can be used for quality control.

The casts that are to be μCT imaged are stained using osmium tetroxide in order to reach better X-ray absorption leading to higher signal to noise ratio (SNR) and, thus, better contrast in X-ray imaging^108^. Afterwards, the osmicated casts are lyophilized overnight and mounted on Plexiglas stubs or other suitable sample holders using cyanoacrylate glue. Optionally, the entire corrosion casts are first imaged with low-resolution desktop-µCT to select desired ROIs. This step is not required if the whole casts are to be imaged.

The corrosion casts are scanned with high-resolution desktop-, or SR-μCT (either preselected ROIs or the whole brain) and analyzed using a computational imaging processing pipeline. The image analysis procedure allows segmentation of the 3D data such that voxels representing vessels are assigned value 1 and all other voxels are assigned value 0. Furthermore, this binary representation is converted into a vessel graph (Figure 4) that facilitates quantitative analysis of all network parameters, optionally separated for capillaries and non-capillaries, including vascular volume fraction (Figures 5,6, Extended Data Figures 3,4, and Supplementary Figures 1,2), branch point density and branch point degree (Figure 7, Extended Data Figures 5–9, and Supplementary Figure 3), segment density, -diameter, -length, -volume and -tortuosity (Figure 8, and Supplementary Figure 4), extravascular distance (Figure 9, and Supplementary Figure 5), and vessel directionality (Figure 10, and Supplementary Figures 6–9). Finally, the data extracted using image analysis procedures are evaluated statistically.

**Figure 4.**
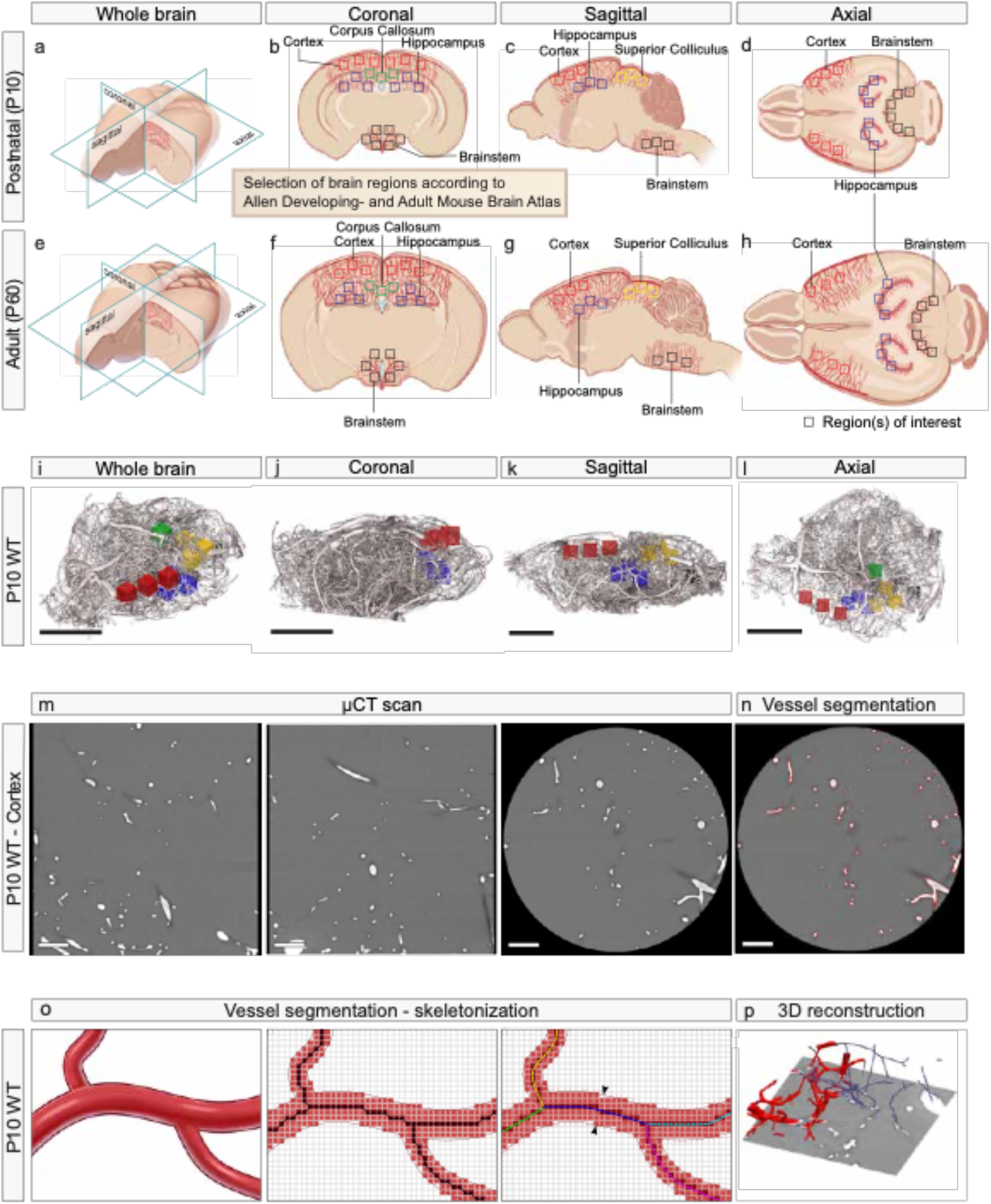
The placement of the regions of interest (ROIs) and process of computational 3D reconstruction. Schematic visualization of the ROI placement in both postnatal (P10) (**a-d**) and adult (P60) (**e-h**) mouse brains with 3D overviews (**a,e**) and detailed panels for the respective anatomical plane (**b,f** coronal, **c,g** sagittal, **d,h** axial). The µCT scanning procedure of a vascular corrosion cast, includes the placement of ROIs, here depicted in the 3D rendering of a whole postnatal (P10) WT brain, shown in the different anatomical planes **(i-l)** (red: cortex, blue: hippocampus, yellow: superior colliculus, green: corpus callosum, and black: brain stem). **m** Raw images of single µCT projections in the x-plane, y-plane, and in the z-plane (from left to right) of a ROI placed in the cortex of a P10 WT animal, followed by **(n)** the same z-plane for which the edges of the segmented regions are shown as red outlines (Otsu’s thresholding method). **o** Visualization of the image segmentation and skeleton tracing process. In the imaging process, the original vessel structure (left) is converted into a digital approximation that is segmented using the Otsu method (middle). The centerlines of the vessels are found using skeletonization (middle, highlighted pixels). The pixel-accurate centerline (middle, black dotted line) is smoothed for better representation of the centerlines of the individual vessel branches (right, colored lines). For each branch its length is measured as a sum of distances between the smoothed centerline points (right, colored points), and its diameter as the average diameter of the branch (right, arrows). **p** Three-dimensional visualization of the original data (P10 WT cortex) showing the CT plane (grayscale), the segmented vessels (red, left side) and the vessel centerlines (blue lines, right side). Part of segmentation has been cut off in the front of the CT plane. Scale bars: 250 µm (**i-l**),100 µm (**m,n**)

**Figure 5.**
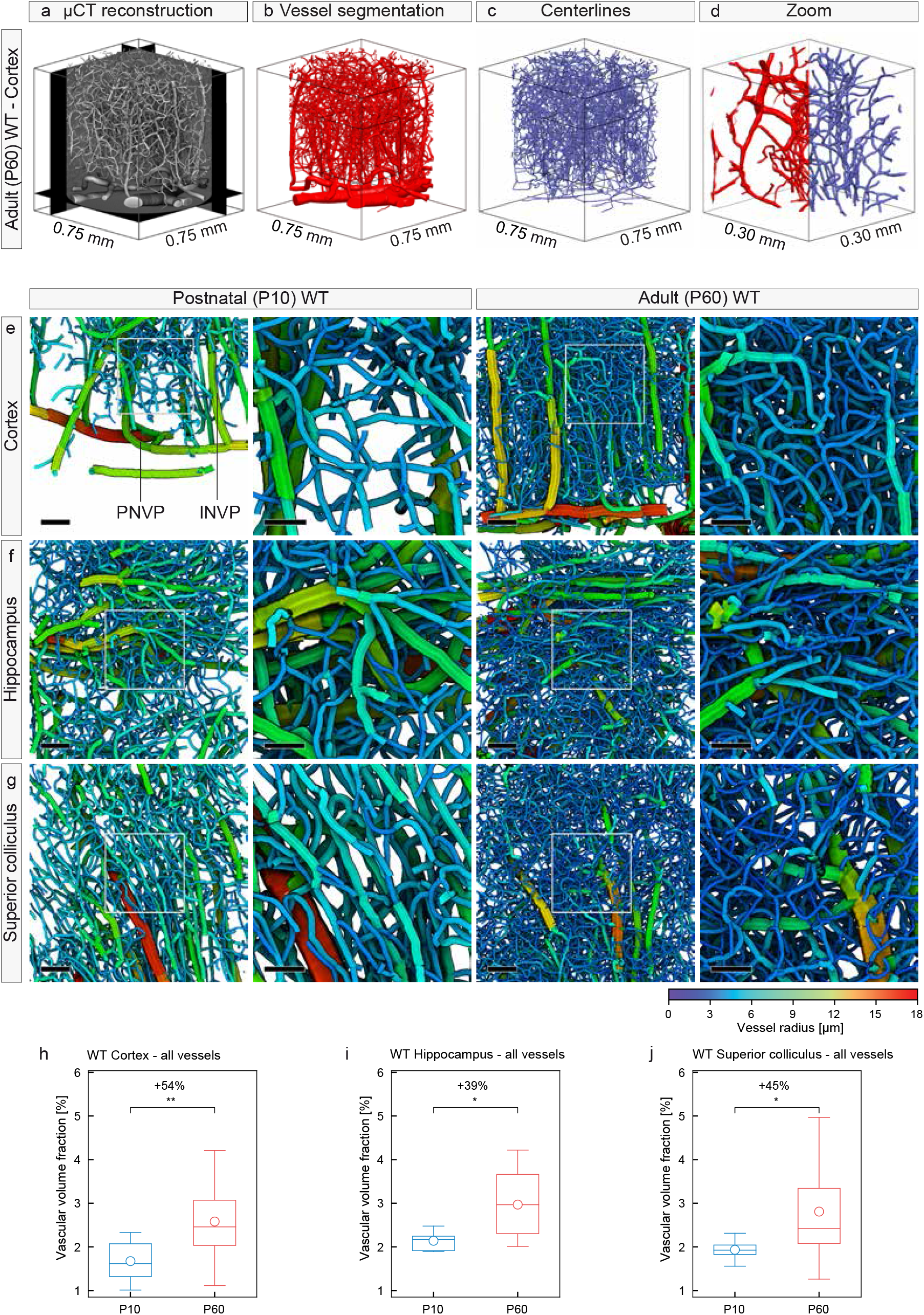
Global vascular network morphometry - Increased vascular volume fraction in various regions of the adult (P60) WT versus postnatal (P10) WT mouse brain. **a-d** Computational 3D reconstructions of the vascular network using surface rendering based on the synchrotron radiation µCT data showing a ROI as volume mask (**a**), after the vessel segmentation process (**b**), and after the placement of centerlines (**c**), with a zoom of both stages combined (**d**). Blood vessel structures of all sizes are well defined. **e-g** Computational 3D reconstructions of µCT scans of vascular networks of the postnatal (P10) WT and adult (P60) WT mouse brains displayed with color-coded vessel thickness. The increased vessel density in the P60 WT cortices (**e**), hippocampi (**f**) and superior colliculi (**g**) is obvious. Color bar indicates vessel radius (µm). The boxed areas are enlarged at right. **h-j** Quantification of the 3D vascular volume fraction in P10 cortex (**h**), hippocampus (**i**), and superior colliculus (**j**), by computational analysis using a global morphometry approach. The vascular volume fraction in all these brain structures was significantly higher in the P60 WT animals than that in the P10 WT animals (n = 4-6 for P10 WT; n = 8-12 for P60 WT animals; and in average three ROIs per animal and brain region have been used). All data are shown as mean distributions, where the open dot represents the mean. Boxplots indicate the 25% to 75% quartiles of the data. *P < 0.05, **P < 0.01, ***P < 0.001. Scale bars: 100 µm (**e-g**, overview); 50 µm (**e-g**, zoom).

**Figure 6.**
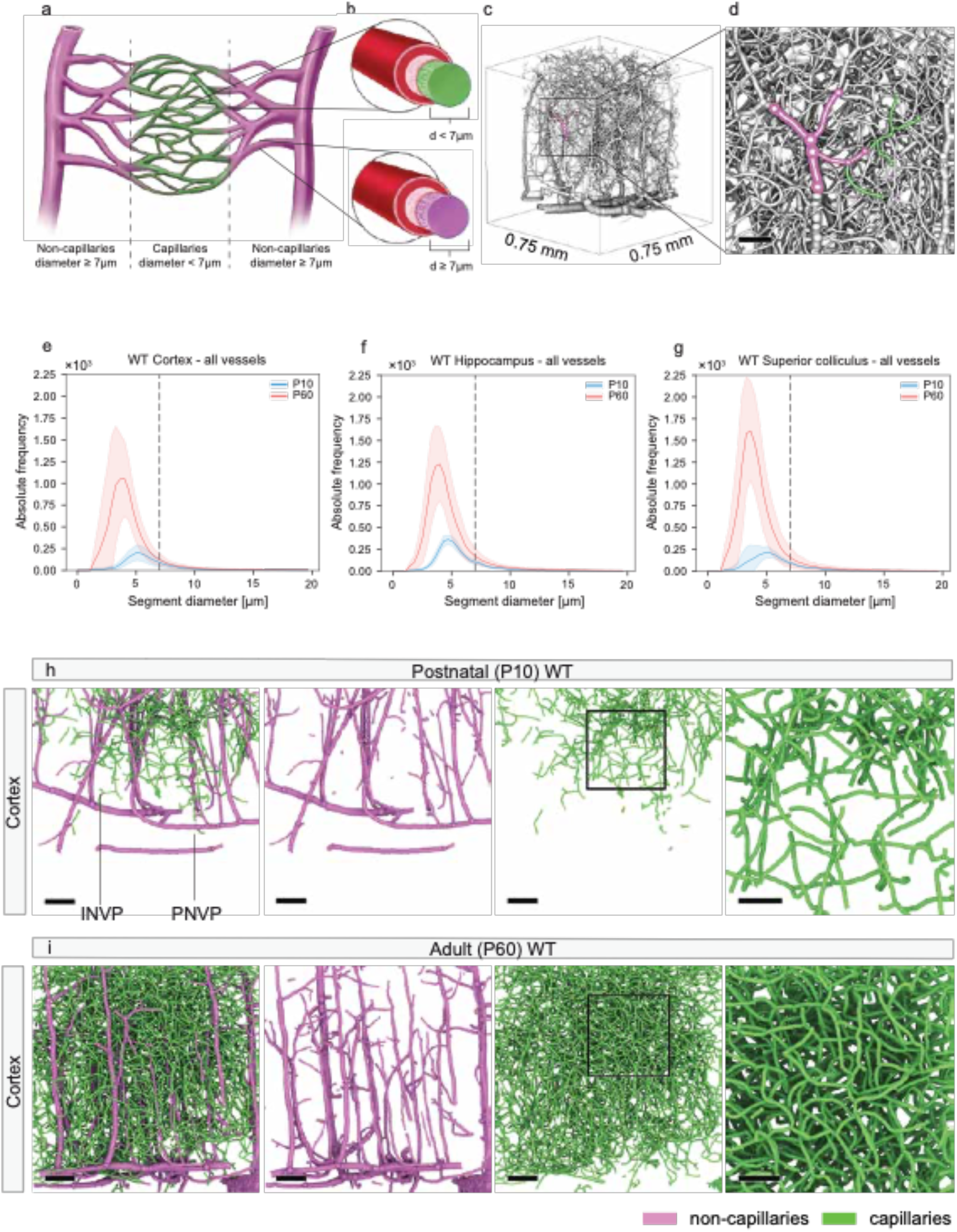
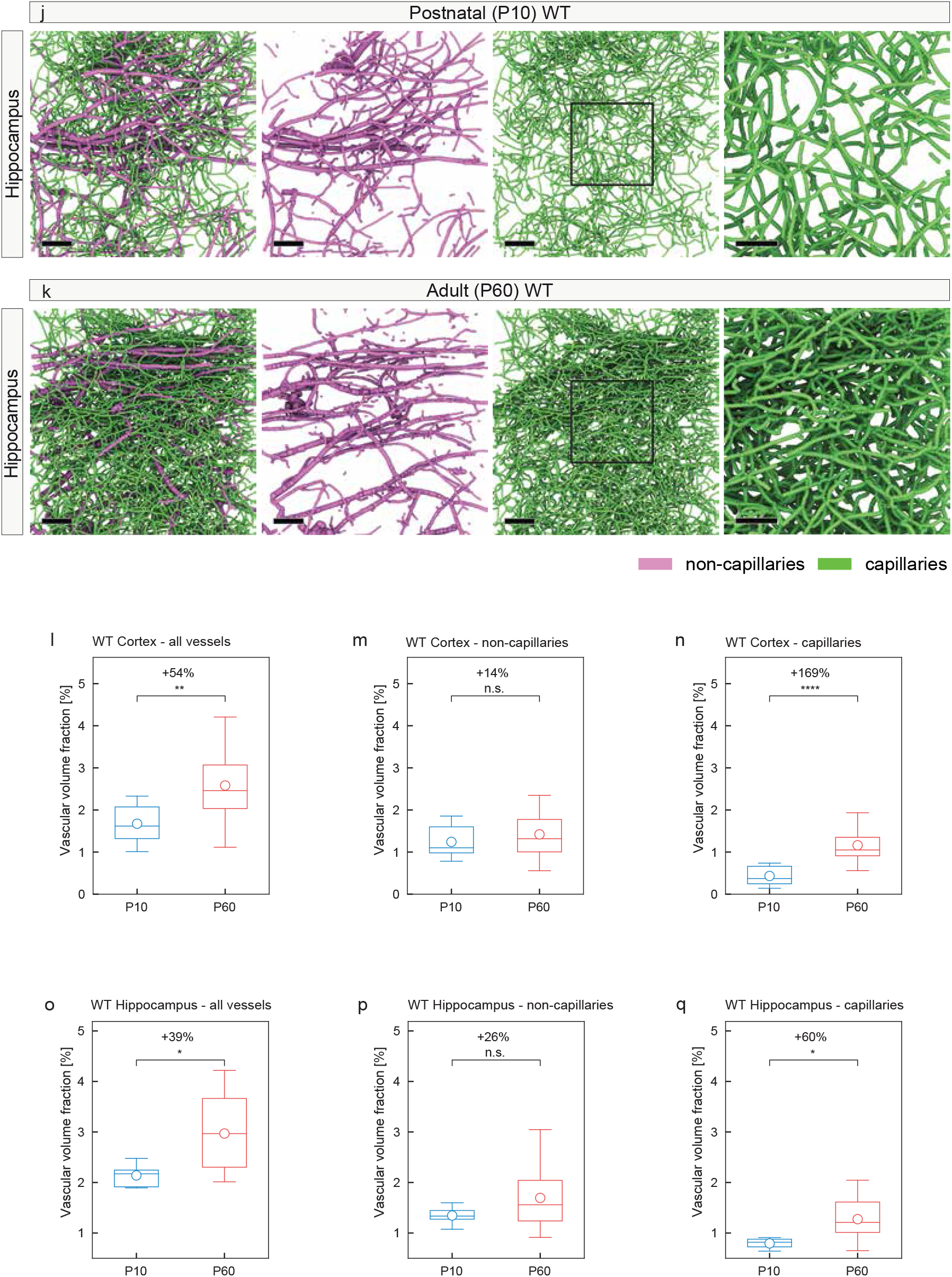
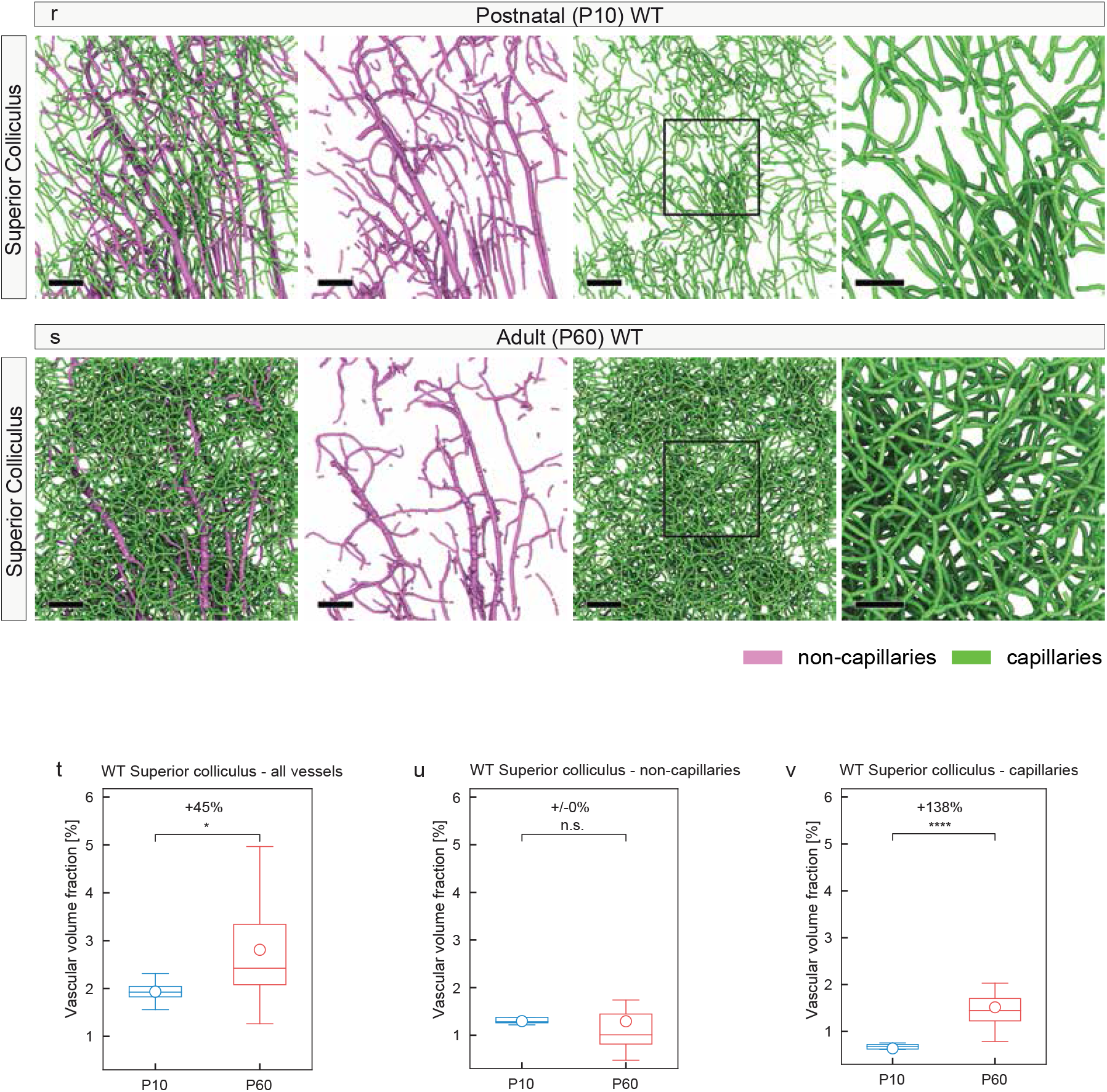
Local vascular network topology – Increased vascular volume fraction of the adult (P60) WT versus postnatal (P10) WT mouse brain mainly found at the capillary level. **a,b** Scheme illustrating a capillary bed and the distinction between capillaries and non-capillaries based on the radius of the inner volume of resin-perfused blood vessels after maceration of the surrounding CNS tissue including the blood vessel wall. A capillary is defined as a blood vessel with an inner diameter (**a,b**, green) of < 7µm, whereas a non-capillary vessel is defined a blood vessel with an inner diameter (**a**,**b**, magenta) of ≥ 7µm, according to^173^. **c,d** Three-dimensional computational reconstruction of µCT scans of cortical vascular networks in a P60 WT sample highlighting a vessel tree with non-capillaries in magenta and capillaries in green. **e-g** Histograms showing the distribution of vessel diameter in P10 WT and P60 WT animals. P60 WT animals show an increased absolute frequency of capillaries as compared to the P10 WT situation, whereas the number of non-capillaries remains largely unchanged in (**e**) the cortex, (**f**) hippocampus, (**g**) and superior colliculus (bin width = 0.36 µm; number of bins = 55). Black dashed line marks the separation between capillaries (< 7µm) and non-capillaries (≥ 7µm). **h-k,r,s** Computational 3D reconstructions of µCT images of vascular networks of P10 WT and P60 WT mice in the cortices (**h,i**), hippocampi (**j,k**), and superior colliculi (**r,s**) separated for non-capillaries (magenta) and capillaries (green). The increased density of capillaries (green) in the P60 WT samples for all three brain regions is evident, density in non-capillaries (magenta) is also increased. The boxed areas are enlarged at the right for capillaries (**h-k,r,s**). **l-q,t-v** Quantification of the 3D vascular volume fraction for all vessels (**l,o,t**), non-capillaries (**m,p,u**), and capillaries (**n,q,v**) in P10 WT and P60 WT calculated by local topology analysis. The significant increase of the vascular volume fraction for all vessels in the P60 WT animals (**l,o,t**) was mainly due to a significant increase at the level of capillaries (**n,q,v**) (n = 4-6 for P10 WT; n = 8-12 for P60 WT animals were used; and in average three ROIs per animal and brain region were used). All data are shown as mean distributions where the open dot represents the mean. Boxplots indicate the 25% to 75% quartiles of the data. The shaded blue and red areas indicate the SD. *P < 0.05, **P < 0.01, ***P < 0.001. Scale bars: 100 µm (**h-k**, overview); 50 µm (**h-k**, zoom).

**Figure 7.**
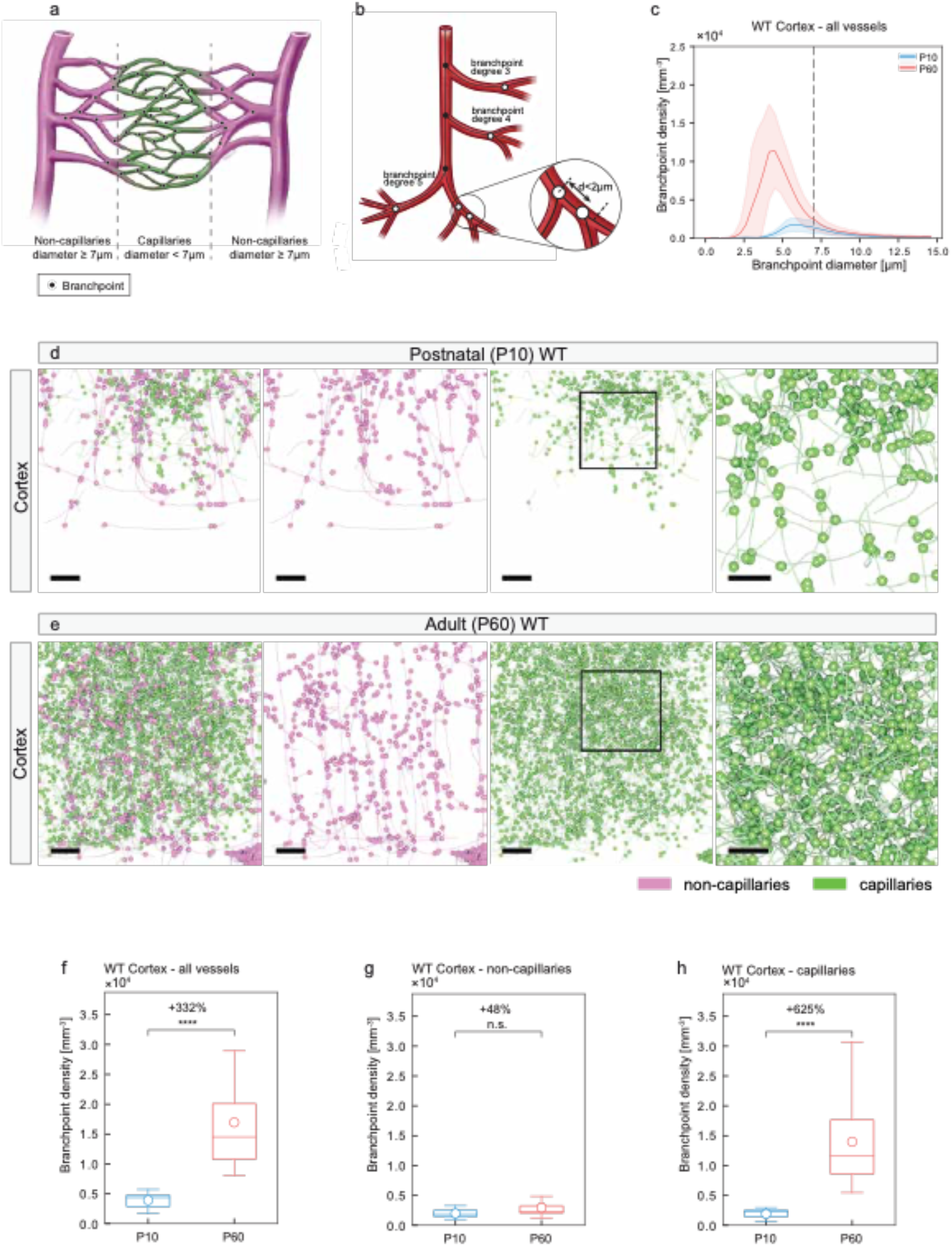
Local vascular network topology – visualization and quantification of vascular branch point diameter and vascular branch point density in various regions of the adult (P60) WT versus the postnatal (P10) WT mouse brain. **a** Scheme illustrating a capillary bed and the distinction between capillaries and non-capillaries based on the radius of the inner volume of resin-perfused blood vessels (see legend figure 6a). **b** Scheme depicting the definition of vascular branch points. Each voxel of the vessel center line (black) with more than two neighboring voxels was defined as a vascular branch point. This results in branch point degrees (number of vessels joining in a certain branch point) of minimally three. In addition, two branch points were considered as a single one if the distance between them was below 2µm (**b**). Branch point are depicted as black dots in **a,b**. **c** Histogram showing the distribution of branch point diameter in P10 WT and P60 WT animals. P60 WT animals show an increased branch point density as compared to P10 WT mice mainly at the capillary level (bin width = 0.38µm; number of bins = 40). Black dashed line marks the separation between capillaries (< 7µm) and non-capillaries (≥ 7µm), as defined in Figure 6. **d,e** Computational 3D reconstructions of µCT images of cortical vascular networks of P10 WT and P60 WT mice with visualizations of the vessel branch points displayed as dots, separately for non-capillaries (magenta) and capillaries (green). The higher density of branch points in the P60 WT cortices especially at the capillary level (green) is obvious, a slight increase of branch point density can be observed at the non-capillary level (magenta). The boxed areas are enlarged at the right. **f-h** Quantitative analysis of the branch point density for all vessels (**f**), non-capillaries (**g**), and capillaries (**h**) in P10 WT and P60 WT cortices by local morphometry analysis. The significant increase of the branch point density for all vessels in the P60 WT animals (**f**) was mainly due to a significant increase at the level of capillaries (**h**) and in part due to a significant increase at the level of non-capillaries (**g**). (n = 4-6 for P10 WT; n = 8-12 for P60 WT animals were used; and in average three ROIs per animal were used). All data are shown as mean distributions where the open dot represents the mean. Boxplots indicate the 25% to 75% quartiles of the data. The shaded blue and red areas indicate the SD. *P<0.05, **P<0.01, ***P<0.001. Scale bars: 100 µm (**d,e**, overviews); 50 µm (**d,e**, zooms).

**Figure 8.**
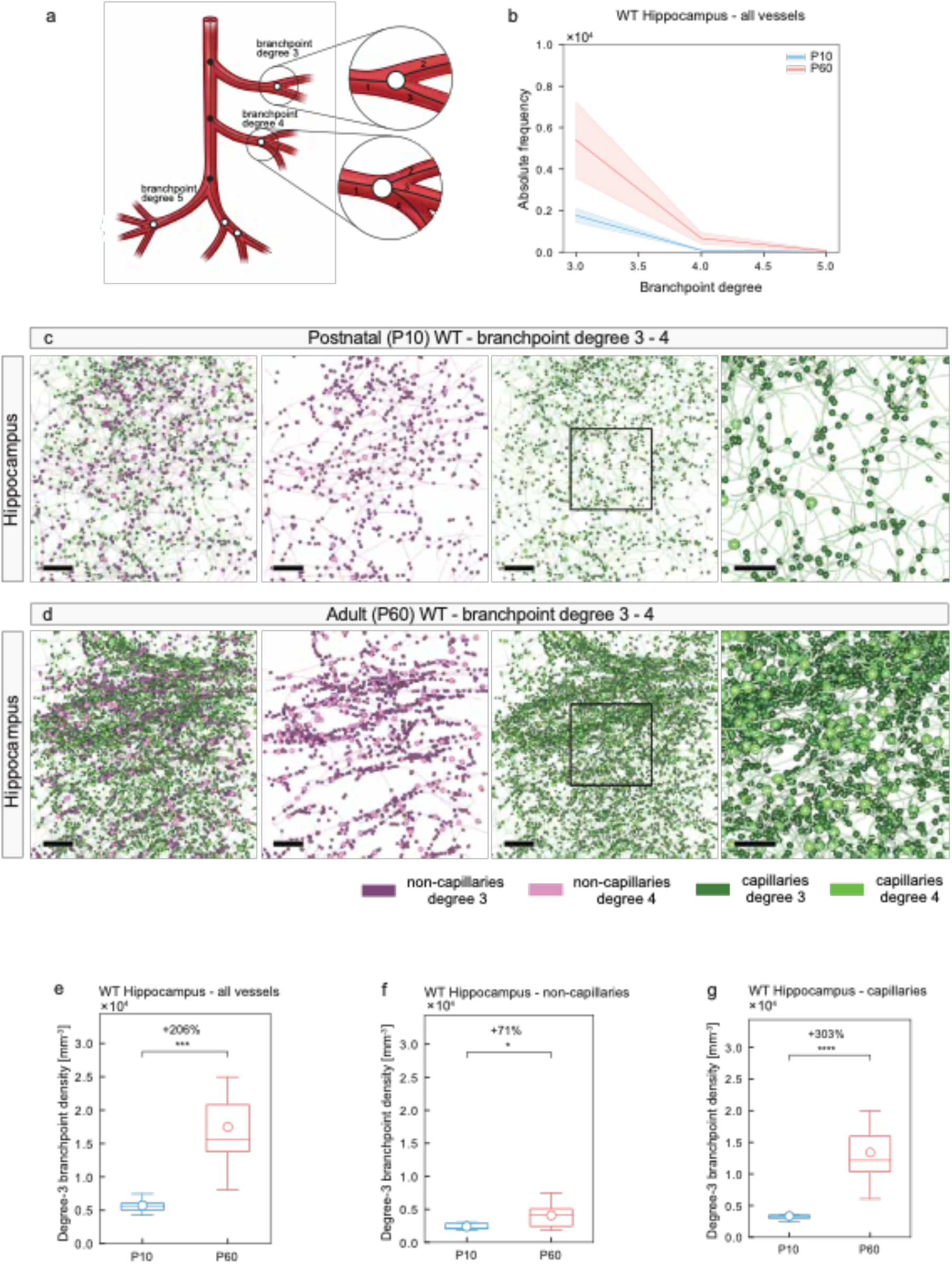
Local vascular network topology – segment density, -diameter, -length, - tortuosity, and -volume of the adult (P60) WT versus the postnatal (P10) WT mouse brain. **a,b** Schemes showing the definition of vessel diameter (**a**), vessel length (**a**), and vessel tortuosity (**b**). The segment diameter is defined as the average diameter of all single elements of a segment (**a**). The segment length is defined as the sum of the length of all single elements between two branch points (**a**). The segment tortuosity is the ratio between the effective distance l_e_ and the shortest distance l_s_ between the two branch points associated to this segment (**b**). The segment density is defined as the number of segments, classified into global, capillary and non-capillary segments, divided by the network volume. **c** Graph depicting the relationship between segment diameter and segment length in P10 WT and P60 WT animals. The ratio “segment diameter-to-segment length” was higher in P10 WT mice compared to P60 WT mice (bin width = 4.93; number of bins = 14). **d-f,g-s** Quantification of the 3D vessel network parameters segment density (**d-f**), segment diameter (**g-j**), segment length (**k-n**), and segment tortuosity (**p-s**) for all vessels, non-capillaries, and capillaries in P10 WT and P60 WT cortices calculated by local morphometry analysis. The segment density (**d**) and segment tortuosity (**q**) were significantly increased for all vessels in the cortices of P60 WT mice as compared to the P10 WT mice, whereas the segment diameter (**h**) and segment length (**l**) were significantly decreased for all vessels in the P60 WT mice as compared to the P10 WT mice. In line with the other parameters, these differences were mainly due to a highly significant increase respectively decrease at the level of capillaries (**f,j,n,s**) and in part due to an increase respectively decrease at the level of non-capillaries (**e,i,m,r**) (n = 4-6 for P10 WT; n = 8-12 for P60 WT animals; and in average three ROIs per animal and brain region were used). (**g** bin width = 0.36 µm; number of bins = 55, **k** bin width = 2.5 µm; number of bins = 40, **o** bin width = 0.013; number of bins = 80, **s** bin width = 2957.74 µm^3^; number of bins = 55). All data are shown as mean distributions where the open dot represents the mean. Boxplots indicate the 25% to 75% quartiles of the data. The shaded blue and red areas indicate the SD. *P < 0.05, **P < 0.01, ***P < 0.001.

**Figure 9.**
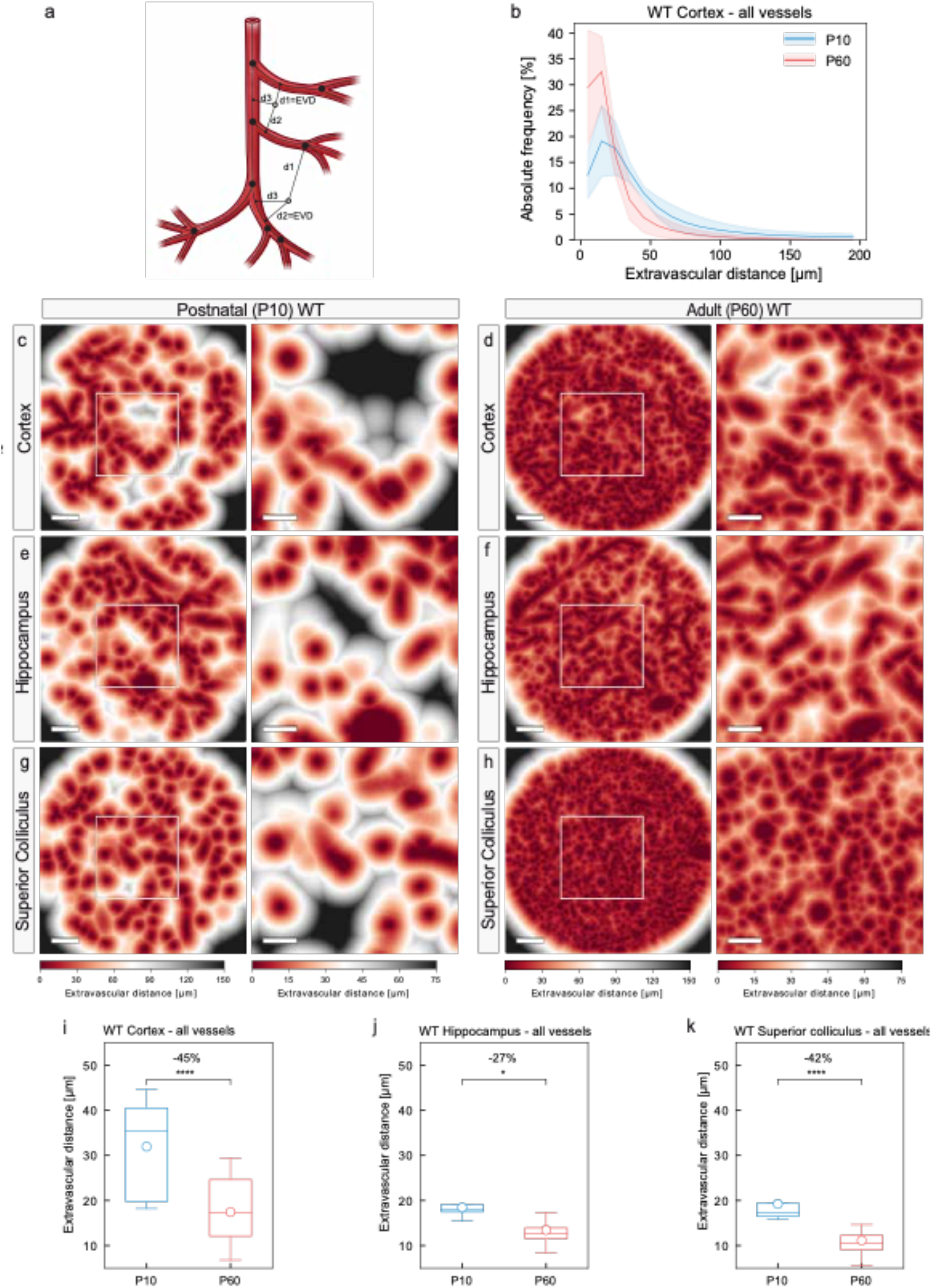
Local vascular network topology – extravascular distance is decreased in various regions of the adult (P60) WT versus the postnatal (P10) WT mouse brain. **a** Schematic defining the extravascular distance as the shortest distance of any given voxel in the tissue to the next vessel structure. **b** Histogram showing the distribution of the extravascular distance in P10 WT and P60 WT animals. P60 WT animals show an increased relative frequency of shorter extravascular distances as compared to the P10 WT situation (bin width = 10; number of bins = 20). **c-h** Color map indicating the extravascular distance in the cortices, hippocampi and superior colliculi of P10 WT and a P60 WT mice. Each voxel outside a vessel structure is assigned a color to depict its shortest distance to the nearest vessel structure. The reduced extravascular distance in P10 WT animals as compared to the P60 WT animals is obvious. The color bars indicate the shortest distance to the next vessel structure. **i-k** Quantification of the extravascular distance in P10 WT and P60 WT in the three brain regions calculated by global morphometry analysis. The extravascular distance in the cortices (**i**), hippocampi (**j**) and superior colliculi (**k**) of P60 WT animals was significantly decreased as compared to the P10 WT animals (n = 4-6 for P10 WT; n = 8-12 P60 WT animals were used; in average three ROIs per animal and brain region were analyzed). All data are shown as mean distributions where the open dot represents the mean. Boxplots indicate the 25% to 75% quartiles of the data. The shaded blue and red areas indicate the SD. *P<0.05, **P<0.01, ***P<0.001. Scale bars: 100 µm (**c-h** overviews); 50 µm (**c-h** zooms).

**Figure 10.**
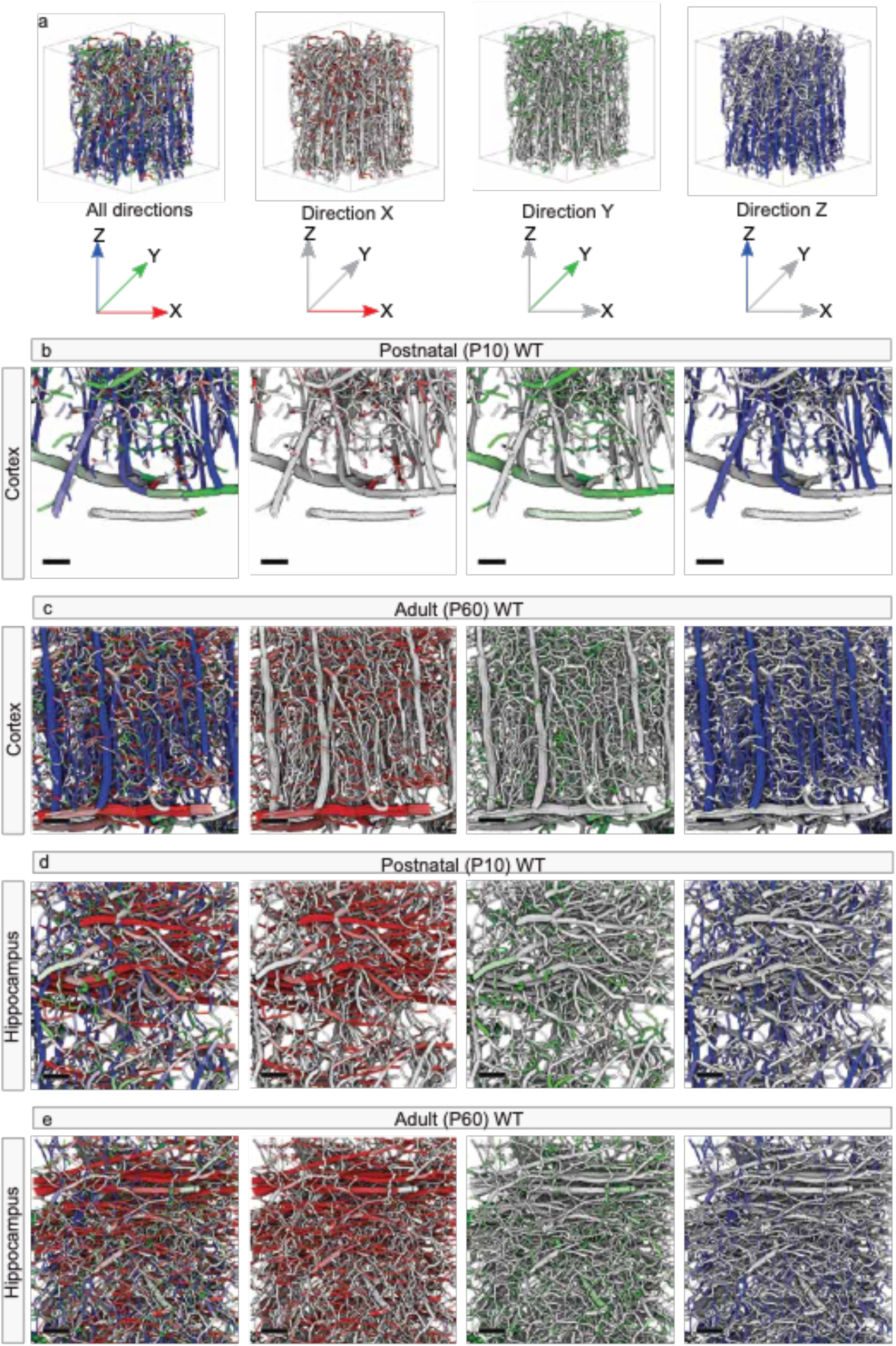
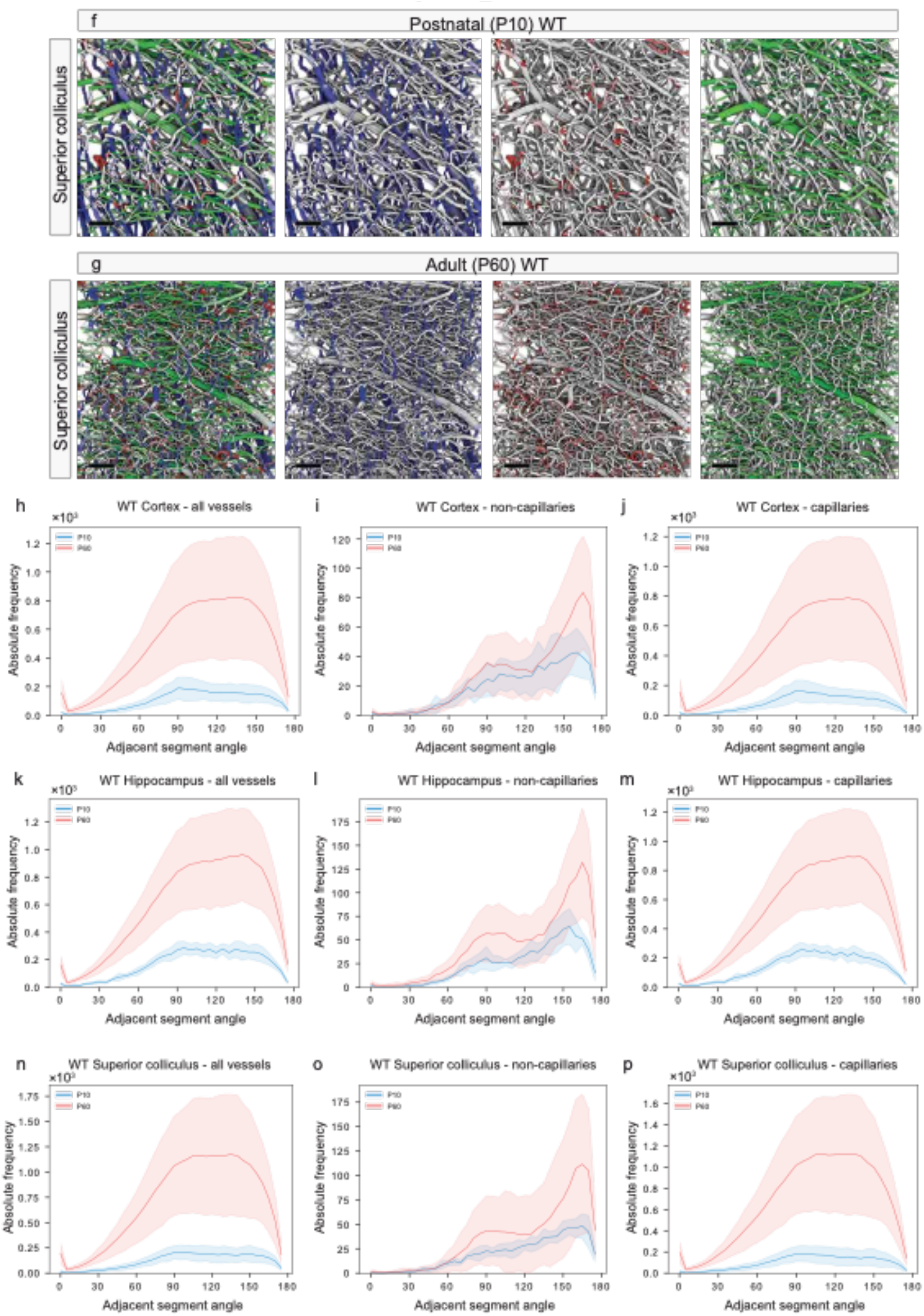
Global vascular network morphology – visualization of distinct patterns of vessel directionality in various brain regions of the adult (P60) WT and postnatal (P10) WT mouse brain. **a** Computational 3D reconstruction of µCT scans of cortical vascular networks of P60 WT displayed with color-coded directionality obtained by attributing every vessel segment to its main direction in the x (red), y (green), and z (blue) planes within the selected ROIs. **b,c** Computational 3D reconstruction of µCT scans color-coded for vessel directionality depicting the P10 WT (**b**) and P60 WT (**c**) cortex. Cortical renderings clearly show that the superficial perineural vascular plexus (PNVP) extended in the x- and y-directions, whereas the interneural vascular plexus (INVP) exhibited a radial sprouting pattern into the brain parenchyma along the z-axis, perpendicular to the PNVP. **d,e** In the hippocampus, horizontally orientated main vessel branches in the x-axis were recognizable, most likely following the distinct curved shape of this anatomical structure. **f,g** In the superior colliculus, a comparable pattern of directionality as in the cortex was observed with recognizable INVP vessels radially sprouting into the brain parenchyma along the z-axis. **h-p** Quantitative analysis showing histograms for the distribution of the relative angles of the vascular segments emanating from a certain branch point (segment angle for adjacent vessels) in P10 and P60 animals in the cortex (**h-j**), hippocampus (**k-m**), and superior colliculus (**n-p**). Non-capillaries revealed two main angles of orientation, namely around 90° and at around 170° in the three brain regions examined (**i** for cortex, **l** for hippocampus, **o** for superior colliculus), and these two main angles of orientation were more pronounced at the adult P60 as compared to the developing P10 stage (**i,l,o**), and were very similar in the three brain regions, suggesting a more general underlying concept of vessel directionality/orientation. The capillaries derived at angles between around 75° and 150° at both the developing P10 and mature P60 stages in the cortex, (**j**), hippocampus (**m**), and superior colliculus (**p**). (**h-p** bin width = 5; number of bins = 55) Scale bars: 100 µm (**b-g**).

### Experimental animals

To illustrate the potency of this technique in detecting the effects of different molecules, proteins and drugs on 3D vessel morphology, we quantified the difference in 3D vessel morphology between healthy P10 and P60 mice as well as wildtype and Nogo-A^−/−^ mice of either age. Although we have used P10 and P60 animals, the present protocol is applicable to mice of any postnatal and adult stages. Importantly, as genetic or other experimental manipulations such as intraperitoneal antibody- or tamoxifen injections were described to affect the growth and thus the weight of animals^121^, it is crucial to weigh the animals prior to resin perfusion. Although genetic modifications are not necessarily reflected to the same extent in all organs, similar weights and ages were chosen as parameters to ensure comparable developmental stages of mice between the different study groups. This is especially true for mouse pups at the early postnatal stage because the vasculature develops relatively fast at that stage^38^. Therefore, careful control of the developmental stage of the mice (including age and weight of the animals) has to be taken and is a prerequisite for any study. Since various genetic modifications were shown to alter the overall postnatal development, animal weight match constitutes an important parameter with regard to vessel structure analysis^122^. Moreover, littermate controls should always be used where available.

Power calculations based on the expected or biologically relevant difference between the groups should be made prior to the experiments. In principle, more pronounced differences between two test groups require a smaller number of animals to detect the effect and vice versa. Incorporation of appropriate controls, for instance, brains from postnatal wild-type littermates, is essential for analyses of angiogenesis and vascular network architecture in genetically modified mice during postnatal CNS development as well as in adulthood. All animal procedures must be carried out in accordance with the guidelines outlined by the institutional and local review committees for animal experiments.

### Vascular corrosion casts - Resin PU4ii infusion

The used resin PU4ii (vasQtec, Zürich, Switzerland) is a polyurethane-based casting resin with superior physical and imaging characteristics, timely polymerization, minimal shrinking and low viscosity, which guarantees perfusion into the finest capillaries^105^. These casts are highly elastic while retaining their original structure to facilitate post-casting tissue dissection and pruning^105^.

The adapted perfusion protocol had to be optimized for postnatal animals in order to fulfil two requirements: i) to perfectly fill the vessels with PU4ii, ii) to simultaneously preserve the mechanical vessel integrity in order to avoid rupture of the more immature vessels (Extended Data Figure 2).

Therefore, mice (P10 or P60) are first intracardially perfused through the left ventricle with 10– 20 ml ACSF containing 25,000 U/L Heparin, followed by 4% paraformaldehyde in PBS, and then by resin PU4ii (vasQtec, Zürich, Switzerland), at 90-100 mmHg for postnatal animals and 100-120 mmHg for adult mice. This adapted infusion pressure for P10 mice as compared to adult animals ensured mechanical stability preventing vessel rupture while ensuring at the same time adequate perfusion of the blood vessel network.

### Resin curing, maceration/decalcification and brain dissection

After resin curing of the perfused mouse at room temperature for at least 24 hours (embedded in a diaper), the tissue is macerated in 7.5% KOH at 50°C for 24 hours followed by decalcification of the surrounding bone structures with 5% formic acid, also at 50°C for 24 hours (Figure 2k,l, and Extended Data Figure 1e,f). The cerebral vasculature is then dissected from the remaining extra-cranial vessels (Figure 2k-n, and Extended Data Figure 1g,h). Subsequently, the cerebral vascular corrosion casts are washed thoroughly in distilled water and dried by lyophilization.

### Imaging and quantification

#### Scanning electronic microscopy (SEM)

For qualitative control of the vascular corrosion casts, we inspected the perfused blood vessels of the entire mouse body including the whole brain by visual observation and conventional light microscopy (Extended Data Figure 2). Then, a subset of the casts was mounted on stubs and sputter-coated with gold^109^ for quality control scanning electron microscopy imaging (SEM; Hitachi S4000)^109,120^. Imaging of resin-perfused, dissected and osmicated P10 and P60, WT and Nogo-A^−/−^ brain tissue samples revealed a dense vascular network in the superficial cortex with well-defined vessel structures of all sizes (Extended Data Figure 2). Moreover, the entire brain vasculature was uniformly filled and the vessel lumen was molded with its finest details (Extended Data Figure 2).

In addition, larger blood vessels showed oval endothelial nuclei imprints on the resin cast surface, a quality feature also observed in vascular corrosion casts of adult mice demonstrating the perfect molding of the vascular lumen, suggesting complete filling (Extended Data Figure 2)^110^. The fine capillary network was well visible and devoid of vessel interruptions in the selected P10 and P60 brain cortices (Extended Data Figure 2).

Taken together, all these morphological blood vessel features indicate that the vascular corrosion casts were of excellent quality.

#### Desktop µCT imaging and region of interest selection

Three-dimensional images of whole-brain samples were acquired using a desktop μCT system (μCT 40, Scanco Medical AG). Details can be found in PROCEDURE part of this paper. The images were evaluated visually and desired ROIs in various anatomical regions of the brain were chosen. To assure precise navigation within the postnatal mouse brain parenchyma, we refer to the Developing Mouse Allen Brain Atlas (http://developingmouse.brain-map.org/docs/Overview.pdf) or – alternatively – to any other Mouse Brain Atlas. In order to find the same ROIs in the SRµCT imaging (see below), the coordinates of the ROIs were determined relative to well-distinguishable details in the sample. For this purpose, we employed two metallic pins in the sample holder rod. When scanning the entire vascular corrosion cast, ROI selection can be omitted.

#### Synchrotron radiation-based micro CT (SRμCT)

High-resolution (pixel size 0.73 µm) SRµCT images of the selected ROIs were acquired at the TOMCAT beamline of the Paul Scherrer Institute (Switzerland). The ROIs were located using the steel pins in the sample holder rod as described above. X-ray projection images were acquired with monochromatic X-rays. Paganin phase retrieval^59,123^ was performed before tomographic reconstruction using the Gridrec algorithm^59,124^ (more details can be found in the PROCEDURE).

Additionally, high-resolution images of selected whole-brain casts were acquired using a similar setup as compared to the ROI-based approach, now with 0.65 µm pixel size. Here, the field of view of a single tomographic image is too small to cover the whole brain, and thus we used mosaic imaging mode where multiple smaller, partially overlapping, images are acquired. The reconstructions are then stitched into a single large image representing the whole brain vasculature^125^. The SRμCT images had an SNR > 10 (and typically in the range 20-30), and therefore they seemed to be well suited for computational reconstruction of the vascular network.

#### Alternatives for broader accessibility

In this protocol, we propose to use a desktop μCT scanner to acquire a low-resolution overview of the whole brain sample after which we use a SRμCT imaging approach to study the selected ROIs at a high-resolution in more detail (Figure 4a-l). Alternatively, both the low-resolution ROI selection and the high-resolution imaging of the selected ROIs (Figure 4a-l) can be performed using SRμCT techniques without the use of a desktop μCT scanner, for instance in case of well accessible SRµCT scanners and in order to avoid relocalization in different CT machines. As another alternative, all the imaging steps in this protocol can also be performed with a high-resolution desktop μCT scanner only (machines from manufacturers such as Scanco, Bruker, Zeiss, Phoenix and Nikon can be used) without the use and need of an often less accessible SRμCT scanner, and if the number of samples is relatively small as scanning times are higher with desktop μCT scanners.

### Computational reconstruction of the 3D vascular network

#### Vessel segmentation

The vessels in the SRμCT reconstructions were segmented to yield a binary image where the foreground represents vessels and background represents the structures in between them (Figure 4m-p). The high SNR allowed us to use simple Otsu thresholding^59,126^, where all pixels with a value above a specific, algorithmically determined, threshold are classified as vessels (Figure 4o). In the Otsu method, the threshold was chosen such that it minimizes the intensity variance of pixels inside both groups. The threshold value does not seem to be very sensitive to the selection of the thresholding method, provided that the selected method is suitable for thresholding images with bivariate histograms. For example, other previously described methods^127–129^ typically give threshold values that are within a 10% range of the value given by the Otsu method.

The thresholding step was followed by morphological closing using a spherical structuring element (radius = 0.75 µm) in order to eliminate small gaps in the vessels. A few regions primarily located in the largest vessels contained small, isolated background regions (i.e. holes or bubbles) that were eliminated by merging all isolated background regions to the foreground.

#### Computational reconstruction of the 3D vascular network

In order to analyze the individual vessel branches, we converted the segmented image into a graph representing the vessel network. To this end, the segmented vessels were first reduced into a one-pixel wide skeleton that approximated the centerline of each vessel branch^59^. This was done by iterative thinning of the structures until only lines were left^59,130^ (Figure 4o,p). The skeleton was traced into a graph structure, where each graph node corresponds to an intersection point between three or more skeleton branches, and the branches are represented by edges between the nodes. For each edge, the locations of the skeleton pixels that form the corresponding branch were stored. Additionally, for each stored location, the distance to the nearest non-vessel pixel was calculated in order to determine the (local) radius of the vessel^59,131^.

An anchored convolution approach was used to smooth the pixel-accurate representation of the vessel branches for more accurate length measurements and visualizations^59,132^. In the anchored convolution the locations of the points forming the branch are convolved with a Gaussian kernel, but the points are not allowed to move more than a specific distance from their original locations. As a result, small deviations in the curve will smooth out but larger ones remain. In our case, the standard deviation of the smoothing Gaussian was 1.125 µm, and the maximum distance a point was allowed to move was 0.5 µm. The endpoints of each branch were not allowed to move at all.

Finally, spurious branches originating from the ill-conditioned nature of the skeletonization procedure and from remaining imaging noise were removed using a simple heuristic. Therefore, all branches with at least one free end and a length of less than twice the radius at the non-free end were removed. Additionally, isolated branches less than 5 µm in length were erased, as these corresponded to small, spurious foreground regions not representing vessels.

The image analysis was made using the freely available pi2 software (version 3.0) (https://github.com/arttumiettinen/pi2) and Python 3.6.

#### Global morphometry and local topological vascular network analysis

Global morphometry and local topological vascular network analysis were performed for each high-resolution image in accordance with our previous work^40^. Global morphometry analysis is based on the vessel segmentation map, it considers the characteristics of the vascular structure on a per-voxel level, and allows to assess the vascular parameters vascular volume fraction and extravascular distance.

In contrast to global morphometry analysis, using local topological analysis of the 3D vascular network individual per-vessel parameters like branch point density and - degree, vessel/segment density, -diameter, -length, -tortuosity, -directionality, and can be calculated in addition to the vascular volume fraction.

The nodes in the previously created graph were interpreted as branch points and the edges in between nodes as vessel segments. Branch points are junctions between at least three vessel segments where the number of vessel segments joining in one branch point is called branch point degree (which equals at least 3). The branch points and all segments were categorized into capillary (diameter < 7µm), non-capillary (diameter ≥ 7µm) and global (without distinction by diameter) being the union of capillaries and non-capillaries.

We defined the branch point diameter as the diameter of the thickest joining segment, allowing the discrimination between capillary and non-capillary branch points. The branch point density per volume was defined as the number of branch points per cubic millimeter (Figure 7, and Extended Data Figures 5–9).

The number of branch points per degree was counted and separated for capillaries, non-capillaries and global branch points in order to plot branch point degree and branch point diameter statistics (Extended Data Figures 7–9). The values per branch point degree were then divided by network volume to obtain branch point density statistics (Figure 7c,f-h, and Extended Data Figures 5,6).

All segments were classified into either capillary or non-capillary, counted and divided by network volume to plot segment density statistics (Figure 8d-f, and Supplementary Figure 4a-c). Segment diameter was obtained as the average value of local vessel diameter over all the points defining the segment (Figure 8g-j, and Supplementary Figure 4d-f). Segment length was calculated as the sum of lengths of line segments between points defining the vessel segment (Figure 8k-n, and Supplementary Figure 4g-i). Tortuosity was calculated as the ratio between the real length of a segment and the shortest distance between the segment’s endpoints (Figure 8o-r, and Supplementary Figure 4j-l). In order to calculate segment volume, each of the line segments was assumed to represent a cylinder whose diameter was taken to be the average vessel diameter at its endpoints, and the sum of all the cylinder volumes gave the segment volume (Figure 8s-v, and Supplementary Figure 4m-o).

In order to determine extravascular distance, the Euclidean distance transform of the background in the segmented image was calculated. The distance transformation gives the distance to the nearest vessel for each background point (Figure 9). Binning the distance values into a histogram gives an estimate of the extravascular distance distribution (Figure 9b,i-k, Supplementary Figure 5).

The volume fraction of global segments was determined as the ratio between the count of vessel voxels and the total count of voxels in the segmented image (Figure 5,6l-qt-v, and Extended Data Figures 1,2). This value was then divided into capillary- and non-capillary volume fractions in proportion to the total volumes of vessels in these two vessel groups (Figure 6l-q,t-v). Quantitative analysis of the relative angles of the vascular segments emanating from a certain branch point was performed for both the developing P10 and mature P60 stages (Figure 10h-j). Three-dimensional renderings of the computationally reconstructed vessel networks were obtained, color-coded by 1) the average vessel radius (Figure 5e-g), 2) the division to capillary and non-capillary vessels (Figure 6h-k,r,s), and 3) vessel directionality (Figure 10b-g).

### Statistical analysis

For the ROI-based approach, the mean of the parameters obtained from the animals in the different groups (P10 WT, P10 Nogo-A^−/−^, P60 WT and P60 Nogo-A^−/−^) were compared using the Welch t-test. Histograms and boxplots were used to visualize the data (Figures 5–10, Extended Data Figures 5–9, and Supplementary Figures 1–5), with whiskers spanning at most 1.5 times the inter-quartile-range (IQR, the height of the boxes) above and below the 25% and 75% quartiles. Whiskers are drawn at the highest/lowest data value if it is inside the 1.5 IQR. In compliance with the standard^133^, the open dot represents the mean, the horizontal line represents the median^133^. Both statistical testing and figure plotting were performed with Matplotlib (https://matplotlib.org) and NumPy (https://numpy.org). For all quantitative analysis in the different brain regions (Figures 5–10, Extended Dat figure 5–8, and Supplementary Figures 1–5) we used 4-6 postnatal (P10), and 8-12 adult (P60) animals. On average, we have analyzed about three ROIs per brain region.

In addition to the ROI-based approach and in order to visualize the variability of the measured quantities in a single brain specimen, we used whole-brain images acquired for one P10 WT and one P60 WT sample. The whole-brain images were manually registered to the Allen Mouse Brain Atlas and divided into regions of the same size as the ROIs used earlier. The vascular network in the cortex, hippocampus, superior colliculus, corpus callosum, hippocampal subregions – CA1, CA3, and dentate gyrus –, basal ganglia, thalamus, hypothalamus, olfactory bulb, cerebellum, pons, and medulla were analyzed using the same approach (Figure 8, and Supplementary Figures 10–14). The anatomical regions were divided into 5-126 parts depending on the size of that region and quantified similarly to the ROI-based approach. The mean of the parameters obtained from one animal in the different groups (n=1 for P10 WT, n=1 for P60 WT) were compared using the Welch t-test. Histograms and boxplot to visualize the data (Figure 8, and Supplementary Figures 10–14), as well as statistical testing and figure plotting were performed in the exact same manner as described for the ROI-based approach above.

### Advantages of the protocol compared with alternatives

The hierarchical imaging of vascular corrosion casts combined with the computational analysis at the level of individual vessel segments up to entire vascular networks in both the postnatal and adult mouse brain as described here has several key advantages as compared to alternative methods:

- This method enables visualization and quantitative assessment of the structure of blood vessel networks in the developing postnatal as well as in the entire adult mature mouse brain down to the level of capillaries, with yet unmet precision (Figures 5–11).
- The brain microvasculature has proven hard to image due to the small size of the capillaries and large regions of interest required for network-level analysis. Traditional, non-invasive, in-vivo imaging modalities such as Doppler tomography^134,135^, μMRI^136^ and magnetic resonance angiography^137^ have a resolution in the range of tens of micrometers but that is not high enough to detect the microvasculature (compared to the near 1 μm resolution in μCT imaging). Confocal single-photon and two-photon imaging can provide the required resolution but have depth limitations and its use *in-vivo* often require cranial windows^21,24^, whereas other, classical (automated) histology-based methods are often limited to smaller regions, can be disturbed by deformations of the tissue slices especially at high resolutions (thin slices) and are therefore not suitable for quantification of the brain vasculature on a network level^138–140^. Recently, light-sheet fluorescence microscopy (LSFM) of the stained blood vessel network in both cleared non-CNS^141,142^ and in cleared CNS tissue^143–146^ has received increasing attention. LSFM has proved to be a very attractive and efficient method for imaging of large vessel networks with relatively high resolution. Contrary to corrosion casting, the LSFM technique requires optically clear or cleared samples (for the excitation wavelength). In corrosion casting, there are no such requirements and therefore no possibility of typical LSFM artefacts such as shadowing or photobleaching. Routinely achievable isotropic resolution of µCT techniques is near 1 µm, but in LSFM the resolution is often anisotropic. For example, here we have an isotropic voxel size of 0.65 µm, compared to 1.625 µm x 1.625 µm x 3 µm achieved through multi-dye vessel staining, tissue clearing, and 3D LSFM in^145^. For analysis, anisotropic voxels are often resampled to isotropic voxel size corresponding to the lowest resolution component, as was also done in^145^, resulting in isotropic voxel size of 3 µm for convolutional neural network vessel segmentation and subsequent analysis of the vasculature of the whole mouse brain^145^.
- Micro CT imaging with use of intravascular contrast agents is used both in-vivo and ex-vivo and made it possible to image the vasculature in whole organs in 3 dimensions with increasing resolution^107,147–150^. Ex-vivo, vascular corrosion casts have shown to be an excellent method to investigate brain vasculature in particular^40,104,106,107,109,110,120^. Corrosion casting, in combination with recent advances in imaging techniques – both for SRμCT and desktop μCT setups –, as used in this protocol, results in a resolution that is at least 3 times higher (0.65 μm vs 2 μm pixel size) than another recent paper that used a similar approach to study the mouse brain vasculature^107^.
- The protocol used here allows to quantitatively describe in detail the postnatal and adult CNS blood vessel network down to the capillary level, (Figures 5–11 and Extended Data Figures 3–9), including visualization and quantification of vessel directionality in the X, Y, Z planes (Figure 10, and Supplementary Figures 6–9,14), which provides important additional information and – to the best of our knowledge – has not been addressed so far. Moreover, it permits to distinguish between capillaries and non-capillaries using clearly defined morphological parameters. In this paper we classified a vessel as a capillary if its inner diameter was less than 7 µm (Figures 3,6–9). Similar morphological classification is more difficult and tedious with other, classical (automated) histology-based methods due to dye color variations, staining artefacts, and distortion of the true vessel diameter by perivascular cells^138–140^. Histology-based methods including recently published LSFM based methods^141–145^ are also less precise in addressing inner vessel diameters.
- Hierarchical imaging and computational reconstruction of the 3D vascular network via global vascular network morphometry and local vascular network topology allow accurate, complete and quantitative analysis and description of the 3D vessel structure in the postnatal- and the adult mouse brain at the network level. These computational 3D analyses result in vessel parameters (Figures 5–9, and Extended Data Figures 3–9) with direct biological relevance instead of single measures such as ‘percentage of section area occupied by vessels’, that are obtained in classical 2D-section analysis methods. For instance, maximum intensity projections of tissue slices are often used, even though they lead to an overestimation of the vessel density, as the vascular volume fractions of multiple optical sections are summed up, an error that adds up with increasing section thickness. A 3D analysis resulting in true vessel volume fractions, therefore, provides more accurate quantifications of the vascular networks and also allow better comparison between different studies. Three-dimensional analyses can be either obtained by combining vascular corrosion casting techniques with computational analysis as described here, or by stereological methods as we^27^, and others^151^ have previously reported.
- In contrast to section analysis (2D, maximum intensity projections), computational analysis of 3D hierarchical images allows – similar to image analysis obtained from confocal *z*-stacks using stereological methods^27^ – characterization of additional parameters of the vascular network such as vessel volume fraction, branch point density and -degree, vessel density, -diameter, -length, -tortuosity, -volume, -directionality, and extravascular distance (Figures 5–9, and Extended Data Figures 3–9).
- Our protocol allows clear identification of capillary (inner vessel diameter < 7µm, Figure 6a,b) and non-capillary (inner vessel diameter ≥ 7µm, Figure 6a,b) blood vessels based on the morphological criteria of the inner vessel diameter (Figures 6–8, Extended Data Figure 4–9, Supplementary Figures 1–8,10–12,14). This biologically important distinction is more difficult to address with classical immunofluorescence^138–140^, *in vivo* imaging modalities such as μMRI^136^, μCT^150,152^, contrast-enhanced digital subtraction and CT angiography^153^, as well as with more recently developed methods such as LSFM analyses, with which vessel diameter cannot be determined as adequately. Moreover, there is also a lack of reliable capillary markers^142–145^ which otherwise might solve this issue for staining-based methods. Indeed, markers for capillary endothelial have been proposed in recent years, for instance, Mfsd2a^154^, TFRC, SKC16a2, CA4, CXCL12, Rgcc, SPOCK2 and others that were recently identified in large endothelial single-cell-RNA sequencing datasets^155–157^. However, to the best of our knowledge, there is currently no established cellular marker that selectively labels capillaries, as all the proposed and above-mentioned markers also label non-capillary blood vessels, even though to a lesser extent^156,157^. Accordingly, the most adequate method for distinguishing capillary and non-capillary blood vessels is still based on the morphological criteria of the inner vessel diameter, e.g. < 7µm for capillaries versus ≥ 7µm for non-capillary blood vessels (Figures 6–8, Extended Data Figure 4–9, and Supplementary Figures 1–8,10–12,14).
- High-resolution hierarchical imaging and computational analyses enable the 3D visualization, quantification and characterization of the entire vascular brain network. These analyses permit to obtain various quantitative parameters at the level of both the entire vascular network as well as of individual vessel segments, such as the volume fraction, extravascular distance, branch point density and -degree, vessel density (entire vascular network), -diameter, -length, -tortuosity, -volume and -directionality (individual vessel segment) (Figures 5–10, Extended Data Figures 5–9, and Supplementary Figures 1–5,10–14). Moreover, the number of vessel branch points enables to better understand the 3D aspects of vessel branching within the process of sprouting angiogenesis (Figure 7, Extended Data Figures 5–9, and Supplementary Figures 3,11), which constitutes the predominant model of vessel formation in the brain^3,27,40^. Interestingly, the combination of vessel branch point analyses with the above-mentioned distinction between capillaries and non-capillaries allows revealing underlying mechanisms of sprouting angiogenesis in different brain regions such as cortex, hippocampus, and superior colliculus (Figures 6–7, and Extended Data Figure 5–9). For instance, branching events emanating from the penetrating vessels of the INVP in the cortex can clearly be visualized and quantified (Figure 6h,i). Such computational/quantitative analyses are not commonly feasible using either classical staining-based methods^138–140^ or novel methods including LSFM^141–145^.
- The method presented here further allows to address functional consequences of the vascular network formation such as the extravascular distance – e.g. the shortest distance of any given point in the tissue to the next vessel structure (Figure 9, and Supplementary Figures 5,13), which are more difficult to address throughout the whole brain with alternative methods.
- In principle, for simultaneous analysis of vasculature and the surrounding tissue, our protocol can also be combined with immunofluorescence using various vascular markers, e.g., pericyte markers (PDGFR-β, CD13), astrocyte markers (GFAP) or neuronal markers (Nf160 and Tub-βIII), to visualize perivascular cell types. This can be done by sectioning the resin perfused brains without tissue maceration and bony structure decalcification, thereby leaving the brains intact, followed by immunofluorescence staining. By mixing in specific fluorescent dyes with the resin solution, it is also possible to improve and specify the inherent fluorescence of PU4ii (Extended Data Figure 10). Accordingly, virtually any protein of interest can be analyzed in the morphological context of the developing postnatal and adult vascular network, without the risk of collapse or deformation of vascular structures that is often encountered with a staining-only, histological approach^138,140^.
- Recent technical improvements allow scanning the whole postnatal and adult brain vascular corrosion cast at once with the same resolution as the method involving ROI selection prior to the scan. This whole-brain scan allows to analyze any region of interest, even after SRμCT scanning (Figure 8, and Supplementary Figures 10–14).

**Figure 11.**
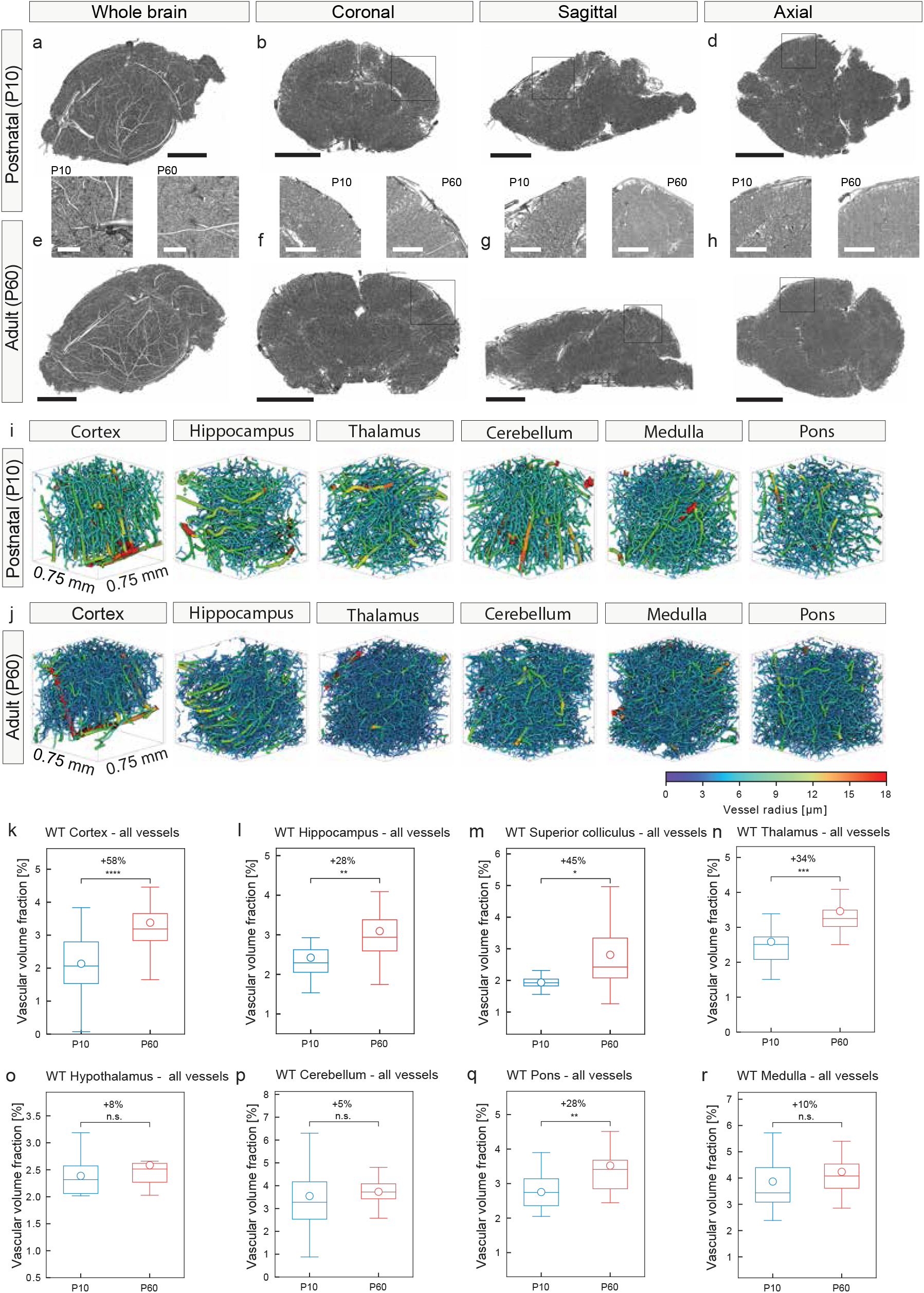
High resolution SRµCT scan of the entire postnatal (P10) and adult (P60) mouse brain – visualization and quantification of various anatomical brain regions reveals increased vascular volume fractions in the P60 versus P10 mouse brain. Cross sections of the whole-brain scan in both P10 (**a-d**) and P60 (**e-h**) mouse brains with 3D overviews (**a,e**) and detailed panels for the respective anatomical planes (**b,f** coronal, **c,g** sagittal, **d,h** axial). Computational 3D reconstructions of µCT scans of vascular networks of the P10 WT and P60 whole brain scans, displayed with color-coded vessel thickness. The increased vessel density in the P60 WT (**j**) cortex, hippocampus, thalamus, cerebellum, medulla and pons, as compared to the P10 WT (**i**) samples is obvious. Color bar indicates vessel radius (µm). **k-r** Quantification of the 3D vascular volume fraction in P10 cortex (**k**), hippocampus (**l**), superior colliculus (**m**), thalamus (**n**), hypothalamus (**o**), cerebellum (**p**), pons (**q**), medulla (**r**) by computational analysis using a global morphometry approach. The vascular volume fraction in all these brain structures was higher in the P60 WT animals than that in the P10 WT animals (the anatomical regions were divided into ROI-sized sections for analysis: between n = 5, dentate gyrus, and n = 126, cortex for P10 WT; and between n = 4, CA3, and n = 76, cortex, of these sections for P60 WT were analyzed, highly dependent on the size of the anatomical region). All data are shown as mean distributions, where the open dot represents the mean. Boxplots indicate the 25% to 75% quartiles of the data. *P < 0.05, **P < 0.01, ***P < 0.001. Scale bars: 2.5 µm (**a-h**, overviews), 0.25 µm (**a,e,** zooms), 1 µm (**b-d, f-h,** zooms).

**Figure 12.**
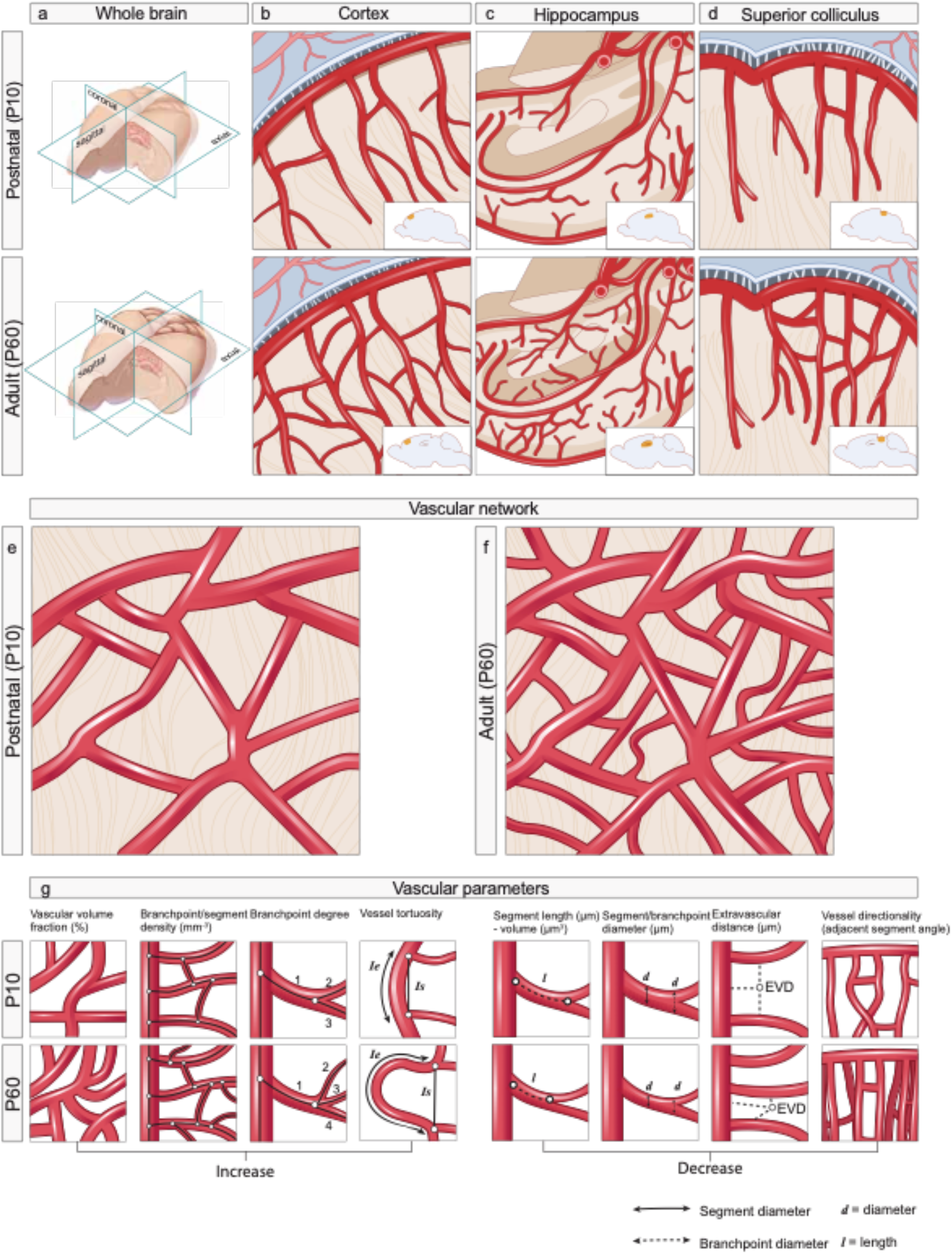
Working model summarizing the differences in 3D vascular network architecture between adult (P60) WT and postnatal (P10) WT mice. **a-d** Schematic illustrations of the postnatal and adult mouse brain vasculature, both as 3D whole brains (**a**) and as enlargements specified for the different anatomical regions, namely cortex (**b**), hippocampus (**c**) and superior colliculus (**d**). **e,f** From the P10 to the P60 stage, an increased vascular volume fraction of functional blood vessels established via increased vascular endothelial sprouting, branching, and -migration mainly at the capillary level can be observed (**e,f**). **g** Schematics summarizing the most important differences in the 3D vascular network architecture of P10- and P60 mouse brains: The vascular volume fraction (percentage of tissue volume occupied by the entire vascular network), the branchpoint- and vessel/segment density, the branchpoint degree, and the vessel/segment tortuosity are increased in the mature P60 as compared to the developing P10 animals. On the other hand, the vascular parameters vessel/segment length, vessel/segment volume, vessel/segment- and branchpoint diameter, and the extravascular distance are decreased in the mature P60 as compared to the developing postnatal P10 mice. The vessel directionality showed more pronounced angles of orientation for non-capillaries at the P60 stage as compared to P10 stage, whereas the orientation of the capillaries seemed largely unchanged between the two developmental stages.

### Limitations of the method

Despite the advantages described above, this method has some disadvantages and limitations:

- In contrast to classical immunofluorescent-based methods, the technique described here is more labor-intensive. For instance, the imaging and analysis of the superficial vascular plexus of the postnatal retina^121,158^ or of the embryonic hindbrain^32^ – which both form a 2D flat vascular structure and are the most commonly used *in vivo* angiogenesis models – can easily be assessed, in principle even with a standard fluorescence microscope. The quantitative analysis of angiogenesis in the postnatal mouse brain^27^ – in which the vasculature develops in 3D with vessel sprouting occurring in all directions – requires confocal imaging for proper analysis, thus resulting in longer image acquisition times and larger data sets as compared to the postnatal retina and embryonic hindbrain models^32,121^. The method using confocal imaging is similar regarding labor-intensiveness to the protocol described here but cannot be applied at once to the whole brain vasculature.
- Comparable to the immunofluorescent-based analysis of the postnatal mouse brain^27^, the structural complexity of the postnatal- and adult mouse brain angioarchitecture requires meticulous navigation within each individual sample to assure that really equivalent brain areas are selected. This is crucial as vascular density and angiogenic activity vary considerably among different brain regions^38,40,107,145^. Navigation within postnatal mouse brain samples as prepared with the present protocol is hampered because even though the combination of vascular corrosion castings with histological staining (such as e.g. Nissl or H&E stains, which would disturb the immunofluorescence) is feasible (Extended Data Figure 10), these different kinds of sample preparation preclude the 3D hierarchical imaging as described. Therefore, for detailed navigation (and ROI placement, see Figure 4a-l) throughout the postnatal and adult mouse brain, we recommend referring to the Developing- and Adult Allen Mouse Brain Atlas or to other commonly used Mouse Brain Atlases (Figure 4a-l).
- The described technique of vascular corrosion casting and computational 3D analysis does not enable to directly identify endothelial tip cells or newly formed but yet immature, non-perfused blood vessels as a percentage of functional, perfused vessels since only the perfused part of the vessel tree can be analyzed. Endothelial tip cells and immature non-perfused blood vessels can, however, be addressed by combining vascular corrosion casting with classical immunofluorescence staining techniques (Extended Data Figure 10). Moreover, blood flow in the functional vessels cannot be measured. In light of the finding that an increased density of perfused blood vessels does not necessarily result in higher blood flow, as it has for instance been shown for transgenic overexpression of VEGF165 (= VEGF-A) in the adult mouse brain^108^, this needs to be taken into account.
- The protocol described here provides only snapshots of brain angiogenesis at given time-points for individual mice, and in consequence requires a relatively large number of animals in order to study the dynamics of angiogenesis and vascular network architecture over time. We have partially addressed this aspect here by comparing the P10 with the P60 mouse brain. Although methods to study *in vivo* angiogenesis in the developing postnatal and adult mouse brain are currently not widely available, methods for real-time imaging of the brain vasculature exist^24,41^, and may, in the future, complement our protocol.
- In contrast to the immunofluorescent-based methods addressing angiogenesis in the postnatal retina, in the embryonic hindbrain, and in the postnatal brain^27,121^, the identification of and distinction between endothelial tip cells, trailing stalk cells, and filopodia is not possible with vascular corrosion casting and computational 3D analysis, thereby limiting the ability to characterize patterns of sprouting angiogenesis and to differentiate between active and quiescent vessel sprouts. Our method allows the distinction between capillaries and non-capillaries based on their inner diameter (Figures 6–8, Extended Data Figure 4–9, and Supplementary Figures 1–8,10–12,14), and this is in principle possible for every other vessel of a given size (Figure 5, and Extended Data Figure 3). However, it does not permit to directly differentiate between arteries and veins or between arterioles and venules, which have similar inner vessel diameters but different functionalities^107^. The differentiation between arteries and veins is possible using SEM images revealing different shapes of nuclei impressions on the vascular cast depending on arterial versus venous flow (e.g. elongated nuclei impression in arteries versus more round nuclei impressions in veins), as we have previously described^105^. SEM, does, however, not allow for computational analysis of the vascular parameters. To circumvent these problems, the combination of vascular corrosion casting and immunofluorescence using classical markers for arteries (such as EphrinB2^159^, Jagged-1^160^, Alk-1^161^ etc.) and veins (such as VCAM-1^162^, COUP-TFII^163^, EphB4^164^ etc.) (Extended Data Figure 10) allows to address both 3D computational analysis of the vascular network as well as parameters and arterio-venous differentiation.
- Extravasation of intravascularly infused resin has been described in experimental liver^165^ and mammary^166^ tumor models. Although not described in brain vascular corrosion casting, and also not observed by us in the postnatal mouse brain with more angiogenic active vessels that are supposed to be leakier than mature vessels, some resin extravasation for example in vascular-dependent CNS pathologies associated with an impaired blood-brain barrier such as stroke, brain trauma or brain tumors cannot be excluded. Another limitation of the described protocol is the invasive nature of the technique as the mice need to be sacrificed for resin perfusion (Figure 2 and Extended Data Figure 1); similar to the other, above-mentioned techniques of mouse postnatal retina-, embryonic hindbrain-, and postnatal brain angiogenesis^27,36,121^. Vessel size index-MRI (VSI-MRI) has been proposed as a quantitative, non-invasive MRI derived imaging biomarker, for the non-invasive assessment of tumor blood vessel architecture and in order to evaluate the effects of vascular targeted therapy. The technique was validated in high-grade human gliomas^167^ and a direct comparison between *in vivo* MRI and post mortem μCT vessel calibers showed excellent agreement in vessel size measurements^168^. However, the parameters addressed with VSI-MRI are limited to an estimation of fractional blood volume and blood vessel size (resolution of 1.875 x 1.875 mm with a total matrix size of 240 x 218 mm), and the additional vascular parameters such as vessel length, tortuosity, vessel directionality, branch point density and extravascular distance addressed with our protocol are not measurable with VSI-MRI techniques^168^.

## MATERIALS

### REAGENTS

**! CAUTION** Some of the chemical substances mentioned below are extremely harmful if inhaled and swallowed, and/or are irritating to the eyes, respiratory system and skin. They may cause sensitization of skin upon contact. Therefore, in principle, wear protective goggles, clothing and gloves as appropriate. Use the chemicals in a fume hood. Carefully read the safety data sheets of all chemicals used in this protocol.

Pentobarbital solution 1.0 mg/mL in methanol (Sigma Aldrich cat. no. P-010)

NaCl, 0.9% (wt/vol) in distilled water (250 ml) (BBraun)

Heparin Sodium Salt from Porcine, Grade 1A (Sigma Aldrich cat. no. H3393-25KU)

Paraformaldehyde (PFA; Sigma-Aldrich, cat. no. P6148)

PU-4ii Resin, Hardener, Color, Fluorescence (www.vasqtec.com)

Ethylmethylketon (EMK) EMPLURA® (Merck cat. no. 106014)

XTEND-IT®, dry Gas Blanket (Kaupo, cat. no. 09291-010-000012)

Potassium hydroxide (KOH) (1 kg) (Merck cat. no. 1050331000)

Formic Acid, 99% techn. (Fisher Scientific cat. no. 270480250)

Osmium tetroxide solution, 2% in water (5 ml) (Sigma Aldrich cat. no. 20816-12-0)

PBS, 0.1 M, or PBS tablets (Sigma-Aldrich, cat. no. P4417)

Mounting medium (Dako, cat. no. 53023)

Instant adhesive (Cyanoacrylate) (Kisling, Ergo, cat. no. 5889)

Distilled water (Millipore, Milli-Q synthesis A10)

10-d-old and 2-month-old mice (wild type WT/Bl6, male or female)

**! CAUTION** Institutional and governmental ethics regulations concerning animal use must be followed. All the animal experiments herein were approved by the Veterinary Office of the Canton of Zurich.

### EQUIPMENT

Tabletop balance (e.g., ED Precision balance, Sartorius)

Plastic foil

Paper towels

Green protection diapers (Polymed Medical Center)

Gloves (M/L) Nitril (Sigma-Aldrich cat. no. Z543500/Z543519)

Syringes, 1 ml (BD Luer-Lok^™^ cat. no. 309628)

Syringes, 50ml (Fisher Scientific cat. no. 13-689-8)

Omnifix Luer-lock syringes (20ml) (BBraun cat. no. 4616200V)

Needles (0.5×16 mm/25G) (Gima cat. no. 23760)

Butterfly needles 23G and 25G (needle: 20mm, 0.6/0.4mm; tube: 300mm, 0.36ml) (Ospedalia AG)

Infusion pump (in house)

Dissecting instruments: gross/fine dissection scissors (Aichele Medico AG)

Dissection tray with corkboard (in house)

Pinning needles (1.2×40 mm/18G) (BBraun)

Beaker cups (120ml/160ml) (Semadeni cat. no. 2294 or 4438)

Pipette with pipette-tips (10ml) (Soccorex cat. no. 312.10)

Wheaton scintillation vials (20ml) (Fisher Scientific cat. no. 03-340-25N)

Vacuum pump (in house)

Lyophilization Machine (benchtop freeze dry system, Labconco)

Light microscope (in house)

Scanning Electron Microscope (Hitachi S4000)

Gold sputter / coater device (Agar Scientific)

Desktop μCT (μCT 40, Scanco Medical AG)

Synchrotron Radiation μCT (TOMCAT beamline of the Swiss Light Source)

### REAGENT SETUP

#### FA solution (4%, wt/vol in 0.1M PBS)

Under a chemical fume hood, prepare 0.1M phosphate buffer (pH = 7.4) by adding 19ml 0.2M NaH_2_PO_4_ x H_2_O (27.58g/l) to 81ml Na_2_HPO_4_ (28.38 g/l). Next, add 8 g of paraformaldehyde (PFA) powder to 100 ml distilled water (aqua millipore). Dissolve the PFA powder by heating the solution (containing a clean magnetic stir bar) to 60 °C on a magnetic stirrer. To facilitate dissolving the PFA powder, add few drops of 5 or 10M NaOH when mixing starts. After PFA powder dissolved completely, filter the PFA solution using filter paper. To dilute the 8% FA solution to the final concentration of 4% PFA in 0.1M PB, add 100 ml of 0.2M PB. When the solution has cooled down to room temperature, check the pH and adjust to pH 7.4 if necessary. Aliquots can be stored at -20 °C for up to 1 year ^36,121^, however, we recommend to freshly prepare the 4% PFA solution for every experiment. Thaw the aliquots in a warm water bath (35 - 40°C) and cool them to 4°C on ice before use.

#### Artificial Cerebrospinal Fluid (ACSF) (with 24KU Heparin Sodium Salt)

Under a chemical fume hood, mix NaCl (7.6 g/l), NaHCO_3_ (2.02 g/l), Glucose, MgSO_4_ x 7H_2_O (0.37 g/l), KCl (0.26 g/l), NaH_2_PO_4_ x H_2_O (0.17 g/l) (water-free 0.15 g/l), CaCl_2_ x 2H_2_O (0.29 g/l) (must be added last). Add 25KU Heparin Sodium salt per liter. If you prepare it one day in advance, store the solution at 4°C. Should be stored for no longer than 4 weeks^169^. Before use, warm up to 37°C in a water bath.

#### Resin (PU4ii)

Dilute 5-10 mg of pigment into 3 g of ethyl methyl ketone (EMK). Then add the EMK/pigment mixture to 5g of PU4ii and mix in a glass vial using a vortex mixer. The hardener (0.8 g) should only be added to the resin mixture after infusion of the formaldehyde fixation solution. Mix this final Resin mixture using a vortex mixer, briefly evacuated in a standard vacuum chamber to remove remaining air bubbles and then transfer into a 10 ml syringe. The hardener has the shortest shelf-life and should be covered with the dry gas (XTEND-IT®) (see reagent list) after each use.

### EQUIPMENT SETUP

#### Dissection board

A dissection board placed into a tray was used for the perfusion steps. The board should be slightly tilted to collect the efflux after cardiac infusion.

#### Light microscopes

We use conventional light microscopes for quality control of vascular corrosion casts.

#### Scanning electron microscope

We assessed the quality of random samples of the corrosion casts before μCT imaging using Hitachi S4000 SEM with backscattering detector. Multiple resolutions and acceleration voltages were used until clear views of the perfused vessels could be attained. The imaging parameters and exact type of the SEM are not critical as the images are used only for visual evaluation and pre-screening of possible less-well perfused samples.

#### Desktop μCT system

We used a cone-beam micro-tomograph to create 3D overview images of the whole brain samples and facilitate selection of suitable regions for high-resolution imaging. A Scanco Medical μCT 40 device with a nominal isotropic voxel size of 16µm was used in this protocol. The imaging settings were set to suitable values as recommended by the device manufacturer (e.g. for the system we used, a resolution of <8 μm 10% MTF and a 3 - 72 µm nominal isotropic voxel size with an image format of 512 x 512 to 4096 x 4096 pixels could be selected). The exact settings are not as critical as the images are not used for quantitative analysis. Similar instruments from other manufacturers can be used as well.

#### Synchrotron Radiation μCT

High-resolution tomographic images of selected regions of interest were acquired at the TOMCAT beamline of the Swiss Light Source. The imaging system was operated at 17.5 keV X-ray energy with a sample-detector distance of 30 mm. For each region of interest, 1001 projections images were acquired with 0.73 µm pixel size and 2s exposure time. Paganin phase retrieval was used on the projection data, followed by reconstruction with the gridrec algorithm. Similar experimental setups are available in most synchrotron facilities, and imaging settings should be selected according to the available instrumentation.

### PROCEDURE

#### Perfusion with Resin PU4ii (day 1) ● TIMING ∼5–10 min per mouse

1. After weighing, anesthetize mice using 100mg/kg pentobarbital intraperitoneally. Check proper anesthetic depth before proceeding (tail punch: mouse must not move anymore). Fix both the postnatal (P10) and adult (P60) animals onto a dissection board (Figure 2c-j, Extended Data Figure 1).

! CAUTION It is crucial to weigh the animals prior to resin perfusion as only similar weights (and ages) ensure comparable developmental stages of mice between the different study groups.

! CAUTION Animal procedures must be carried out in accordance with relevant institutional and governmental ethics guidelines.

2. Test with tail punch if the mouse is properly anesthetized and turn it on its back.

3. For the surgical preparation, then lift the skin with forceps and cut perpendicular to expose the sternum (Fig. 2g,d,g,h Extended Data Figure 1a),

4. Next, lift the sternum and make lateral cuts through the rib cage to expose the chest cavity. Pull back the cage flap and fix using a pin/needle. Carefully remove the pericardium using rounded forceps (Fig. 2d,h, Extended Data Figure 1b).

5. Lift the heart slightly using forceps and then insert butterfly needle (23G butterfly needle for adult mice, needle: 20mm, 0.6/0.4mm; tube: 300mm, 0.36ml, and 25G butterfly needle for P10 mice) into the left ventricle (Figure 2e,i, Extended Data Figure 1c). Note the angle of insertion of around 30°C in the sagittal plane (Figure 2e).

- CRITICAL STEP: Do not insert the needle too deep to avoid damage of the atrioventricular valve or puncturing the heart basis. The inserted perfusion needle should be glued to the heart using instant glue to prevent the needle from sliding out during perfusion and leakage of the perfusion fluids. Only a small drop is enough.

6. Open the right atrium with a pair of sharp and pointed scissors. Immediately start the perfusion with 20 ml (for adult animals) and 5 ml (for P10 animals) artificial cerebrospinal fluid containing 25,000 U/L Heparin through the left ventricle, then stop the pump, block the tube with a hemostat and exchange the syringe.

- CRITICAL STEP: Before reconnecting the syringe, make sure that all air in the tube is completely removed.
- CRITICAL STEP: for P10 animals, adapt the amount of ACSF containing 25,000 U/L Heparin according to the animal age.

? TROUBLESHOOTING

7. Start perfusion with 4% PFA (in PBS). Immediate muscle fasciculation indicates good quality of fixation.

8. Add the hardener (0.8 g) to Resin PU4ii solution (prepared by diluting 5-10 mg of pigment into 3 g of ethyl methyl ketone (EMK), adding the EMK/pigment mixture to 5g of PU4ii, see Reagent Setup) and mix using a vortex mixer (Figure 2b,f,j and Supplementary Video 1). Briefly evacuate in a standard vacuum chamber to remove remaining air bubbles and then transfer into a 10 ml syringe. When perfusing the Resin solution, tilt the rear of the perfusion pump upwards to prevent occasional air bubbles to leave the syringe. The tray should be tilted to avoid a mess due to the efflux from the atrium.

- CRITICAL STEP: The resin perfusion is successful if the mouse turns bluish immediately. This can be particularly well monitored at the paws, snout or tail (Figure 2f,j and Supplementary Video 1).

? TROUBLESHOOTING

#### Resin curing (days 2-3) ● TIMING ∼1 d

9. Keep animals for at least 24 hours at room temperature without manipulation. The resin curing time does not differ between P10 and P60 animals.

- CRITICAL STEP: Do not manipulate the sample during the 24 hours. Otherwise, this may influence the curing and may lead to deformations in the vascular corrosion cast.

#### Tissue maceration of soft tissue (day 2-3) ● TIMING ∼1 d

10. Transfer the animal into 300-400 ml (for P60 animals) 7.5% KOH and incubate at 50°C for at least 24 hours. Place only one animal into the solution at one time and make sure to cover the entire body.

#### Decalcification of bony structures (days 3-4) ● TIMING ∼1d

11. Incubate the animal in 300-400 ml (for P60 animals) 5% formic acid at 50°C for at least 24 hours. Place only one animal into the solution at one time and make sure to cover the entire body.

#### Brain dissection (day 4) TIMING ● ∼5 min per mouse

12. Free the cerebral vasculature from remaining extra-cranial vessels using fine scissors and forceps (Figure 2k-n). Optionally, the extracranial vessels can also be further processed and analyzed according to a similar protocol as the one described here, depending on the vessels/organs of interest.

- CRITICAL STEP: Perform the brain vascular corrosion cast dissection/pruning carefully (Figure 2k-n and Supplementary Video 1) because the vessel cast is mechanically sensitive.

#### Wash in distilled water (day 4) TIMING ● ∼5 min per animal

13. Wash the brain vascular corrosion cast in distilled water. Place only one brain at the time into 15 ml of distilled water.

- CRITICAL STEP: Use plenty of distilled water in order to eliminate all residues of tissue and dissected pieces of the extracranial vessels.

#### For quality control, prepare 1-2 brains for Scanning Electron Microscopy (SEM) (day 4) TIMING ● variable

14. For routine SEM, mount a small random sample of the brain casts mounted on stubs and sputter-coat them with gold. The SEM scanning procedure should be performed according to the device manufacturer manual.

#### Osmification with osmium tetroxide (OsO_4_) of brain vascular corrosion cast (day 4) TIMING ● ∼24 hours

15. Transfer the brain vascular corrosion casts not used for SEM into 2% OsO_4_ in 5ml water at room temperature and incubate for 24 hours in order to reach better absorption contrast, and reduce CNR during the SRμCT imaging.

!CAUTION: OsO_4_ is very toxic. Work under a fume hood and use protection gloves, glasses and lab coat. Properly dispose the OsO_4_ waste.

#### Wash in distilled water (days 4-5) TIMING ● ∼5 min per animal

16. Wash the brain vascular corrosion cast thoroughly in distilled water. Place only one brain at the time into 15 ml of distilled water and if the protocol is paused here, keep the cast emerged until the next step, changing the water every 12 hours.

- CRITICAL STEP: Use plenty of distilled water in order to eliminate all residues.

#### Lyophilization by freeze-drying (days 4-5) TIMING ● ∼ 24 hours

17. Dry the brain vascular corrosion casts by freeze-drying in a lyophilization machine overnight at -54°C in a vacuum.

#### Mounting on Plexiglas stubs (days 5-6) TIMING ● ∼5 min per brain resin cast

18. Mount each brain resin cast using cyanoacrylate glue on Plexiglas stubs (size of the stubs is dependent on the µCT set-up and brand) equipped with stainless steel pins for sample repositioning in the μCT.

#### Scanning Electron Microscopy (day 5) TIMING ● ∼15 min per brain resin cast

19. Evaluate all corrosion casts by light microscopy, and then analyze the subset of corrosion casts selected in step 14 with SEM as quality control. The qualitative analysis consists of the presence of a dense vascular network without holes or irregularities and the presence of endothelial cell nuclei imprints on large vessel casts (Extended Data Figure 2).

#### Desktop μCT imaging (day 5) TIMING ● ∼15 min per animal

20. Acquire 3D images of whole-brain samples not used for SEM using a desktop µCT system by selecting the various imaging parameters according to manufacturer’s instructions (or optionally using a SRμCT scanner, see paragraph *Alternatives for wider accessibility*). We used a µCT 40, Scanco Medical AG setup, operated at 50kVp and 160µA with a nominal isotropic voxel size of 16µm. Per scan, we acquired 1000 projection images with an integration time of 200ms and an additional three times frame averaging for improved SNR. The exact settings depend on the desktop μCT system used.

#### Selection of the regions of interest (ROIs) (day 5) TIMING ● ∼15 min per animal

21. Reconstruct, segment and import tomographic images into custom-made or existing ROI picker software. Place one to three ROIs of 1 mm³ per animal and brain region in the cerebral cortex, hippocampus, superior colliculus and brain stem (Figure 4a-l). Then calculate relative coordinates of ROIs with respect to the two-sample holder pins and save them for subsequent re-orientation and measurement at the synchrotron beamline. This step is not needed when the entire brain cast is scanned in high resolution.

#### High-resolution SRμCT imaging of selected ROIs or the whole brain (day 5-6) TIMING ● ∼30 min per animal (∼24 h for one whole brain scan)

22. Acquire high-resolution 3D images of selected ROIs or the whole brain using a previously described local tomography setup. Briefly, the system we used operates at 17.5 keV with a sample-detector distance of 30 mm for distinct edge enhancement, and an optical magnification resulting in an isotropic voxel size of 0.73 µm. Each measurement includes 2000 projections acquired at an integration time of 0.1s each. Perform phase-retrieval with the Paganin method. Reconstruct data using the gridrec algorithm. The net volume of the final images is 2560×2560×2160 voxels, corresponding to 5.5 mm^3^ of tissue volume.

23. When the entire brain vascular corrosion cast is scanned in high resolution (as compared to the high-resolution scanning of pre-selected ROIs), the mosaic images should be stitched together. To construct an artifact-free image mosaic from the sub-images, we used a non-rigid stitching approach, according to^125^. This image mosaic can then be registered to the Allen Mouse Brain Atlas (http://developingmouse.brain-map.org/docs/Overview.pdf) or – alternatively – to any other Mouse Brain Atlas and divided into segments. We divided the anatomical structures of interest into segments with the exact same dimensions as the previously used ROIs (in the ROI-based approach), in order to proceed with similar analysis process.

#### Computational 3D reconstruction of the 3D vascular network – vessel segmentation (day 7-9) TIMING ● ∼15 min per animal

24. OPTIONAL STEP: If the reconstructed CT images are noisy, apply bilateral filtering in order to improve the signal-to-noise ratio. This step is not required if the SNR is high. In order to separate vessels from the background in the reconstructed CT images, use the Otsu thresholding method^126^ (Figure 4m-p).

25. Erase small gaps and holes in vessels by performing morphological closing with a spherical structuring element (with a suggested radius of 1-3 voxels).

26. Perform a hole filling operation that merges isolated background regions into the foreground.

27. Calculate the distance map of the segmented vessels, where each vessel voxel is assigned the distance to the nearest background pixel.

28. Calculate the skeleton of the segmented vessels, where each vessel is reduced into a one-voxel wide centerline (Figure 4o).

29. Trace the centerlines into a graph where graph nodes represent bifurcation points and graph edges represent vessels. For each edge, store a list of locations of voxels that form the edge. Store also the distance map value at each edge voxel as it describes the local radius of the vessel at that pixel (Figure 4o).

30. Use anchored convolution to smooth the voxel list of each edge (suggested sigma = 1.125 µm, maximal displacement = 0.5 µm). Do not allow any displacement in the ends of the edge.

31. Calculate the length of each edge by summing distances between consecutive smoothed edge voxels.

32. Remove all isolated branches whose length is small, suggested threshold length = 5 µm.

33. Remove all branches that have one free end and that satisfy L < 2 max (r_i_,r_2_) where L is the length of the branch, and r_!_ are the radii at the endpoints of the branch.

34. Remove all nodes that have exactly two edges connected to them (these might be created in the previous step) and combine the two branches into one.

35. Remove all nodes that have no edges connected to them.

#### Global vascular network morphometry and local vascular network topological analysis (day 7-9) TIMING ● ∼15 min per animal

36. Categorize vessels and branch points into capillary (diameter < 7µm), non-capillary (diameter ≥ 7µm) and global (without distinction by diameter). Global is the union of capillaries and non-capillaries.

37. Calculate the degree of each branch point. Count the number of branch points per degree, for capillary, non-capillary and global branch points. Use this to plot branch point degree statistics (Extended Data Figures 7–9).

38. Take the above-described values per degree and divide them by network volume. Notice that the tomographic image that the graph is based on is usually cylindrical and the volume of the cylinder should be used. Use this to plot branch point density statistics (Figures 7c,f-h, and Extended Data Figure 5,6). The branch point density per volume is defined as the number of branch points per cubic millimeter.

39. Use the branch point diameter, classified into global, capillary, and non-capillary, to plot branch point diameter statistics (Figure 7c, and Extended Data Figures 5b,6b).

40. Classify segments into global, capillary, and non-capillary. Count all occurrences per classification. Divide by network volume to get the segment density. Use this to plot segment density statistics (Figure 7c,f-h, and Supplementary Figure 5–9).

41. Take segment diameter, and -length per classification to plot segment diameter, and -length statistics (Figure 8c,g-n).

42. Use the values from step 41 and relate length to diameter to plot their relation (Figure 8c).

43. Use the values from step 41 and calculate the volume of the segment as the sum of volumes of cylinders corresponding to line segments in that segment. Use this to plot segment volume statistics (Figure 8t-w).

44. Calculate tortuosity of each segment as the relation between the real length of the segment and the shortest distance of this segment’s first and last point. Use this to plot the tortuosity statistics (Figure 8p-s).

45. Use the image containing the segmented vessels and calculate the Euclidean distance transform of non-vessel regions. This creates a volume containing the distance to the closest vessel for each non-vessel voxel. Create a histogram of the distance values to estimate the extravascular distance distribution (Figure 9b,i-k, and Supplementary Figure 5).

46. Count the number of background and vessel voxels. Calculate vascular volume fraction (volume fraction of all vessels) as the ratio between number of vessel voxels and total number of voxels. (Figure 5h-j, Figure 6l-q,t-v).

47. Calculate the sum of all segment volumes for global, capillary, and non-capillary segments. Divide the capillary volume by the global volume. Use this value to scale the volume fraction calculated in step 46 to get the approximate capillary volume fraction. Do the same for non-capillaries. Use these values to plot the capillary and non-capillary vascular volume fraction statistics (Figures 6l-q,t-v).

48. Derive the vessel directionality by attributing every vessel to its main direction (x, y, z) in the 3D space within the selected ROI. For each line of each segment, normalize the vector that represents the line. To quantify the directions of vessels emanating from each branchpoint, calculate the relative angles between all segments that join at a branch point. This allows for a quantitative representation of how orthogonal (90 degrees) or straight (0 and 180 degree ranges) vessels go in and out of branchpoints. This normalized direction (x,y,z) is directly mapped to “redness” (r), “blueness (b), and “greenness” (g) (r,g,b), and then scaled by 1.75 to get a better color saturation and clamped to [0,1]. Render the computational 3D reconstructions with color mapped to 1) directionality, 2) vessel radius, and 3) classification (capillary or non-capillary) of each vessel (Figure 10, and Supplementary Figures 6–9).

#### Statistical analysis

Compare the mean of the parameters of each ROI obtained from the different sample groups using the Welch t-test (statistically significant differences are defined as P-values below 0.05). Use Matplotlib (https://matplotlib.org) and NumPy (https://numpy.org) or comparable software to visualize the data with histograms and boxplots. For the whole-brain scan method, the mean of the parameters obtained from one animal in the different groups (P10 WT, P60 WT) were also compared using the Welch t-test. Histograms and boxplot to visualize the data (Figures 5–9, Extended Data Figures 5–9, and Supplementary Figures 1–5,10–14), as well as statistical testing and figure plotting were performed in the exact same manner as described for the ROI-based method.

## ANTICIPATED RESULTS

With the protocol described here, perfused blood vessels of the total vascular network can be quantitatively assessed in any desired mouse brain region, starting from the postnatal (at least as early as P10) to the adult stage by analyzing various vessel parameters such as vascular volume fraction, branch point density and degree, segment density, -diameter, -length, - tortuosity, -volume, -directionality, and the extravascular distance. The vascular volume fraction, the vessel segment- and branch point density in various brain regions including cortex, hippocampus, and superior colliculus were significantly increased in the adult (P60) as compared to the postnatal (P10) mouse brain (Figure 5h-j, Figure 6l,o,t, Figure 7f, and Extended Data Figures 5e,6e,7e), indicating active sprouting angiogenesis and vascular network formation from the developing postnatal- to the mature adult stage of mouse brain development. Vascular corrosion casting can be combined with immunofluorescent or other staining which may ease the navigation within the vascular network and allows direct correlation with molecules or cell structures of interest (Extended Data Figure 10), e.g. perivascular cells of the neurovascular unit such as pericytes, astrocytes, or neurons. We recommend using additional labelling with other markers such as DAPI to label cell nuclei when using markers for any other cell or protein of interest (Extended Data Figure 10).

The 3D vasculature of mice during postnatal brain development was well visible in the polyurethane resin PU4ii-perfused P10 WT and P60 WT mice (Extended Data Figure 2). Light microscopy and SEM of brain tissue samples from resin-perfused P10 and P60 WT mice revealed a dense vascular network in the superficial cortex with well-defined vessel structures of all sizes (Extended Data Figure 2). During resin perfusion, endothelial cell nuclei are exposed to the passing resin material, resulting in imprints of blood vessels on the cast surface (Extended Data Figure 2f), which were observed in vascular corrosion casts of both P10 and P60 mice, demonstrating the perfect molding of the vascular lumen^110^. Moreover, the fine capillary network was well defined and visible and devoid of vessel interruptions in both the P10 and the P60 brain cortices (Extended Data Figure 2), and all these morphological features were in line with the characteristics of the normal cerebral vasculature in postnatal adult mice that we and others have previously published^40,104,109^. The raw images of the SRμCT of the vascular corrosion casts, that were the basis for subsequent computer-aided 3D-reconstruction (Figure 5a), showed excellent quality (Figure 4m,n, and Figure 5a-d). The vascular corrosion casting technique is, therefore, an excellent method to visualize the 3D vessel network architecture in the brains of both the P60 and also the developing P10 mice.

Here, we demonstrated that with minor adaptations vascular corrosion casting can indeed be applied in very young and small mice at the stage of postnatal brain development at which vascular sprouting is highly active^2,3,23,38,39,76^.

Functional, perfused blood vessels that constitute the 3D brain vascular network can be best visualized with a marker that is perfused into the circulation of an intact mouse. For that reason, we used polyurethane resin PU4ii that labels the inner vascular lumen of actively, *in vivo*– perfused blood vessels (Figure 2k-n, and Figure 3). Thereby, resin PU4ii-perfused blood vessels and in consequence, the entire 3D brain vascular network can be imaged and computationally analyzed in both postnatal and adult mouse brains (Figures 5–10), allowing to address the development of the vascular network between two developmental stages.

Perfused blood vessels were hierarchically imaged by SRµCT and subsequently computationally analyzed and quantified by an image analysis pipeline (Figure 4m-p, Figure 5a-d, and Figure 6a-d). Various vessel parameters (see above) of PU4ii resin-perfused vessels, as well as the extravascular distance, can be quantified using computational analysis consisting of either global vascular morphometry analysis or local topological vascular network analysis^40,121,126,130–132^.

Global vascular network morphometry – which allows quantification of properties of each dataset without considering the form- and shape properties of individual vascular structures, generating an abstract quantification of the data (Figures 5h-j, Figure 6l-q,t-v, and Figure 7f-h,8c-s) - revealed that in the P10 WT animals, between 1.67% in the cortex, 1.93% in the superior colliculus, and 2.14% in the hippocampus of the total volume of the postnatal mouse brain tissue was occupied by perfused vessel structures, and this percentage was between 2.58% in the cortex, 2.81% in the superior colliculus, and 2.97% in the hippocampus of the total volume of the adult mouse brain tissue, data which are in accordance with previous findings by our^27,40,108^ and other groups^21,24^. In the P60 WT mouse brain, the vascular volume fraction was significantly increased as compared with P10 WT animals in the cortex (P60 WT vs. P10 WT (mean ±SD; applies also for all subsequent values): 2.58 ±0.77% vs. 1.67 ±0.45%, P = 0.0032), hippocampus (P60 WT vs. P10 WT: 2.97 ±0.70% vs. 2.14 ±0.22%, P = 0.019), and superior colliculus (P60 WT vs. P10 WT: 2.81 ±1.15% vs. 1.93 ±0.26%, P = 0.048) (Figure 5h-j). The increase of the vascular volume fraction of perfused vessels between the two developmental stages P10 and P60 was between 39% and 54% (+54% in cortex, +39% in hippocampus, +45% in superior colliculus), in line what we observed previously using stereological methods^39^. However, we are not aware of any previous study that compared postnatal and adult mouse brain vasculatures at the network level.

To illustrate the sensitivity and broad applicability of the proposed method, we used a knockout mouse model of the well-known anti-angiogenic molecule Nogo-A (Nogo-A^−/−^)^27,39,40,170–172^, at both the developing P10 as well as at the P60 stages. Interestingly, the increase in vascular volume fraction in the P10 Nogo-A^−/−^ as compared to the P10 WT was similar to the comparison between the P60 WT and the P10 WT mice in all brain regions, whereas these effects were no longer visible between P60 Nogo-A^−/−^ and P60 WT mice (Supplementary Figure 2), indicating i) a hypervascularization/accelerated vascularization in the Nogo-A^−/−^ mice during postnatal brain development, and ii) that these effects are compensated/strongly attenuated in the adulthood^39^, in agreement with what we previously showed^39,40^. Given the redundancy of mechanisms acting on the vascular system, effects of genetic deletion of angiogenic/anti-angiogenic molecules on angiogenesis and the vasculature network that are observed during embryonic or postnatal development are often compensated at the adult stage, and this is also the case for the Nogo-A^−/−^ mouse^39,40^ (Supplementary Figures 2–5).

In contrast to global vascular morphometry, local vascular network topology analyzes each individual vascular structure and bifurcation point separately, thereby allowing to quantify the form and shape of the vascular network (Figure 6a-d). The vessel-specific properties of each individual vascular structure (= each vessel segment) and each bifurcation point within the dataset thereby affects the quantification and analysis of the 3D vascular network (Figure 6) and local topological analysis of the 3D vascular network allows the characterization of individual per-vessel parameters like segment density, -diameter, -length, -tortuosity, -volume, and branch point density and -degree (Figures 6–8). Importantly, local topological analysis allows to differentiate between capillary (< 7 µm in diameter) and non-capillary vessels (≥ 7 µm in diameter) of the highly interwoven hierarchical 3D vessel/capillary network (Figure 6)^173,174^.

For quantitative assessment of angiogenic features such as branch point density and -degree, segment density, -diameter, -length, -tortuosity, -volume and vascular branch points, we therefore referred to local topological vessel analysis provided by a computational image analysis pipeline (Figures 6–9, PROCEDURE steps 24-48). Examples of the quantification of the aforementioned vessel parameters in different regions of the postnatal- and adult mouse brains are shown in Figures 5h-j, 6e-g,l-q,t-v, 7c,f-h, 8b,e-j, 9c-s, 10b,i-k, and Extended Data Figures 5b,e-g, 6b,e-g, 7b,e-j, 8b,e-j. The absolute values for the vascular volume fraction are in accordance with previous findings by our group using immunofluorescent staining and stereological analysis^27,39^ and with studies using similar^104,108^ or different 3D methods^21,175^.

Based on the distinction between capillaries and non-capillaries, we reveal a very important significant increase of the density of capillary blood vessels (as revealed by the vascular volume fraction) between the developing postnatal- and the mature adult mouse brain vascular network (Figure 6n,q,v), whereas the non-capillary blood vessels showed only a non-significant increase between these two time points, in various brain regions (Figure 6m,p,u). Local vascular network topology revealed that the 3D vascular volume fraction in the P60 WT cortex was increased by 54% as compared to the P10 WT cortex (Figure 6l), and by 39% and 45% in the hippocampus (Figure 6o) and superior colliculus (Figure 6t), respectively, and these values were all similar to the fold change calculated by global vascular morphometry (Figure 5h,i,j). Deeper insight into the 3D vascular network architecture is provided by addressing specific properties of each individual vessel structure, for instance by addressing the vessel diameter (Figure 6).

Accordingly, of the total vessel tissue volume (= vascular volume fraction) of 1.67% in the P10 cortex, 1.24% (74,2% of the total vessel volume) was occupied by non-capillary vessels (Figure 6m) and 0.43% (25.8% of the total vessel volume) by capillaries (Figure 6n) whereas these values were 1.35% for non-capillaries (62.9% of the total vessel volume) (Figure 6p), 0.79% for capillaries (37.1% of the total vessel volume) (Figure 6q) in the P10 hippocampus, and 1.3% for non-capillaries (33% of the total vessel volume) (Figure 6u) and 0.64% for capillaries (67% of the total vessel volume) (Figure 6v) in the P10 superior colliculus, respectively.

Adult (P60) WT animals showed an increased absolute frequency of capillaries as compared to P10 WT animals throughout the different brain regions, whereas the absolute frequency of bigger vessels was very similar between the two developmental stages (Figure 6e-g). Computational 3D analysis revealed that the increased vascular volume fraction in the cortices, hippocampi, and superior colliculi of P60 WT animals was mainly due to a significant increase of the vascular volume fraction at the capillary level (cortex: P60 WT vs. P10 WT: 1.16 ±0.37% vs. 0.43 ±0.22, P = 5.8 × 10^−6^; hippocampus: P60 WT vs. P10 WT: 1.28 ±0.37% vs. 0.79 ±0.1, P = 0.011; superior colliculus: P60 WT vs. P10 WT: 1.52 ±0.42% vs. 0.64 ±0.14, P = 4.1 × 10^−6^, whereas a slight increase of the vascular volume fraction of larger vessels (cortex: non-capillaries, P60 WT vs. P10 WT: 1.42 ±0.56% vs. 1.24 ±0.37%, P = 0.41; hippocampus: non-capillaries, P60 WT vs. P10 WT: 1.7 ±0.57% vs. 1.35 ±0.17%, P = 0.20; superior colliculus: non-capillaries, P60 WT vs. P10 WT: 1.29 ±0.83% vs. 1.3 ±0.2%, P = 0.99) did not reach statistical significance (Figure 6e-g,l-q,t-v). Quantitatively, the vascular volume fraction was increased by 169% for capillaries and by 14% for non-capillaries in the cortex, by 60% for capillaries and by 26% for non-capillaries in the hippocampus, and by 138% for capillaries and by 0% for non-capillaries in the superior colliculus (Figure 6e-g,l-q,t-v). Taken together, these results indicate that the increased vascular volume fraction observed from the postnatal to the adult stage is mainly due to additionally formed capillaries whilst the number of larger non-capillaries remains relatively unaffected.

The observed increased vascular volume fraction may be due to increased vessel branching and concomitant newly formed vessel segments or caused by an increased vessel diameter, by an increased vessel length, or a combination of these factors. To address the origin of the additionally formed vessels, the number of vascular branch points in the cortices of P10 WT and P60 WT animals referring to local topological analysis was determined (Figure 7). Vessel branch points are characterized by their degree^108^, defined as the number of vessels adjacent to a vessel branch point, usually ranging from degree 3 to degree 5^176^ (Figure 7b). Accordingly, vascular branch points were defined as points where three (branch point degree 3), four (branch point degree 4), or five (branch point degree 5) vessel segments converge (Figure 7b). For discrimination between “capillary branch points” and “non-capillary branch points,” the branch point diameter was considered to be that of the thickest vessel adjacent to a given branch point (Figure 7b). Accordingly, if the thickest vessel adjacent to a given branch point was <7 μm, the branch point was classified as a “capillary branch point,” whereas if the thickest vessel adjacent to a given branch point was ≥7 μm, the branch point was classified as a “non-capillary branch point.” Adult (P60) WT animals showed a pronounced increase of branch point density (of all degrees) at the capillary level (P60 WT vs. P10 WT: 13994.1 ±7395.6mm^−3^ vs. 1929.1 ±819.1mm^−3^, P = 5.3 × 10^−5^) and a slighter increase of the number of branch points for non-capillaries (P60 WT vs. P10 WT: 2959.9 ±1559.7mm^−3^ vs. 1993.3 ±683.7mm^−3^, P = 0.10 as compared to the P10 WT mice in the cortex, as well as in the hippocampus, and superior colliculus (Figure 7c,f-h and Extended Data Figure 5b,e-g). Computational reconstructions revealed an increased number of vessel branch points in cortices, hippocampus, and superior colliculi of P60 WT animals as compared with P10 WT mice (Figure 7d,e and Extended Data Figure 5c,d). Quantitatively, the density of branch points for all vessels in the different brain regions cortex, hippocampus, and superior colliculus was significantly – by 332%, 230%, 435%, respectively – increased (cortex: P60 WT vs. P10 WT: 16954.0 ±8329.6mm^−3^ vs. 3922.4 ±1376.9mm^−3^, P = 9.6 × 10^−5^; hippocampus: P60 WT vs. P10 WT: 19814.4 ±6726.6mm^−3^ vs. 6010.0 ±1098.8mm^−3^, P = 0.00019; superior colliculus: P60 WT vs. P10 WT: 23907.3 ±11030.9mm^−3^ vs. 4471.1 ±1581.4mm^−3^, P =4.0 × 10^−5^) in adult (P60) WT as compared to postnatal (P10) WT animals (Figure 7d,f-h and Extended Data Figure 5e-g), predominantly due to a more than seven fold increased (+625%) number of branch points in the capillary bed in the cortex, a more than four times increased number of branch points in the capillary bed in the hippocampus (+333%), and a more than eight times increased (+728%) number of branch points in the capillary bed in the superior colliculus (cortex: P60 WT vs. P10 WT: 13994.1 ±7395.6mm^−3^ vs. 1929.1 ±819.1mm^−3^, P =5.3 × 10^−5^; hippocampus: P60 WT vs. P10 WT: 15257.6 ±5269.4mm^−3^ vs. 3523.7 ±691.1mm^−3^, P = 0.022; superior colliculus: P60 WT vs. P10 WT: 19990.9 ±8594.8mm^−3^ vs. 2415.8 ±1575.1mm^−3^, P = 4.7 × 10^−5^ (Figure 7d,f-h and Extended Data Figure 5e-g) and an additional, smaller increase (+48% in the cortex, +78% in the hippocampus, +91% in the superior colliculus) at the level of non-capillaries (cortex: P60 WT vs. P10 WT: 2959.9 ±1559.7mm^−3^ vs. 1993.3 ±683.7mm^−3^, P = 0.10; hippocampus: P60 WT vs. P10 WT: 4556.8 ±1974.1mm^−3^ vs. 2486.3 ±471mm^−3^, P = 0.035; superior colliculus: P60 WT vs. P10 WT: 3916.4 ±2911mm^−3^ vs. 2055.3 ±307.6mm^−3^, P = 0.091, (Figure 7d,f-h and Extended Data Figure 5e-g). Interestingly, in the adult animals, this was mainly due to an increased number of vessel branch points of degree 4 for all vessel sizes (cortex: +839% for all vessels, P60 WT vs. P10 WT: 1769.4 ±1047.0mm^−3^ vs. 188.4 ±62.2mm^−3^, P = 0.00015, +182% for non-capillaries, and +1345% for capillaries; hippocampus: +648% for all vessels, +345% for non-capillaries, and +792% for capillaries; superior colliculus: +1313% for all vessels, +552% for non-capillaries, and +1631% for capillaries (Extended Data Figures 7h-j,8h-j,9h-j, and Supplementary Figure 1e,f,i), while showing a highly significant increase also for vessel branch points of degree 3 for all vessel sizes (cortex: +304% for all vessels, +42% for non-capillaries, and +578% for capillaries; hippocampus: +206% for all vessels, +71% for non-capillaries, and +303% for capillaries; superior colliculus: +392% for all vessels, +76% for non-capillaries, and +670% for capillaries (Extended Data Figures 7e-g,8e-g,9e-g, and Supplementary Figure 1e,f,i), indicating both formation of new vessel branch points (degree 3) and further branching of existing degree 3 branch points (degree 4). Vessel branch points of degree 5 were not detected in either study group (Extended Data Figures 7b,8b,9b). Moreover, the mean vessel branch point degree for the different vessel sizes showed an overall increase between the two developmental stages (cortex: +2% for all vessels, +1% for non-capillaries, and +2% for capillaries; hippocampus: +2% for all vessels, +2% for non-capillaries, and +2% for capillaries; superior colliculus: +3% for all vessels, +2% for non-capillaries, and +3% for capillaries (Extended Data Figure 7b,8b,9b), indicating that branch point degree increases during physiological vascular brain development. Together, these data reveal that the enhanced vascular volume density observed from the postnatal to the adult stage can be mainly explained by an increased number of vascular branch points mainly at the level of capillaries.

Interestingly, we observed a significant increase (+39%** for all vessels) for the vascular volume fraction (again mainly at the level of capillaries, +71%**) when comparing data on the cortex of P10 Nogo-A^−/−^ with P10 WT mice, and these effects were reminiscent of the comparison between P10 WT and P60 WT mice mentioned above (Supplementary Figure 2). At the adult stage, however, the effect of genetic deletion of Nogo-A did not show significant differences in the brain vasculature between the WT and Nogo-A^−/−^ groups, suggesting that the inhibitory effect of Nogo-A on sprouting angiogenesis is compensated by the redundancy of mechanisms acting on the vascular system during these later stages of cerebrovascular development^39^.

Both the effects of the developmental stage (e.g. adult P60 versus developing P10) as well as the effect of deletion of the Nogo-A gene at the postnatal (but not adult) stage were consistent throughout various brain regions including cortex (P10 Nogo-A^−/−^ vs. P10 WT 2.32 ±0.35% vs. 1.67 ±0.45%, P = 0.0024), hippocampus (P10 Nogo-A^−/−^ vs. P10 WT 2.92 ±0.56% vs. 2.14 ±0.22%, P = 0.027), and superior colliculus (P10 Nogo-A^−/−^ vs. P10 WT 2.33 ±0.37% vs. 1.93 ±0.26%, P = 0.037) (Supplementary Figure 2).

In addition to these measured values, parameters such as the average segment density (Figure 8d-f, and Supplementary Figure 4a-c), -diameter (Figure 8g-j, and Supplementary Figure 4d-f), -length (Figure 8k-n, and Supplementary Figure 4g-i), -tortuosity (Figure 8p-s, and Supplementary Figure 4j-l) and the average maximal perfusion distance were derived (Figure 9, and Supplementary Figure 5). Given the high angiogenic activity in the postnatal rodent brain^27,38^, the precise values of the aforementioned parameters will obviously depend on the exact developmental stage of the mouse.

The functional consequences of the increased vascular volume fraction in the P60 (and the Nogo-A knock-out animals), can be addressed by determining the extravascular distance – defined as the shortest distance of any given extravascular voxel in the tissue to the nearest vessel structure (Figure 9, and Supplementary Figure 5) – which measures the diffusion distance. The extravascular distance in P10 WT cortices was 31.9µm on average and thus similar to our previous work^40,176^, whereas the P10 WT hippocampus and P10 WT superior colliculus showed lower average values of 18.4 µm and 19.2 µm, respectively (Figure 9i-k, and Supplementary Figure 5). Overall, the extravascular distance was markedly reduced in P60 WT mice as compared to P10 WT cortices, hippocampi, and superior colliculi (Figure 9i-k, and Supplementary Figure 5). Accordingly, histogram distribution analysis revealed a marked shift towards smaller extravascular distances in the P60 animals (Figure 9i-k, and Supplementary Figure 5), which is a consequence of the higher vascular volume fractions observed in the P60 as compared to the P10 WT animals. Quantitative analysis showed a highly significant reduction of the extravascular distance in P60 WT animals to almost half (−45%) as compared to P10 WT mice in the cortex, (P60 WT vs. P10 WT: 17.42 ±6.52 μm vs. 31.94 ±10.63 μm, P =1.6 × 10^−5^, Figure 9i), hippocampus (P60 WT vs. P10 WT: 13.4 ±3.93 μm vs. 18.44 ±2.17 μm, P = 0.014, Figure 9j), and superior colliculus (P60 WT vs. P10 WT: 11.15 ±3.55 μm vs. 19.22 ±4.29 μm, P =1.7 × 10^−5^, Figure 9k). This absolute value of 17.42μm (P60 WT) is comparable with a study by Tsai et al., which, using automated histological imaging and analysis tools simultaneously mapping the locations of all neuronal and nonneuronal nuclei and centerlines of the entire vasculature within slabs of mice adult mice neocortex, found that each individual neuron is no more than 15 μm away from the closest microcapillary, thereby assuring proper oxygen delivery^21^.

In line with the findings presented above, the extravascular distance was also reduced in the P10 Nogo-A^−/−^ animals as compared to the P10 WT animals in the different regions, and the effects observed were similar to the comparison between P60 WT mice and P10 WT mice: cortex (P10 Nogo-A^−/−^ vs. P10 WT: 16.56 ±2.30 μm vs. 31.94 ±10.63 μm, P = 0.00013, Supplementary Figure 5), hippocampus (P10 Nogo-A^−/−^ vs. P10 WT: 16.34 ±0.93 μm vs. 18.44 ±2.17 μm, P = 0.085, Supplementary Figure 5), superior colliculus (P10 Nogo-A^−/−^ vs. P10 WT: 16.58 ±3.06 μm vs. 19.22 ±4.29 μm, P = 0.41, Supplementary Figure 5).

Together, these data suggest that as the 3D vascular network undergoes significant changes (e.g. increased vascular volume fraction and increased vessel branching) from the developing P10 to the P60 mature stage, these results in a markedly lowered vessel spacing and diffusion distance throughout various brain regions.

Further, we derived the parameter vessel directionality by determining for every vessel segment to its main angle to x, y, and z in the 3D space within the selected ROIs (Figure 10) and observed different patterns of vessel directionality throughout the different brain regions examined:

i. in the P10 cortex, the superficial PNVP extended in the x- and y-directions, whereas the INVP exhibited a radial sprouting pattern into the brain parenchyma along the z-axis, perpendicular to the PNVP (Figures 10b,c), in accordance with literature27,32,145. The distinction between non-capillaries and capillaries revealed that the PNVP and INVP were mainly composed of non-capillary vessels (Supplementary Figure 6), and that non-capillaries also formed the base of vessel sprouts emanating from the INVP (and to a lesser extent from the PNVP) mainly in the x- and y-directions (Supplementary Figure 6). Capillaries, on the other hand, vascularized the cortical CNS parenchyma via sprouting from non-capillaries and extended equally in all directions (x-,y-,z-), thereby assuring adequate micro vascularization and perfusion of the brain (Supplementary Figure 6). At the adult stage, these patterns of vessel directionality observed at the postnatal developing stage were confirmed and further strengthened (Figure 10, and Supplementary Figure 6).
ii. in the P10 hippocampus, horizontally orientated main vessel branches in the x-axis were recognizable, most likely following the distinctive curved shape of this anatomical structure (Figure 10d,e, and Supplementary Figure 7). Differentiation between non-capillary and capillary vessel structures indicated that these main vessel branches were mainly composed of non-capillary vessels (Figure 10d,e, and Supplementary Figure 7). Similar to what we observed in the cortex, these non-capillaries also formed the base of vessel sprouts emanating from these main branches, mainly in the x- and y-directions (Figure 10, and Supplementary Figure 7). Capillaries mostly formed the smaller side branches sprouting into the four subregions of the hippocampus proper (CA1, CA2, CA3, dentate gyrus), similar to the human hippocampal vascularization pattern^177,178^ (Supplementary Figure 7).
iii. in the P10 superior colliculus, a comparable pattern of directionality as in the cortex was observed with a recognizable PNVP extending in the x- and y-directions, combined with the INVP vessels radially sprouting into the brain parenchyma along the z-axis (Figure 10f,g, and Supplementary Figure 8).

Quantitative analysis of the relative angles of the vascular segments emanating from a certain branch point revealed that the capillaries in the cortex, hippocampus, and superior colliculus derived at angles between around 75° and 150° at both the developing P10 and mature P60 stages (Figure 10h-j). The non-capillaries, however, showed two main angles of orientation, namely around 90° and at around 170° in three brain regions examined (Figure 10h-j). Interestingly, these two main angles of orientation were more pronounced at the adult P60 as compared to the P10 stage (Figure 10h-j), indicating some sort of preferred vessel orientation of the larger vessels (non-capillaries) that is established throughout vascular brain development. Strikingly, these results were again very similar in the three brain regions, suggesting a more general underlying concept of vessel directionality/orientation. Even though we do not fully understand these data at present, one might think of these preferred orientations of non-capillaries as predefined vascular “routes” into specific brain structures that then terminate into the capillary beds which are more randomly orientated.

Comparison of the quantification of individual vascular parameters obtained by scanning the entire mouse brain in high resolution with that of the two-step ROI-based approach during which the ROIs first have to be placed on a lower resolution overview μCT scan with subsequent high-resolution SRμCT scanning of selected ROIs, revealed very similar results. The exact values might vary in different experimental settings and thus correlate with the angiogenic activity of the specificity of investigated tissue^78^. Nevertheless, we feel that the selected parameters, obtained by our improved protocol might be helpful readouts for better understanding angiogenesis in various biological settings.

Concluding, 3D computational reconstructions of corrosion casts are a powerful tool to investigate any biologically relevant question regarding angiogenesis (genetic mechanisms in physiology and pathophysiology as well as the effect of drugs) because it allows quantification of the morphological features of entire vascular networks. Although corrosion casts represent only a snapshot they can for example also serve as templates for blood flow simulations that can, in addition to simulating cerebral blood flow in healthy brains, be used to simulate impaired oxygen and nutrient supply as a consequence of altered network architecture in different cerebrovascular pathologies, such as brain tumors, brain vascular malformations, and stroke. This protocol of vascular corrosion casting techniques combined with SEM, with or without SRµCT imaging, and computational network analysis was used to demonstrate quantitatively the morphological changes occurring in the mouse brain between birth and adult stage of vascular brain development. In addition, we further illustrated its sensitivity by using a genetic knockout model. We provide a methodology to visualize and analyze 3D vascular networks in the mouse brain at any developmental stage, that could after validation be used also in other organs.

**Table 1:** Troubleshooting

| Step | Problem | Possible reasons | Possible solution |
| --- | --- | --- | --- |
| 6-8 | Air bubbles in the ACSF containing heparin, in the 4% PFA, or in the resin solutions that can kill the animal (due to air embolism) instantly, lead to incomplete perfusion and/or to rupture of the vessel wall. | Air was trapped in the syringe while filling it with ACSF containing heparin, the 4% PFA, or the resin. Air bubbles are difficult to see in the ACSF containing heparin, in the 4% PFA, or the resin solutions | Look under the light, very precisely and remove any air bubble from the syringe. Tilt syringe forward and let air bubble accumulate at the stopper. Stop injection before air enters tubing. |
| 8 | Animal does not become immediately bluish. | Resin leaks out or flows retrogradely into the lung due to mitral valve damage. | Check for leakage between needle and ventricle wall. If leakage is detected, stop infusion and try sealing the leakage using instant glue. In case of retrograde perfusion of the lung exclude animal. Puncture left ventricle very carefully. |
| 8 | Animal does not become immediately blue, and the ACSF containing Heparin, the 4% PFA, or the resin solution comes out of the nose (the respective solution went into the lungs because of damage of lung blood vessels due to retrograde perfusion caused by injury to the mitral valve during needle puncture of the left ventricle). | Injury of the mitral valve due to needle puncture while puncturing the left ventricle. | Low chance to obtain complete perfusion of tissue and organs. Animal needs to be excluded.<br>Inject/infuse a new animal.<br>Puncture the left ventricle very carefully and inject/infuse very carefully. |

## ACKNOWLEDGMENTS

We thank Katrien De Bock, Marc Schwab, Moheb Ghobrial, and Jing Zhang for help with the animal perfusions, Sebastian Eichelbaum for help with the computational analysis and 3D visualizations, and Nancy Chu Ji for help with the illustrations. T.W. was supported by the OPO Foundation, the Swiss Cancer Research foundation (KFS-3880-02-2016-R, KFS-4758-02-2019-R), the Stiftung zur Krebsbekämpfung, the Kurt und Senta Herrmann Foundation, Forschungskredit of the University of Zurich, the Zurich Cancer League, the Theodor und Ida Herzog Egli Foundation, the Novartis Foundation for Medical-Biological Research, and the HOPE Foundation. P.C. was supported by long-term structural Methusalem funding by the Flemish Government (14/08), and a European Research Council (ERC) Advanced Research Grant (EU-ERC269073). J.V. was supported by the Swiss National Science Foundation (no. 310000 120321/1). All the animal experiments were conducted in J.V.’s laboratory and were approved by the Veterinary office of the Canton of Zurich.

## AUTHOR CONTRIBUTIONS

T.W. had the idea for the study, designed the experiments, wrote the manuscript, and made the figures. T.W., A.U.S., E.P.M., and C.H. conducted the experiments. T.W., J.V., A.M. and J.B. analyzed the data. T.W. wrote the manuscript, J.B. helped editing the manuscript and figures. R.W. helped with the animal experiments and gave critical inputs to the manuscript. A.M., A.U.S., T.K., K.D.B., P.C., J.V:, and M.S. gave critical inputs to the manuscript. All authors read and approved the final version of the manuscript.

## EXTENDED DATA FIGURES

**Extended Data Figure 1.**
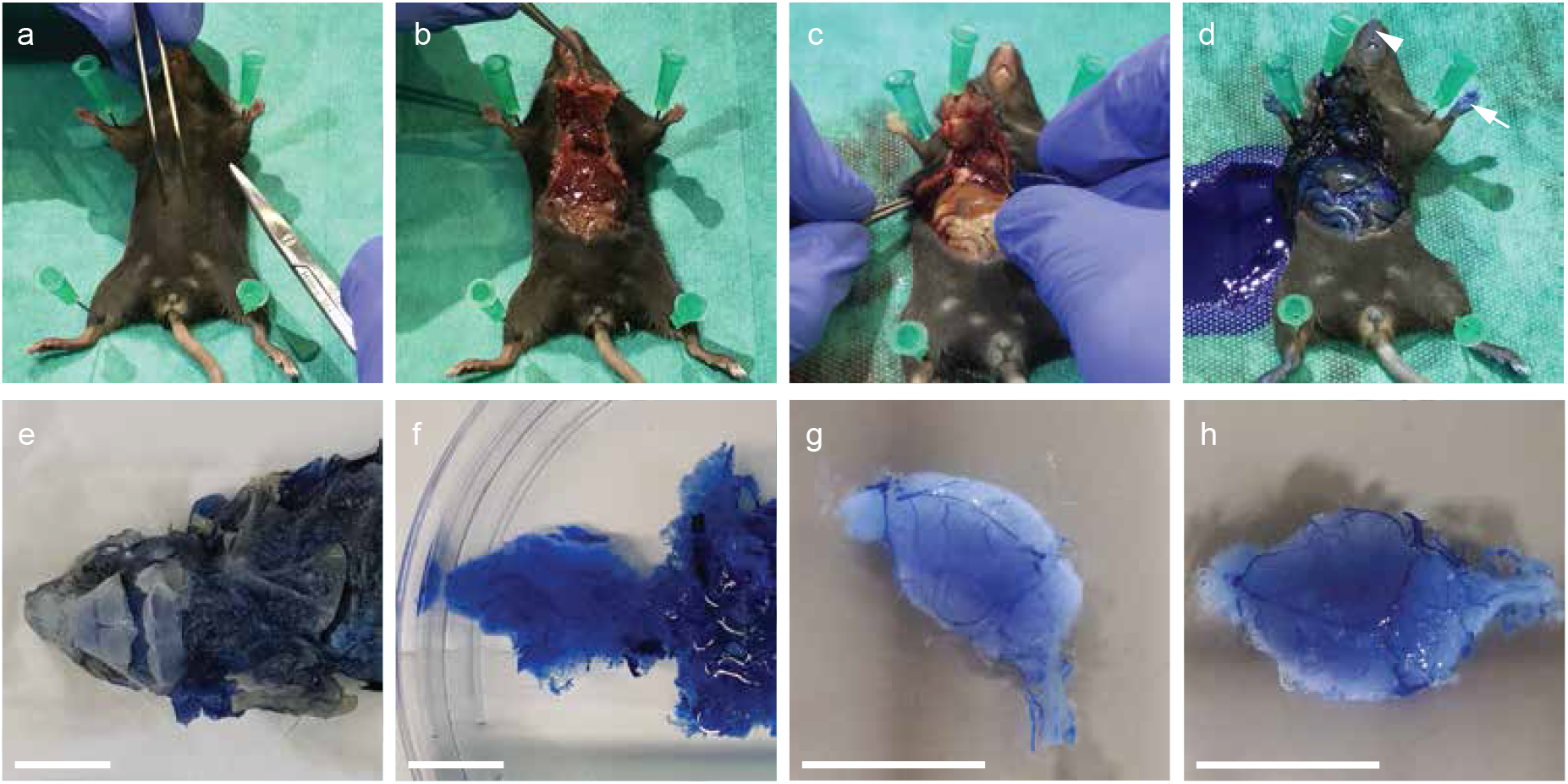
Intracardial resin perfusion and brain dissection for adult (P60) animals. The main steps of resin perfusion and brain dissection are shown as photographs for the perfusion and brain dissection of adult (P60) mice. **a-c** Fixation (**a)**, surgical opening (**b,c**) and site of needle insertion (**c**) for intracardial perfusion (via the left ventricle) with 10–20 ml artificial cerebrospinal fluid containing 25,000 U/L Heparin, followed by 4% paraformaldehyde in phosphate buffered saline, and then by the polymer resin PU4ii (vasQtec, Zürich, Switzerland), all infused at the same rate (8 ml/min and 100-120 mmHg) in an adult (P60) mouse (see also Supplementary Video 1). **d** Successful perfusion is indicated by a bluish aspect of the animal, which can be easiest observed at the paws (white arrow) or the snout (white arrowhead). **e,f** Resin cast of a P60 animal after soft tissue maceration, and before (**e**) and after (**f**) decalcification of bone structures. **f-h** The cerebral vasculature is sharply dissected from the extracranial vessels resulting in the isolated brain vascular corrosion cast of the P60 mouse brain (**g,h**). Scale bars, 10 mm (**e-h**).

**Extended Data Figure 2.**
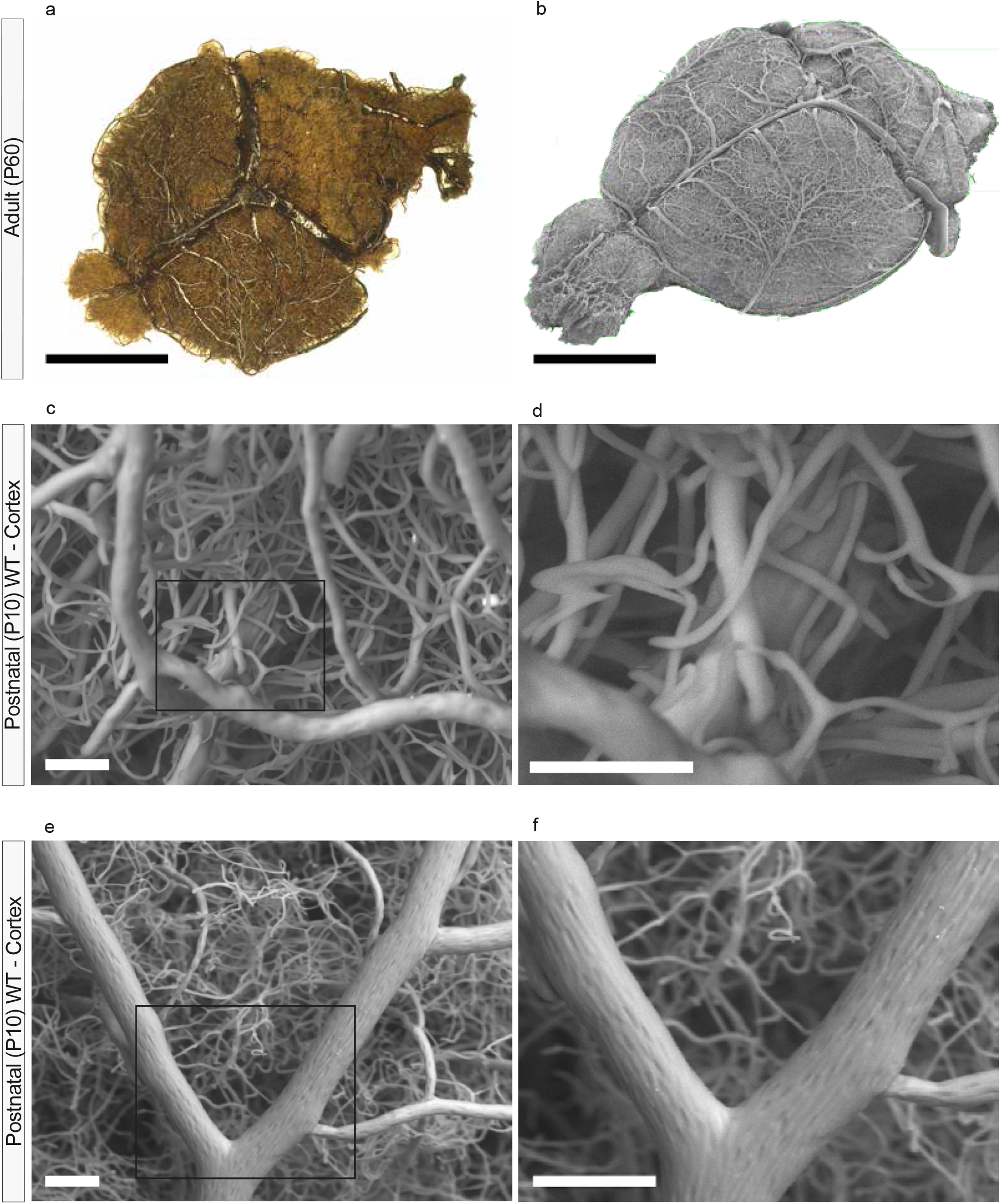
Validation of vascular corrosion casting in postnatal (P10) and adult (P60) mice using light microscopy (LM) and scanning electron microscopy (SEM) **a-f** Quality control of the vascular corrosion casts of P10 and P60 WT mice by visual inspection by LM (**a**), and by SEM (**b-f**). **a** Light microscopy image of the entire brain vasculature of the P60 WT mice. **b** SEM image of the entire brain vascular corrosion cast illustrating the dense vascular network including blood vessels of all sizes with recognizable vascular anatomy. **c-f** The entire brain was uniformly filled with resin and the lumen of the vessels was molded with its finest details. **e** The capillary network in the cortex is well defined and devoid of interruptions. **f** Note the elongated cellular and nuclear imprints on the large vessel that are also typical for good-quality vascular corrosion casts in adult mice^110^. Scale bars: 4 mm (**a**); 4 mm (**b**); 50 μm (**c**); 50 μm (**d**), 100 μm (**e**), 100 μm (**f**) *Reproduced with permission from:* **a,d-f** Wälchli et al. Nogo-A Regulates Vascular Network Architecture in the Postnatal Brain. *J Cereb Blood Flow Metab* 2017 Feb;37(2):614-631 **b** Krucker et al. New polyurethane-based material for vascular corrosion casting with improved physical and imaging characteristics. *Microsc Res Tech* 2006 Feb;69:138–147

**Extended Data Figure 3.**
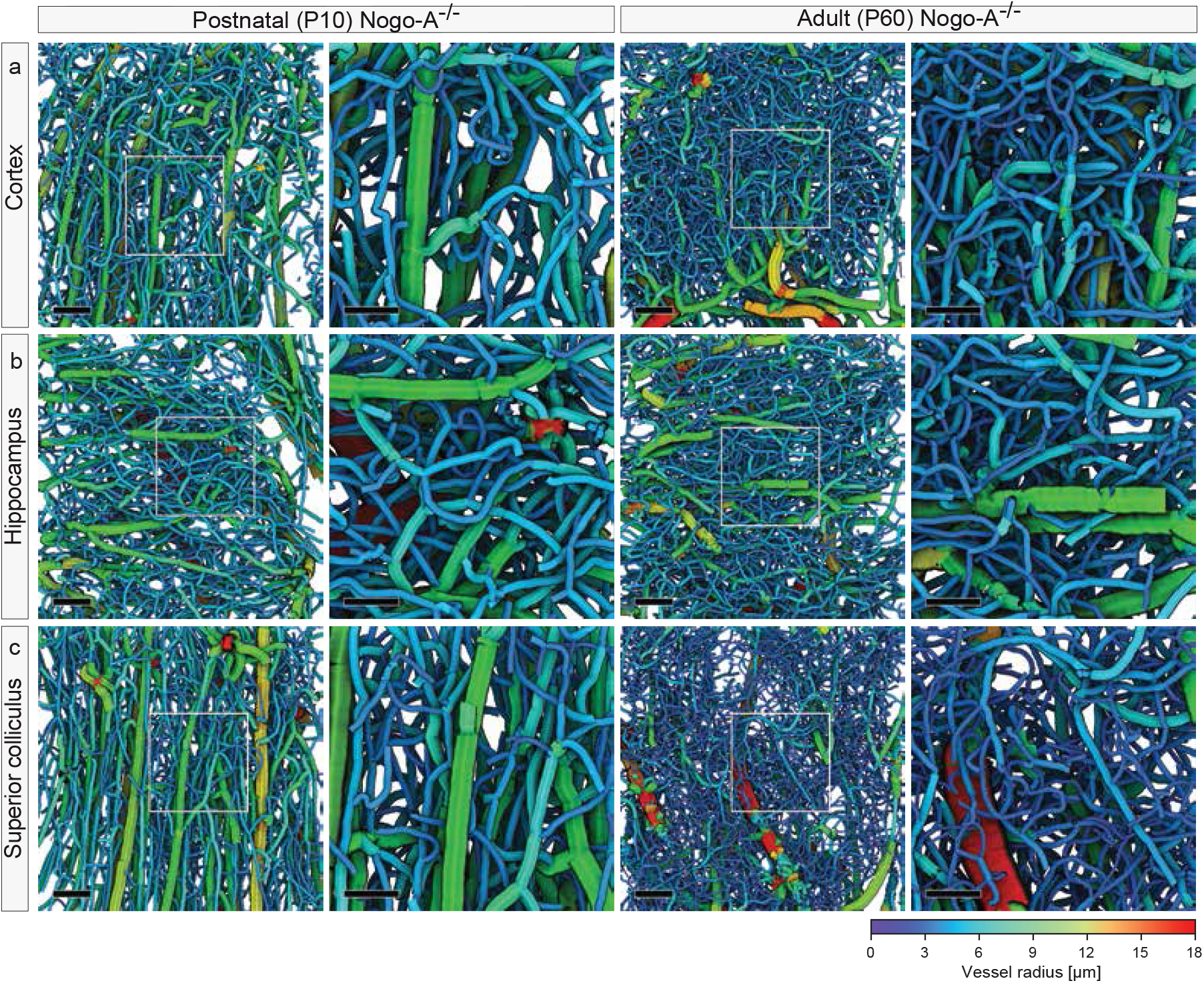
Global vascular network morphometry - Increased vascular volume fraction in various regions of the adult (P60) Nogo-A^−/−^ versus postnatal (P10) Nogo-A^−/−^ mouse brain. Genetic deletion of Nogo-A leads to increased vascular volume fraction in various regions of the postnatal mouse brain. **a-c** Computational 3D reconstructions of μCT scans of vascular networks of P10 and P60 Nogo-A^−/−^ mice displayed with color-coded vessel thickness. The increased vessel density in the Nogo-A^−/−^ animals, as compared to the WT animals (Figure 5) in the cortices (**a**), hippocampi (**b**) and superior colliculi (**c**) is obvious. Color bar indicates vessel radius. The boxed areas are enlarged at right. (n = 11 for P10 Nogo-A^−/−^ ; n = 13 for P60 Nogo-A^−/−^ animals; and in average three ROIs per animal and brain region were used). Scale bars: 100 µm (**a-c** overviews); 50 µm (**a-c** zooms).

**Extended Data Figure 4.**
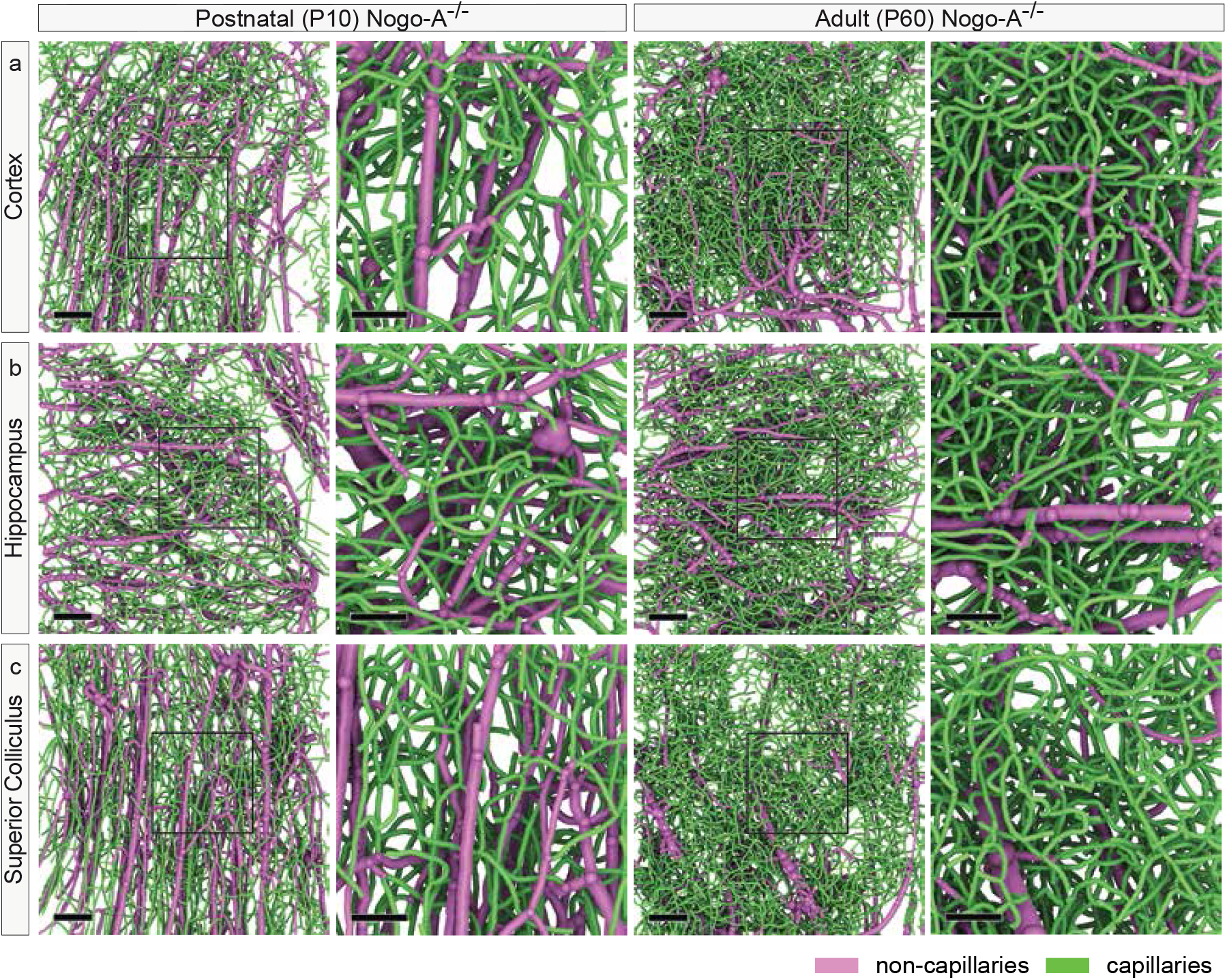
Local vascular network topology – Increased vascular volume fraction of the adult (P60) Nogo-A^−/−^ versus postnatal (P10) Nogo-A^−/−^ mouse brain mainly found at the capillary level. Genetic deletion of Nogo-A leads to increased vascular volume fraction in various regions of the P10 mouse brain. **a** Computational 3D reconstruction of μCT images of vascular networks of P10 and P60 Nogo-A^−/−^ mice separated for non-capillaries (magenta) and capillaries (green). The increased density of capillaries (green) in the P60 and P10 Nogo-A^−/−^ samples is evident when compared to the WT animals (Figure 6), non-capillaries (magenta) appear largely unchanged. The boxed areas are enlarged at right (**a-c**). Scale bars: 100 mm (**a-c**, overviews); 50 mm (**a-c**, zooms).

**Extended Data Figure 5.**
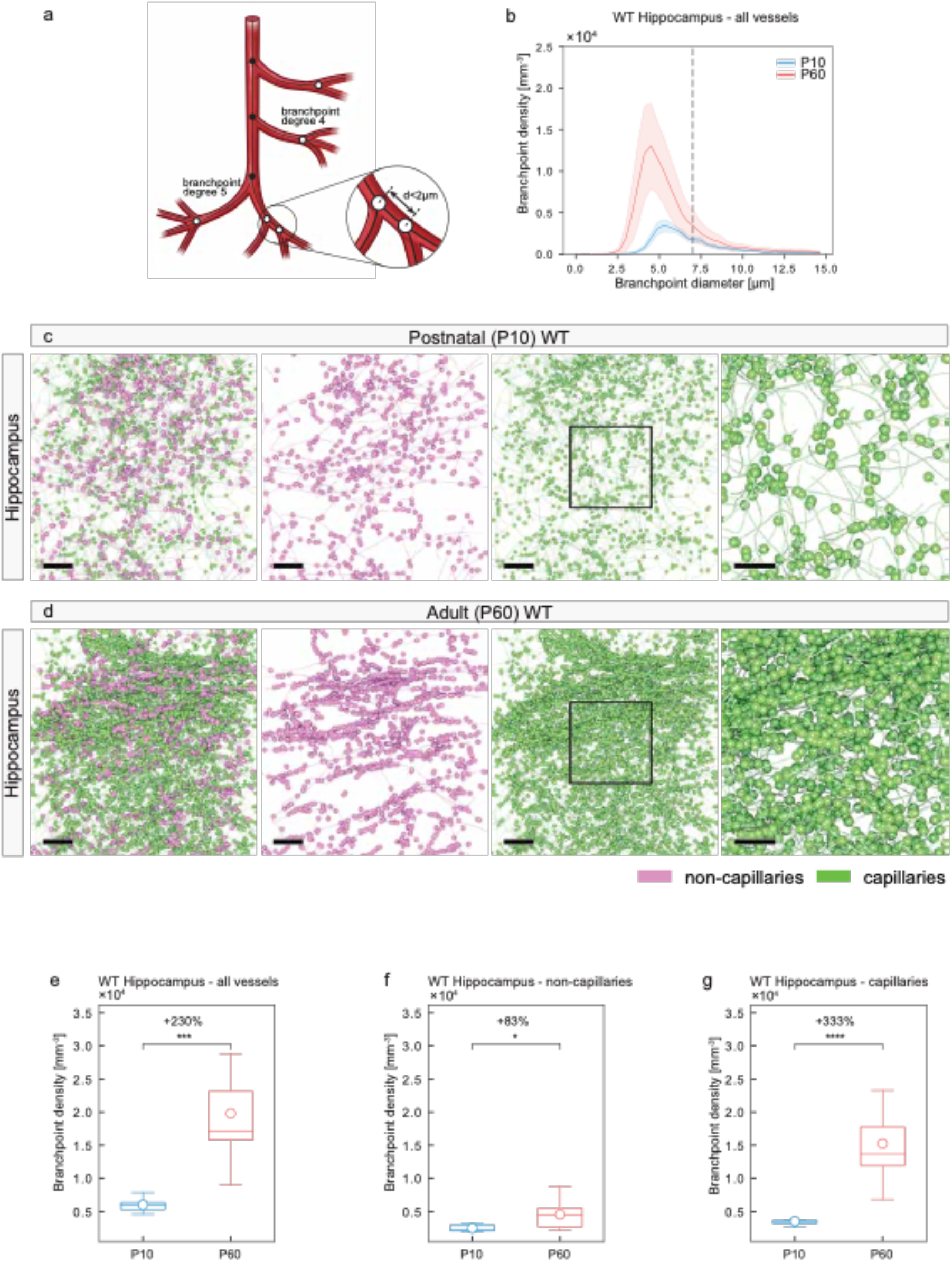
Local vascular network topology - visualization and quantification of vascular branch point diameter and vascular branch point density in the hippocampus of the adult (P60) WT versus the postnatal (P10) WT mouse brain. The vascular branch point diameter and vascular branch point density are significantly increased in the P60 as compared to the P10 mouse brain hippocampus. **a** Scheme depicting the definition of vascular branch points (see figure 7b for details) **b** Histogram showing the distribution of branch point diameter plotted against the branch point density in P10 WT and P60 WT animals. P60 WT animals show an increased branch point density as compared to P10 WT (bin width = 0.38μm; number of bins = 40). Black dashed line marks the separation between capillaries (< 7 μm) and non-capillaries (≥ 7 μm), as defined in Figure 6. **c, d** Computational 3D reconstructions of μCT images of vascular networks in the hippocampi of P10 WT and P60 WT mice with visualizations of the vessel branch points displayed as dots, separated for non-capillaries (magenta) and capillaries (green). The higher density of branch points in the P60 WT hippocampi especially at the capillary level (green) is obvious, a slight increase of branch point density can be observed at the non-capillary level (magenta). The boxed areas are enlarged at right. **e–g** Quantitative analysis of the branch point density for all vessels (**e**), non-capillaries (**f**), and capillaries (**g**) in P10 WT and P60 WT hippocampi by local morphometry analysis. The significant increase of the branch point density for all vessels in the P60 WT animals (**e**) was mainly due to a significant increase at the level of capillaries (**g**) and in part due to a significant increase at the level of non-capillaries (**f**). In average, n = 2-7 animals were used for the hippocampi and in average three ROIs per animal and brain region were used. All data are shown as mean distributions where the open dot represents the mean. Boxplots indicate the 25% to 75% quartiles of the data. The shaded blue and red areas indicate the SD. *P<0.05, **P<0.01, ***P<0.001. Scale bars: 100 µm (**c-d** overviews); 50 µm (**c-d** zooms).

**Extended Data Figure 6.**
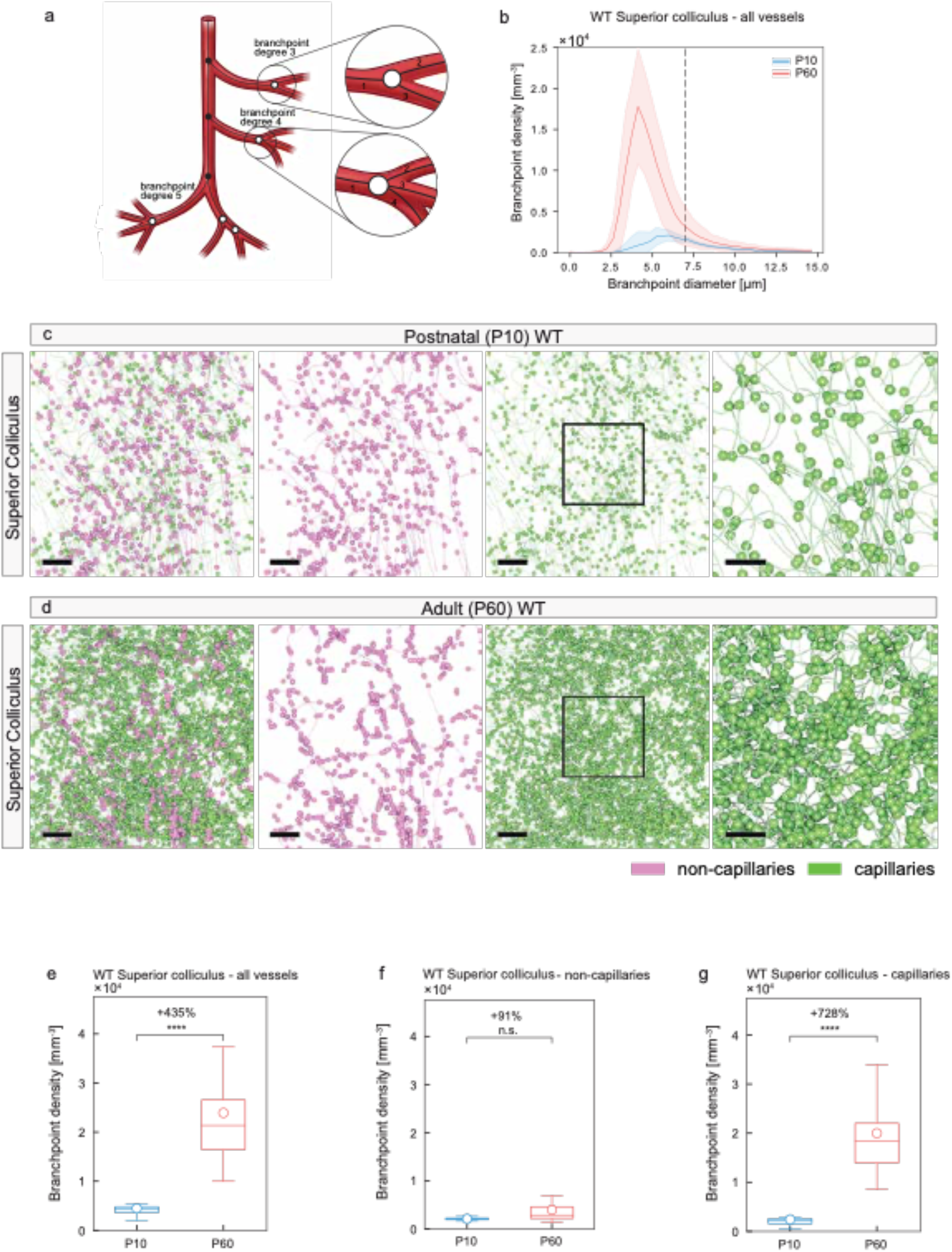
Local vascular network topology - visualization and quantification of vascular branch point diameter and vascular branch point density in the superior colliculus of the adult (P60) WT versus the postnatal (P10) WT mouse brain. The vascular branch point diameter and vascular branch point density are significantly increased in the P60 as compared to the P10 mouse brain superior colliculus. **a** Scheme depicting the definition of degree 3 and 4 vascular branch point (see figure 7b for details). **b** Histogram showing the distribution of branch point diameter plotted against the branch point density in P10 WT and P60 WT animals. P60 WT animals show an increased branch point density as compared to P10 WT mice mainly at the capillary level (bin width = 0.38; number of bins = 40). Black dashed line marks the separation between capillaries (< 7 μm) and non-capillaries (≥ 7 μm), as defined in Figure 6. **c,d** Computational 3D reconstructions of μCT images of vascular networks of the superior colliculi of P10 WT and P60 WT mice with visualizations of the vessel branch points displayed as dots, separately for non-capillaries (magenta) and capillaries (green). The higher density of branch points in the P60 WT superior colliculi especially at the capillary level (green) is obvious, a slight increase of branch point density can be observed at the non-capillary level (magenta). The boxed areas are enlarged at right. **e–g** Quantitative analysis of the branch point density for all vessels (**e**), non-capillaries (**f**), and capillaries (**g**) in P10 WT and P60 WT superior colliculi by local morphometry analysis. The significant increase of the branch point density for all vessels in the P60 WT animals (**e**) was mainly due to a significant increase at the level of capillaries (**g**) and in part due to a significant increase at the level of non-capillaries (**f**). In average, n = 2-4 animals were used for the superior colliculus; and in average three ROIs per animal and brain region were used. All data are shown as mean distributions where the open dot represents the mean. Boxplots indicate the 25% to 75% quartiles of the data. The shaded blue and red areas indicate the SD. *P<0.05, **P<0.01, ***P<0.001. Scale bars: 100 μm (**c,d**, overviews); 50 μm (**c,d**, zooms).

**Extended Data Figure 7.**
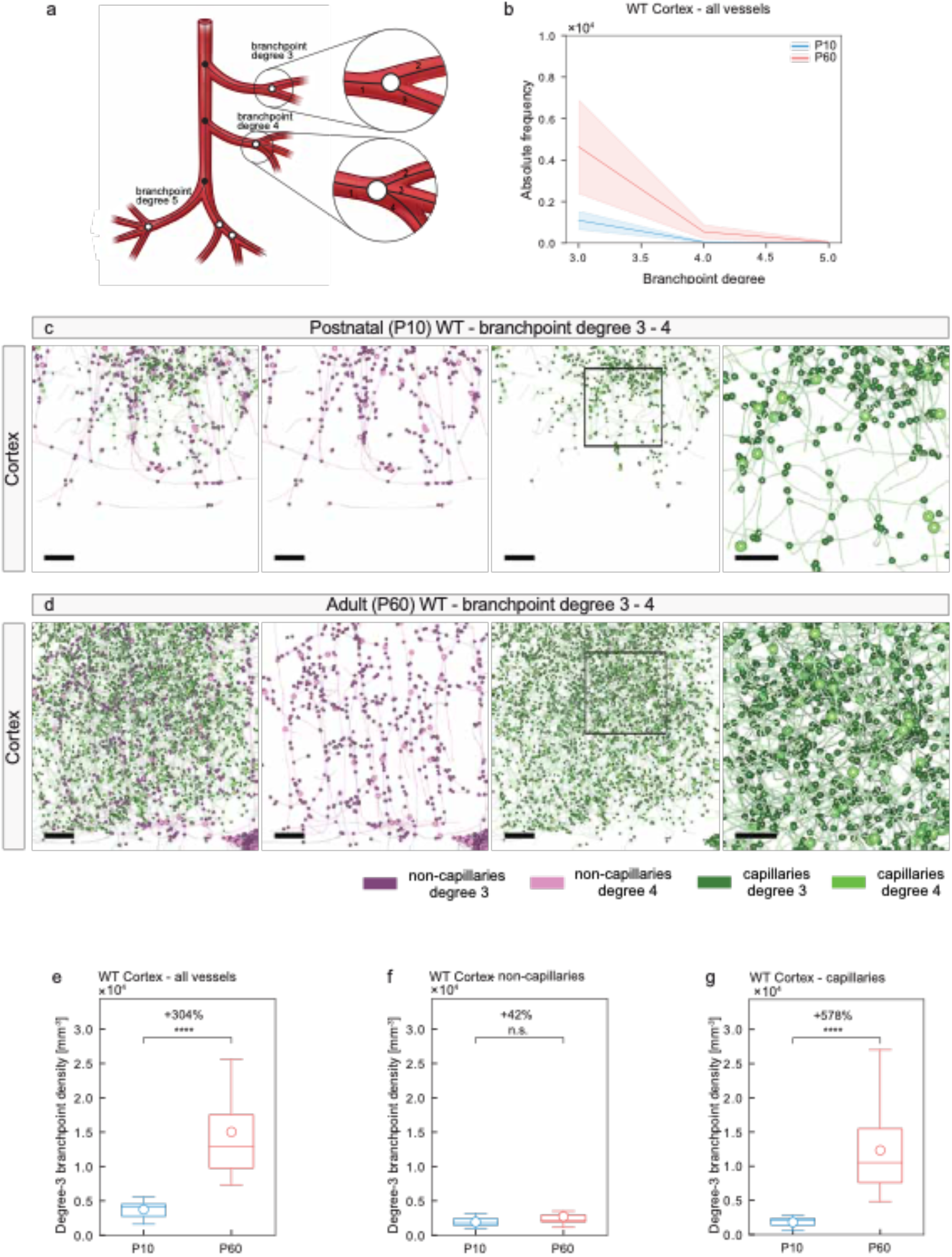
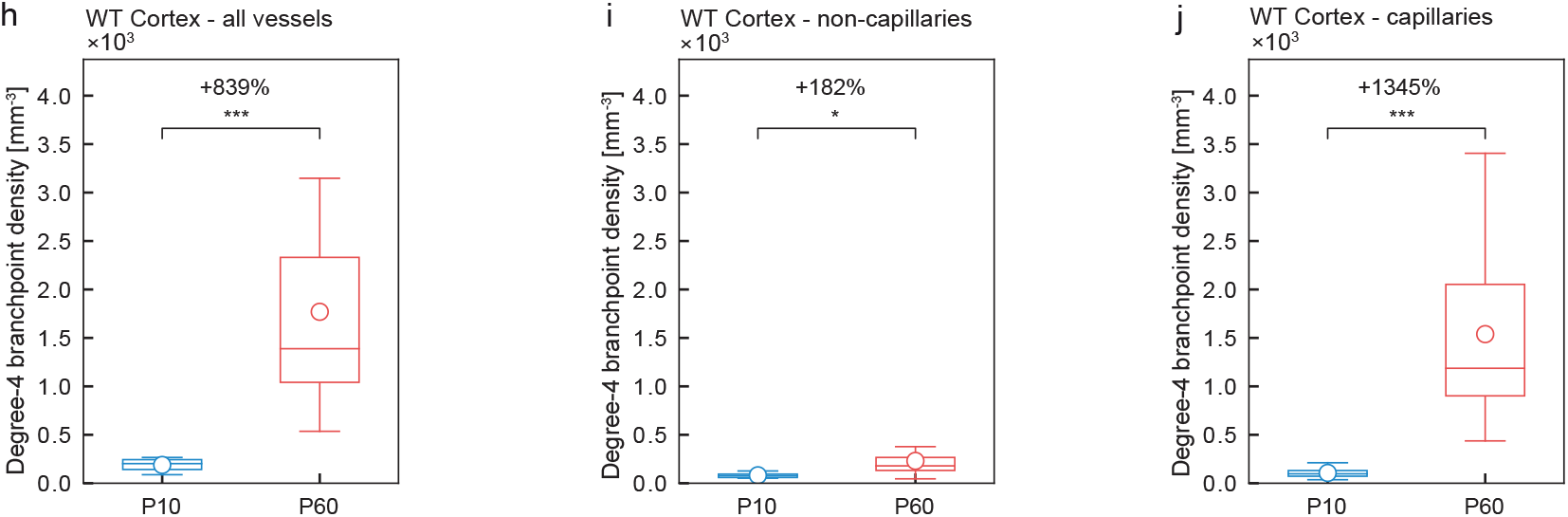
Local vascular network topology – visualization and quantification of vascular branch point diameter, vascular branch point density and branch point degree in the cortex of the adult (P60) WT versus the postnatal (P10) WT mouse brain. The vascular branch point diameter and vascular branch point density are significantly increased in the P60 as compared to the P10 mouse brain cortex. **a** Scheme depicting the definition of degree 3 and 4 vascular branch point (see figure 7b for details). **b** Histogram showing the distribution of branch point degree in P10 WT and P60 WT animals. P60 WT animals show an increased branch point degree as compared to P10 WT mice mainly at the capillary level. Of note, nearly all branch points had a degree of 3. Branch point degrees of four or even higher accounted together for far less than 1% of all branch points (bin width = 1; number of bins = 3). **c, d** Computational 3D reconstructions of µCT images of cortical vascular networks of P10 WT and P60 WT mice with visualizations of the vessel branch points displayed as dots, separately for non-capillaries (magenta) and capillaries (green) and for branch point degree 3 and 4 where branch points with degree 4 are enlarged and brighter than branch points with degree 3 (also see legend below **d**). The higher density of branch points in the P60 WT cortices especially at the capillary level (green) is obvious, a slight increase of branch point density can be observed at the non-capillary level (magenta). Furthermore, an increase in the branch points with degree 4 can be observed in the P60 WT (**d**) as compared to the P10 WT (**c**) reconstructions. The boxed areas are enlarged at right (**c,d**). Quantitative analysis of the branch point degree-3 density for all vessels (**e**), non-capillaries (**f**), and capillaries (**g**) in P10 WT and P60 WT cortices by local morphometry analysis shows that the significant increase in density of branch points with degree-3 for all vessels in P60 WT animals (**e**) was mainly due to a significant increase at the level of capillaries (**g**) and in part due to a significant increase at the level of non-capillaries (**f**). **h-j** The same quantitative analysis for branch point degree-4 (**h-j**) shows that the significant increase of the branch points with degree-4 for all vessels in P60 WT animals (**h**) was mainly due to a highly significant and very large increase at the level of capillaries (**j**) and in part due to a significant increase at the level of non-capillaries (**i**). In average, n = 3-12 animals were used for the cortices data; and in average three ROIs per animal were used). All data are shown as mean distributions where the open dot represents the mean. Boxplots indicate the 25% to 75% quartiles of the data. The shaded blue and red areas indicate the SD. *P<0.05, **P<0.01, ***P<0.001. Scale bars: 100 µm (**c, d**, overviews); 50 µm (**c, d**, zooms).

**Extended Data Figure 8.**
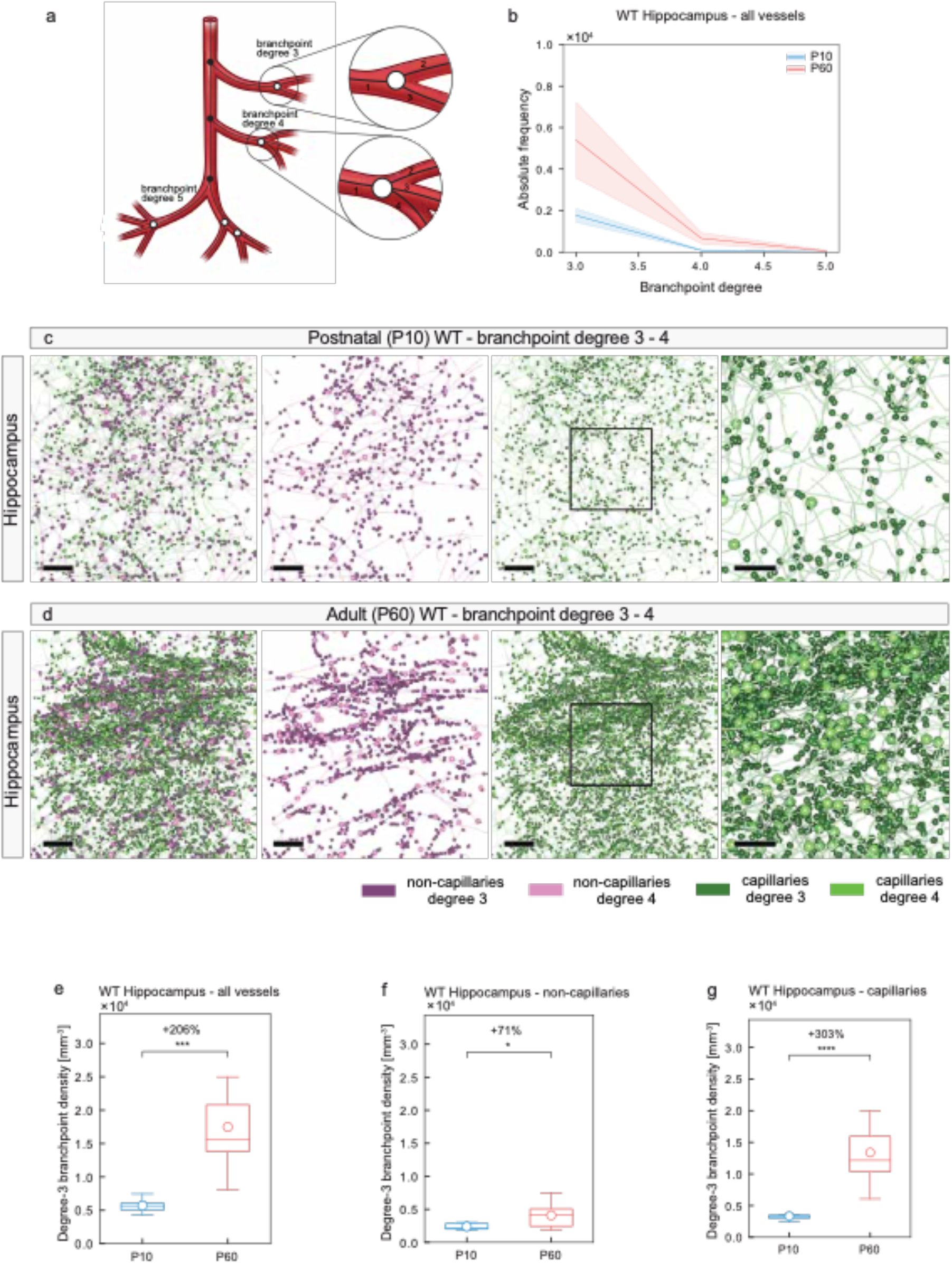
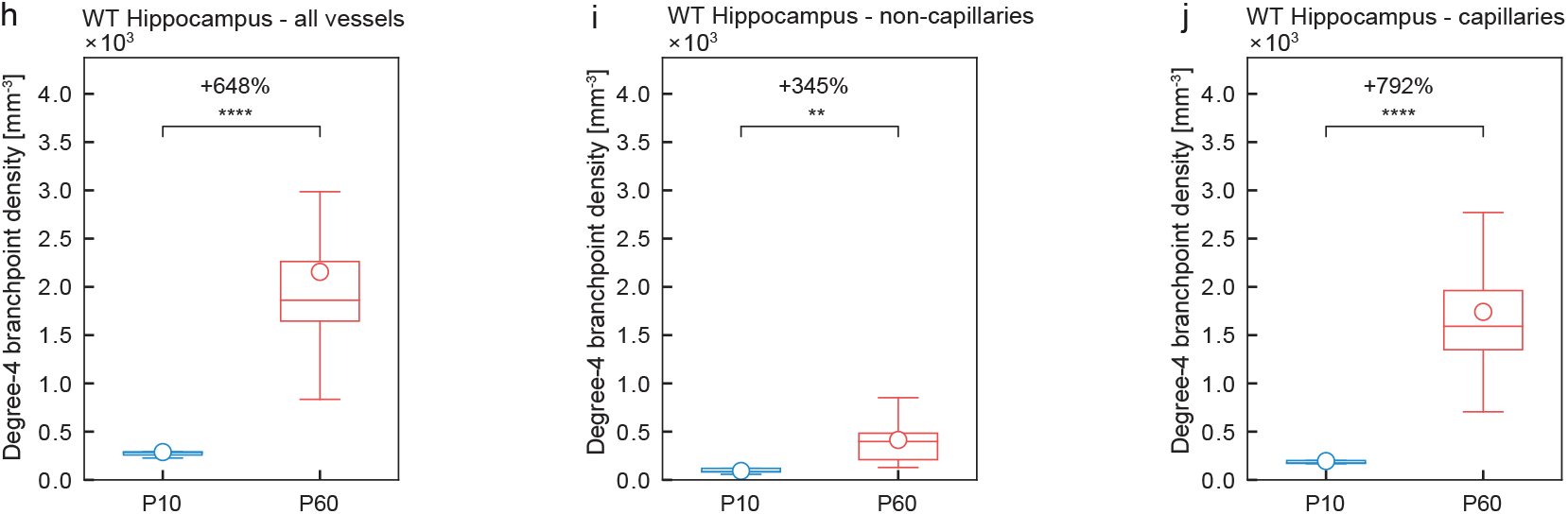
Local vascular network topology - visualization and quantification of vascular branch point diameter, vascular branch point density and branch point degree in the hippocampus in the adult (P60) WT versus the postnatal (P10) mouse brain. The vascular branch point diameter and vascular branch point density are significantly increased in the P60 as compared to P10 mouse brain hippocampus. **a** Scheme depicting the definition of vascular branch points (see figure 7b for details). **b** Histogram showing the distribution of branch point diameter in P10 WT and P60 WT animals. P60 WT animals show an increased branch point degree as compared to P10 WT mice mainly at the capillary level. Of note, nearly all branch points had a degree of 3. Branch point degrees of four or even higher accounted together for far less than 1% of all branch points (bin width = 1; number of bins = 3). **c,d** Computational 3D reconstructions of μCT images of vascular networks of P10 WT and P60 WT mice with visualizations of the vessel branch points displayed as dots, separately for non-capillaries (magenta) and capillaries (green) and for branch point degree 3 and 4 where branch points with degree 4 are enlarged and brighter than branch points with degree 3 (see legend). The higher density of branch points in the P60 WT hippocampi especially at the capillary level (green) is obvious, a slight increase of branch point density can be observed at the non-capillary level (magenta). The boxed areas are enlarged at right (**c,d**). Quantitative analysis of the branch point degree-3 density for all vessels (**e**), non-capillaries (**f**), and capillaries (**g**) in P10 WT and P60 WT hippocampi by local morphometry analysis shows that the significant increase in density of branch points with degree-3 for all vessels in the P60 WT animals (**e**) was mainly due to a significant increase at the level of capillaries (**g**) and in part due to a significant increase at the level of non-capillaries (**f**). **h-j** The same quantitative analysis for branch point degree-4 (**h-j**) shows that the significant increase of the branch points with degree-4 for all vessels in the P60 WT animals (**h**) was mainly due to a highly significant and very large increase at the level of capillaries (**j**) and in part due to a significant increase at the level of non-capillaries (**i**). In average, n = 2-7 animals were used for the hippocampi; and in average three ROIs per animal and brain region were used. All data are shown as mean distributions where the open dot represents the mean. Boxplots indicate the 25% to 75% quartiles of the data. The shaded blue and red areas indicate the SD. *P<0.05, **P<0.01, ***P<0.001. Scale bars: 100 µm (**c,d**, overviews); 50 µm (**c,d**, zooms).

**Extended Data Figure 9.**
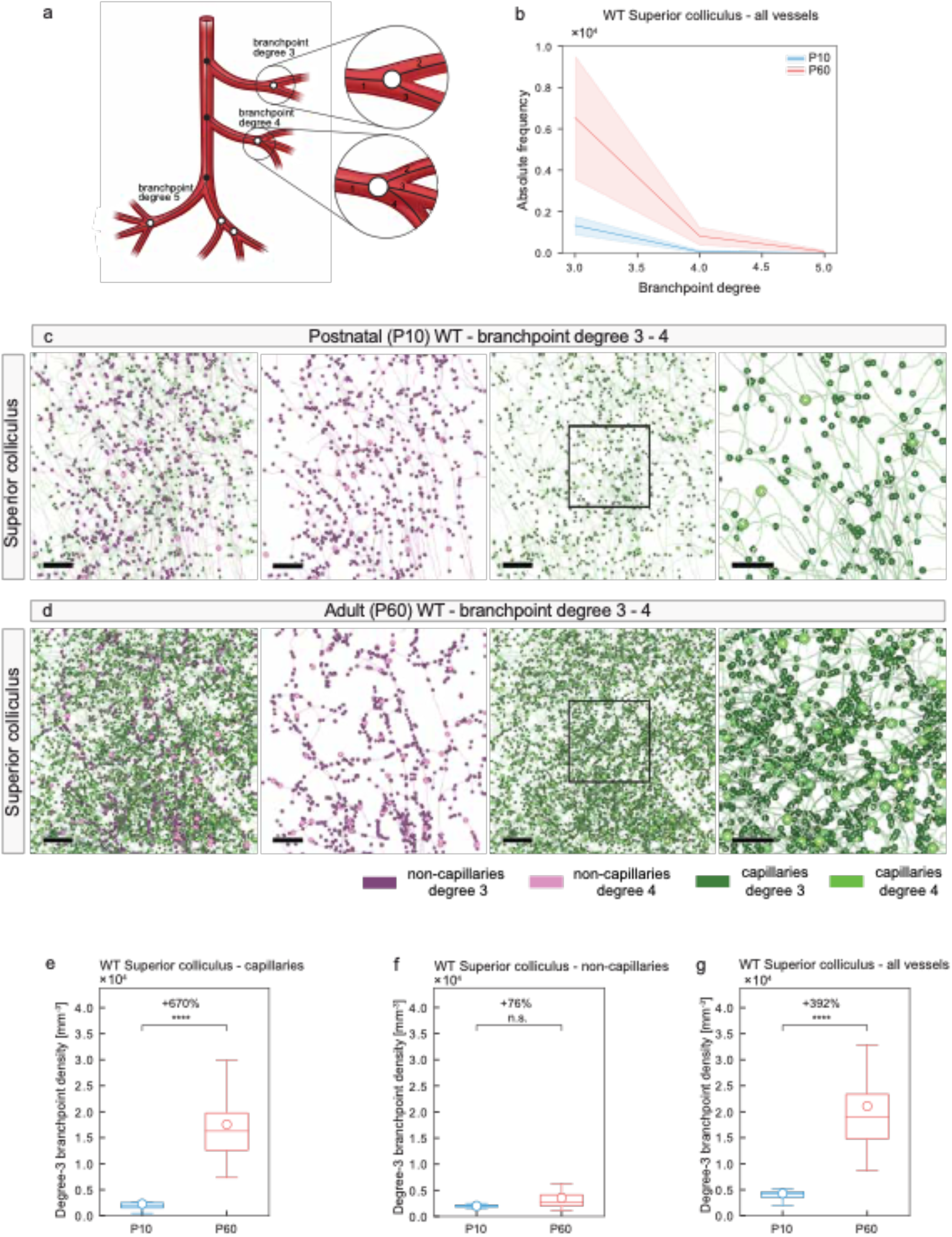
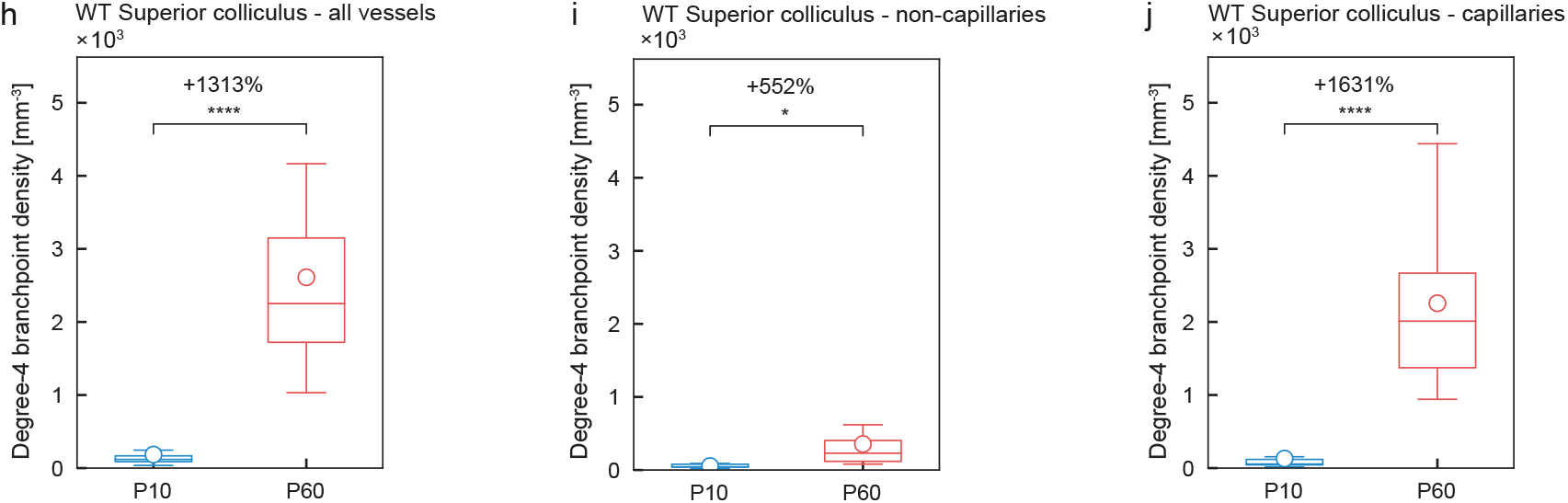
Local vascular network topology – visualization and quantification of vascular branch point diameter, vascular branch point density and branch point degree in the superior colliculus of the adult (P60) WT versus the postnatal (P10) WT mouse brain. The vascular branch point diameter and vascular branch point density are significantly increased in the P60 as compared to the P10 mouse brain superior colliculi **a** Scheme depicting the definition of vascular branch points (see figure 7b for details). **b** Histogram showing the distribution of branch point diameter in P10 WT and P60 WT animals. P60 WT animals show an increased branch point density as compared to P10 WT mice mainly at the capillary level. Of note, nearly all branch points had a degree of 3. Branch point degrees of four or even higher accounted together for far less than 1% of all branch points (bin width = 1; number of bins = 3). **c,d** Computational 3D reconstructions of μCT images of vascular networks of P10 WT and P60 WT mice with visualizations of the vessel branch points displayed as dots, separately for non-capillaries (magenta) and capillaries (green) and for branch point degree 3 and 4 where branch points with degree 4 are enlarged and brighter than branch points with degree 3 (see legend). The higher density of branch points in the P60 WT superior colliculi especially at the capillary level (green) is obvious, a slight increase of branch point density can be observed at the non-capillary level (magenta). The boxed areas are enlarged at right. Quantitative analysis of the branch point density for all vessels (**e**), non-capillaries (**f**), and capillaries (**g**) in P10 WT and P60 WT superior colliculi by local morphometry analysis. The significant increase of the branch point density for all vessels in the P60 WT animals (**e**) was mainly due to a significant increase at the level of capillaries (**g**) and in part due to a significant increase at the level of non-capillaries (**f**). **h-j** The same quantitative analysis for branch point degree-4 (**h-j**) shows that the significant increase of the branch points with degree-4 for all vessels in the P60 WT animals (**h**) was mainly due to a highly significant and very large increase at the level of capillaries (**j**) and in part due to a significant increase at the level of non-capillaries (**i**). In average, n = 2-4 animals were used for the superior colliculi; and in average three ROIs per animal and brain region were used. All data are shown as mean distributions where the open dot represents the mean. Boxplots indicate the 25% to 75% quartiles of the data. The shaded blue and red areas indicate the SD. *P<0.05, **P<0.01, ***P<0.001. Scale bars: 100 μm (**c,d,** overviews); 50 μm (**c,d,** zooms).

**Extended Data Figure 10.**
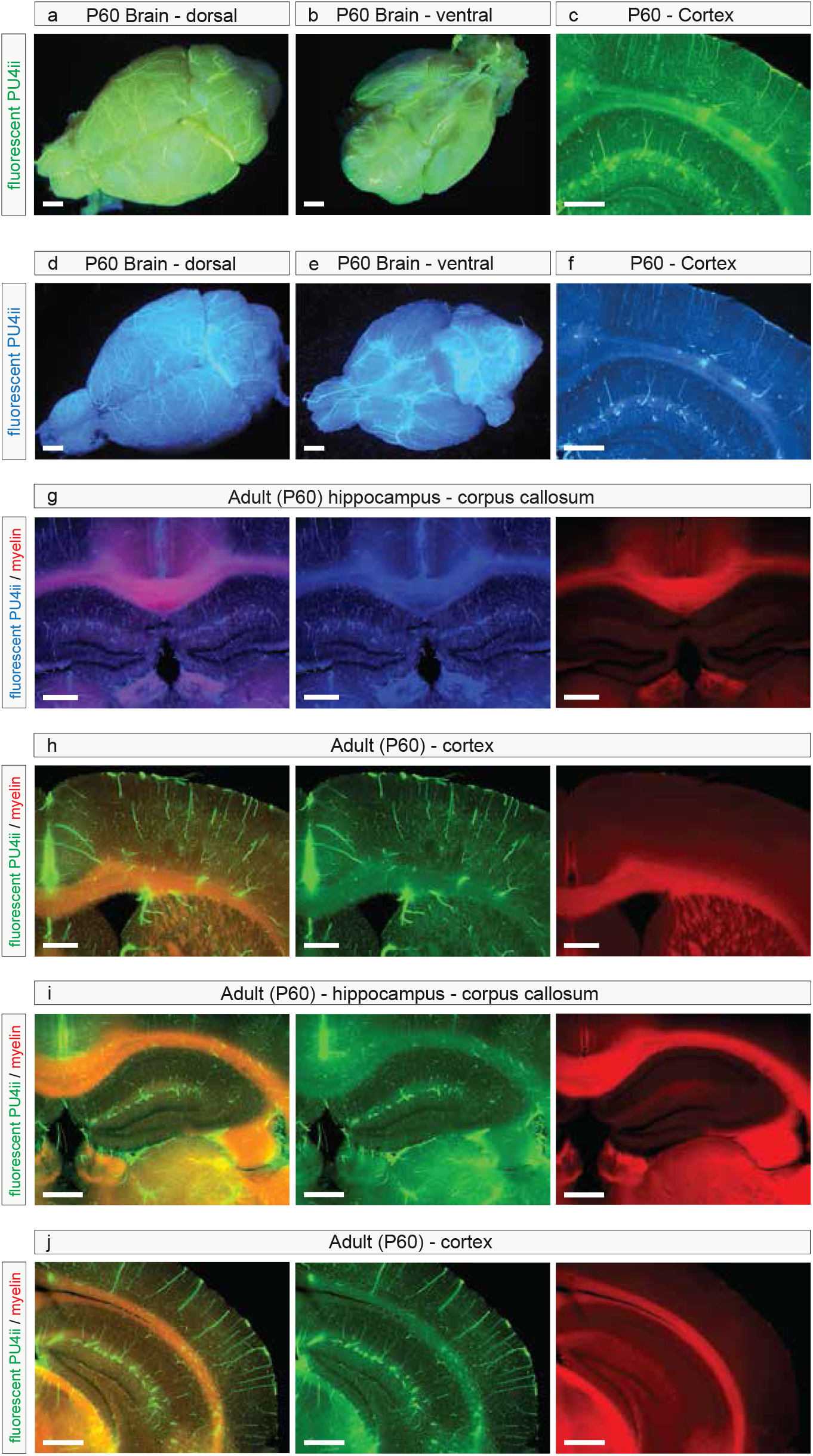
Resin PU4ii vascular corrosion casting combined with immunofluorescence. Visualization by light microscopy of the brain vasculature after the addition of fluorescent pigments to the PU4ii resin solution in the P60 mouse brain. **a,b,d,e** Overview of mouse brain with yellow/green (**a,b)** and UV-blue (**d,e**) fluorescent dyes (vasQtec) added to the resin showing the brain vasculature; overviews: dorsal (**a,d**), ventral view (**b,e**). The selection of the brain regions analyzed here has been done on 150μm coronal sections in the structures selected using the Allen Mouse Brain Atlas. **c,f,g-j** Light microscopy overviews of multiple brain regions in 150 μm histological sections of a P60 coronal mouse brain perfused with either yellow/green (**c,h-j**) or UV-blue (**f,g**) fluorescent pigments added to the PU4ii resin solution, showing the cortical layers and part of the hippocampus (**c,f**), and sections in which soft tissue was co-labeled with a red fluorescent myelin stain (FluoroMyelin Red, Molecular Probes Inc) to visualize the corpus callosum and other highly myelinated structures (red). Scale bars, 1 mm (**a,b,d,e**); 500 μm (**c,d,g-j**). Excitation/emission of the yellow/green pigments at 450nm/550nm, excitation/emission of the UV-blue fluorescent pigment at 375nm/430nm.

## SUPPLEMENTARY FIGURES

**Supplementary Figure 1.**
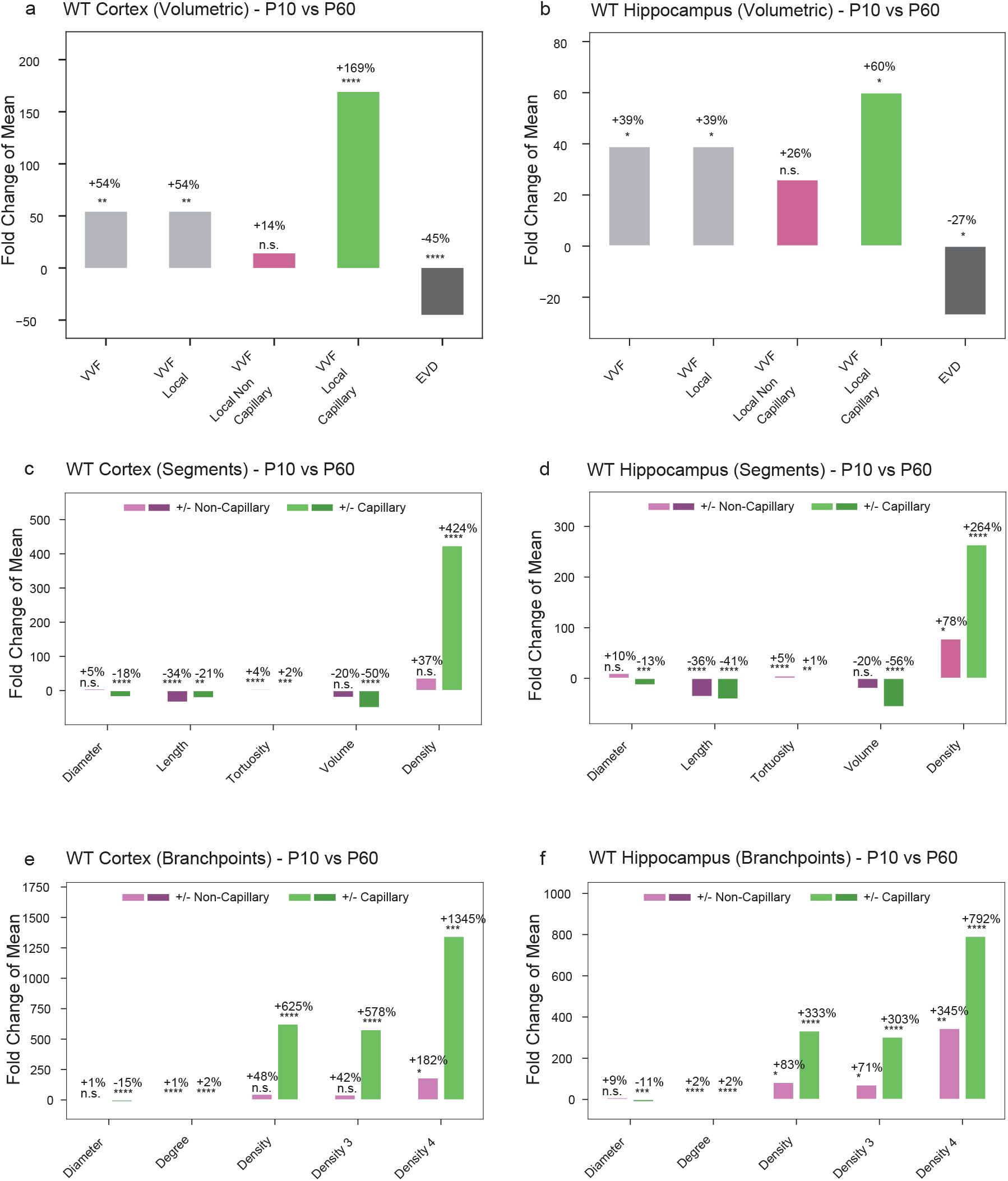
Overview of the main vascular parameters in various regions of the wildtype postnatal and adult mouse brain. Quantification of the main vascular parameters, depicted as fold change of the mean in the WT P60 mouse brain as compared to the P10 mouse brain. **a,b,g** On a volumetric level, the vascular volume fraction (VVF), was analyzed by computational analysis using the global morphometry, and local topology approach in the cortex (**a**), hippocampus (**b**) and superior colliculus (**g**). The vascular volume fraction in all these brain structures was significantly higher in the WT P60 animals than that in the P10 WT animals. **c,d,h** On the level of vascular segments, the parameters segment-diameter, - length, - tortuosity, - volume and -density are depicted as fold change of the mean. The parameters segment diameter, - length and -volume show a highly significant decrease or non-significant increase, whereas the parameters segment tortuosity and -density show an increase. All these changes are most pronounced on the capillary level. **e,f,i** Investigating the difference in branchpoint characteristics between P60 and P10 wildtype mouse brains, the vascular parameters branch point diameter, -degree, -density, branch point degree 3 density and branch point degree 4 density were analyzed. Branch point diameter shows a significant decrease in the P60 as compared to the P10 mouse brain on a capillary level in all three brain regions. The vascular parameters branch point degree, - density, branch point degree 3 density and branch point degree 4 density all show an relative increase, again most pronounced on the capillary level, in all three brain regions (n = 4-6 for P10 WT; n = 8-12 for P60 WT; n = 3-5 for P10 Nogo-A^−/−^; n = 8-11 for P60 Nogo-A^−/−^ animals; and in average three ROIs per animal and brain region were used). All data are shown as fold change of the mean distributions, where the magenta bars correspond to the non-capillaries and the green bar corresponds to the capillaries. *P < 0.05, **P < 0.01, ***P < 0.001.

**Supplementary Figure 2.**
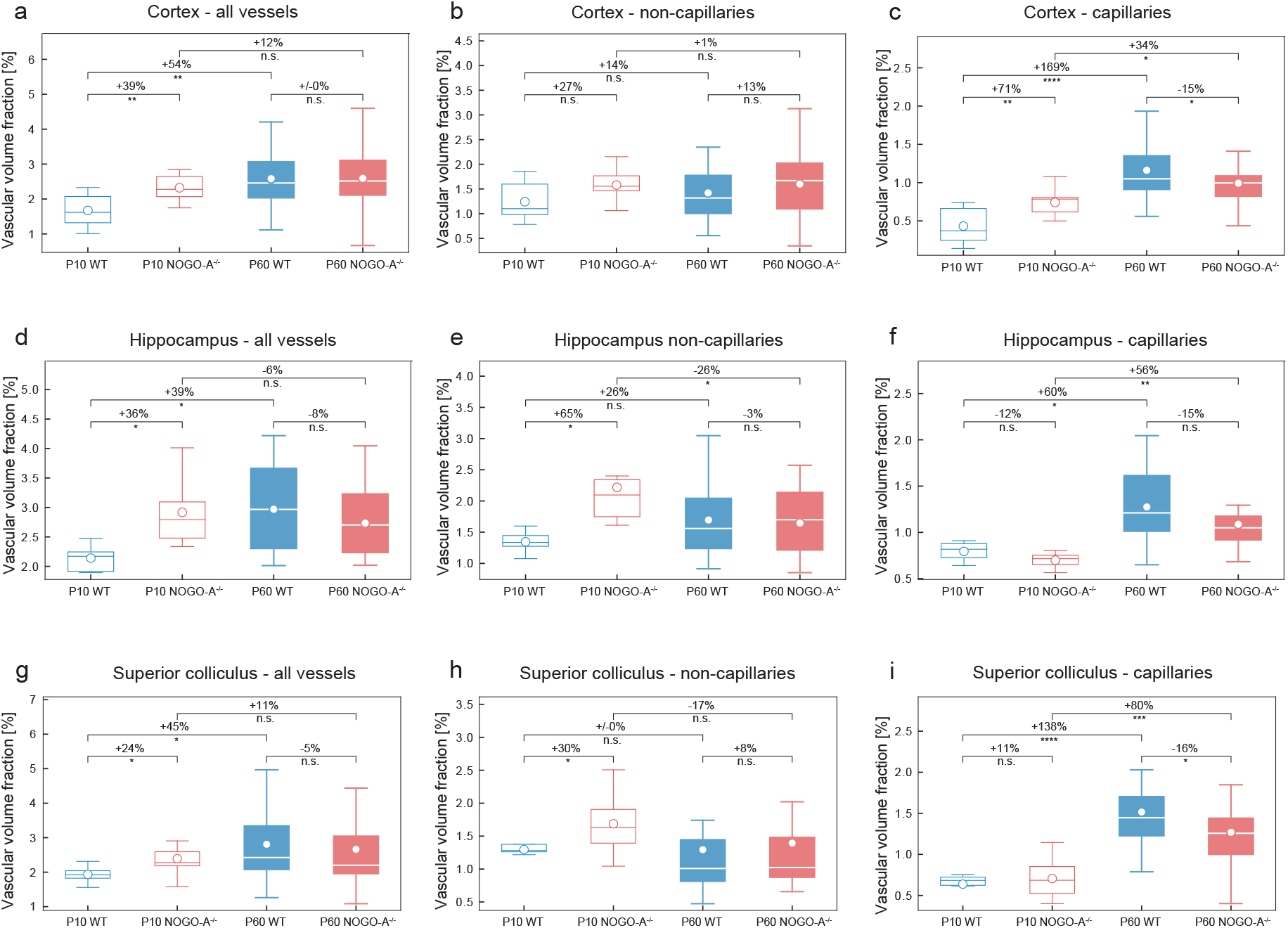
Overview of the effects of physiological ageing and of genetic deletion of Nogo-A on the vascular volume fraction in various regions of the adult (P60) and postnatal (P10) mouse brain. **a-i** Quantification of the 3D vascular volume fraction in cortex (**a-c**), hippocampus (**d-f**), and superior colliculus (**g-i**), by computational analysis using global morphometry. The vascular volume fraction in all these brain structures was significantly higher in the Nogo-A deficient animals than that in the WT control animals. This same increase can be observed in the P60 WT group, when compared to the P10 WT animals, and is most pronounced at the level of capillaries (**c,f,i**) (n = 4-6 for P10 WT; n = 8-12 for P60 WT; n = 3-5 for P10 Nogo-A^−/−^; n = 8-11 for P60 Nogo-A^−/−^ animals; and in average three ROIs per animal and brain region were used). *P < 0.05, **P < 0.01, ***P < 0.001.

**Supplementary Figure 3.**
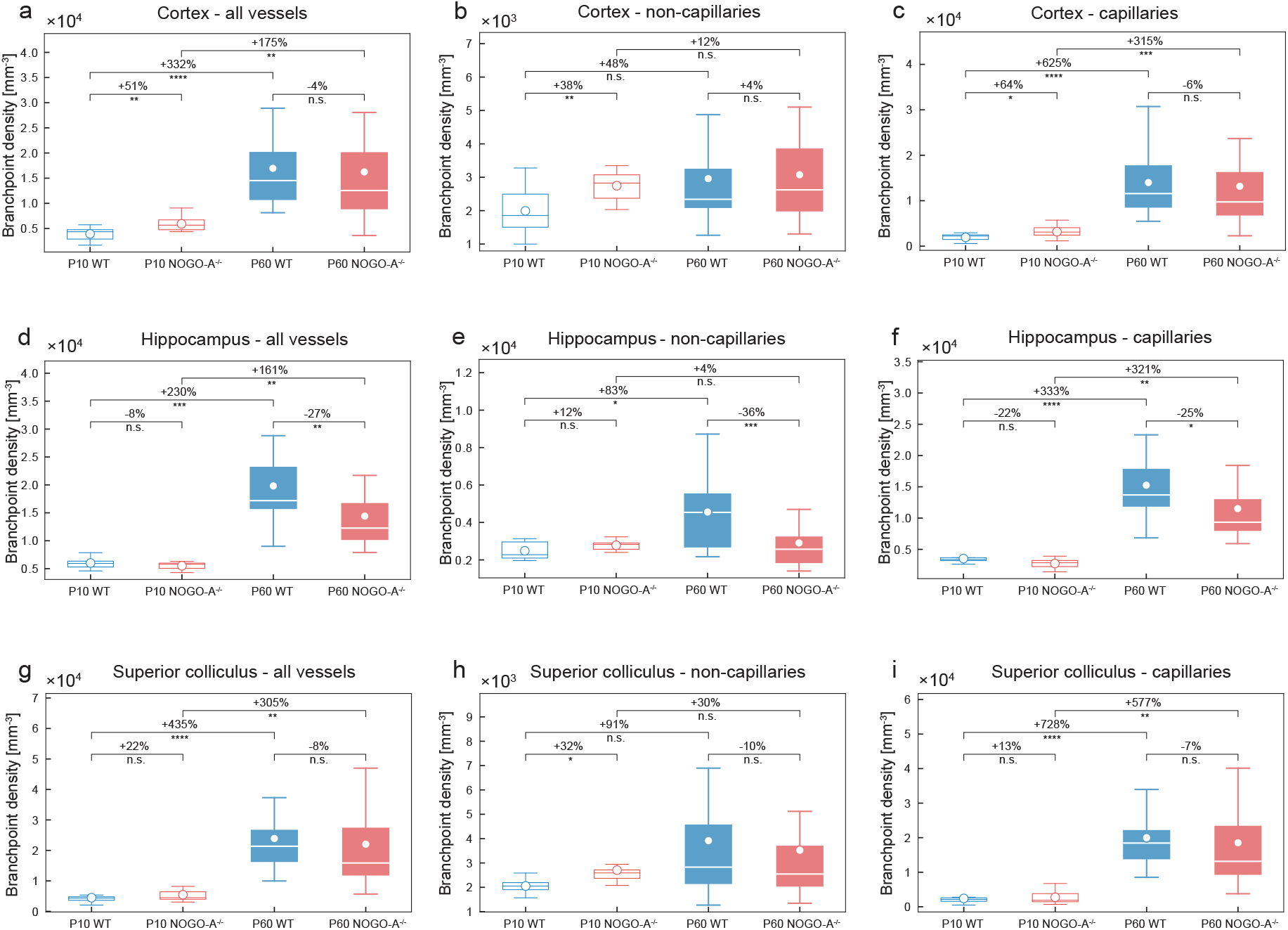
Overview of the effects of physiological ageing and of genetic deletion of Nogo-A on the branch point density in various regions of the adult (P60) and postnatal (P10) mouse brain. **a-i** Quantification of the branch point density in cortex (**a-c**), hippocampus (**d-f**), and superior colliculus (**g-i**), by local topology analysis. The branch point density in all these brain structures was significantly higher in the Nogo-A deficient animals than that in the WT control animals. This same increase can be observed in the P60 WT group, when compared to the P10 WT animals, and is most pronounced at the level of capillaries (**c,f,i**) (n = 4-6 for P10 WT; n = 8-12 for P60 WT; n = 3-5 for P10 Nogo-A^−/−^; n = 8-11 for P60 Nogo-A^−/−^ animals; and in average three ROIs per animal and brain region were used). *P < 0.05, **P < 0.01, ***P < 0.001.

**Supplementary Figure 4.**
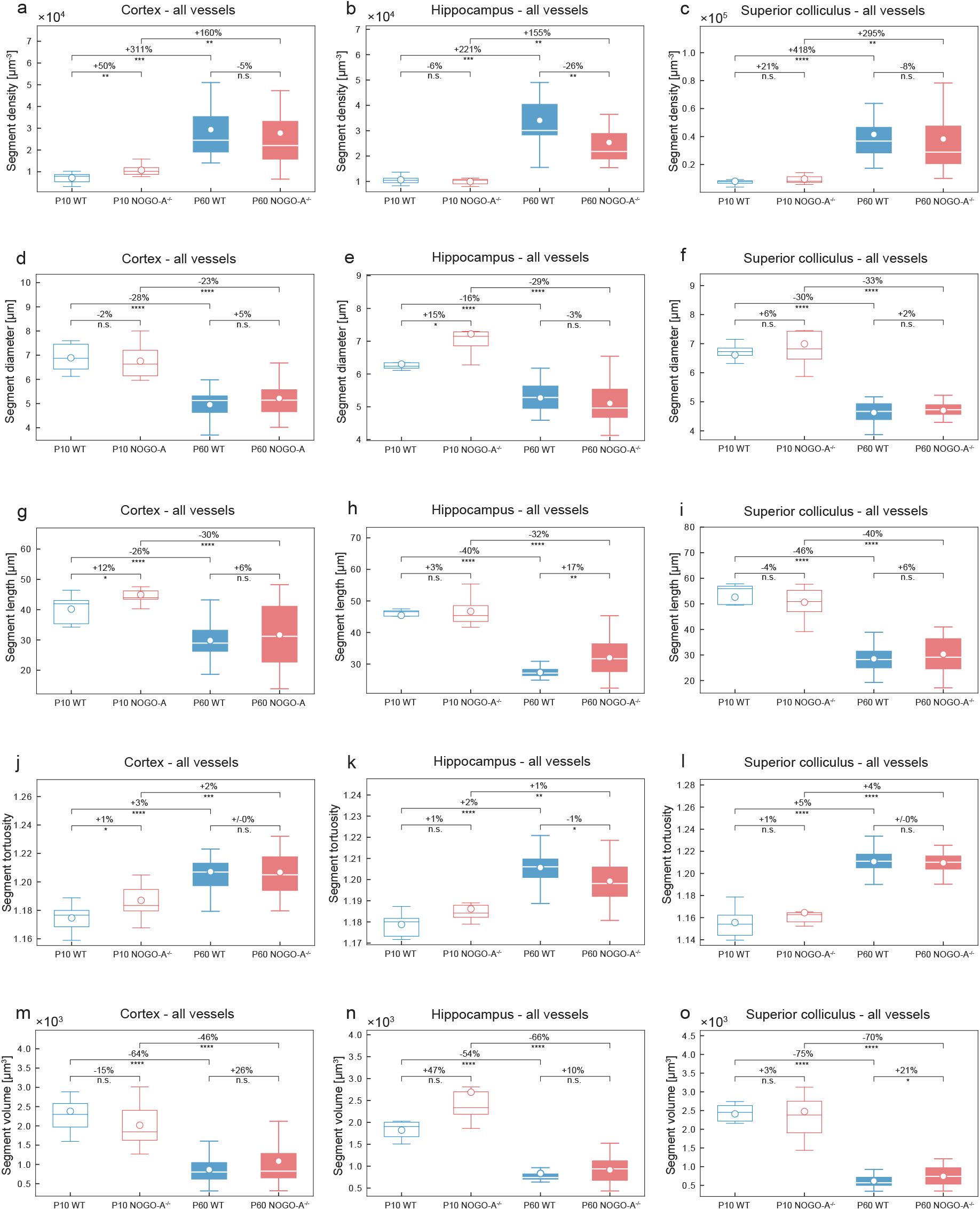
Overview of the effects of physiological ageing and of genetic deletion of Nogo-A on the segment density, -diameter, -length, -tortuosity and -volume in various regions of the adult (P60) and postnatal (P10) mouse brain. **a-o** Quantification of segment density, -diameter, -length, -tortuosity and -volume in the cortex (**a,d,g,j,m**), hippocampus (**b,e,h,k,n**), and superior colliculus (**c,f,i,l,o**), by local topology analysis. The segment density and -tortuosity in all these brain structures was significantly higher in the Nogo-A deficient animals than that in the WT control animals. This same increase can be observed in the P60 WT group, when compared to the P10 WT animals, and is most pronounced at the level of capillaries (**c,l**). The parameters segment diameter, -length, and - volume are significantly decreased in the Nogo-A deficient animals as compared to the WT control animals. Here, also this same increase, predominantly at the capillary level (**f,i,l**) can be observed in the P60 WT group, when compared to the P10 WT animals (n = 4-6 for P10 WT; n = 8-12 for P60 WT; n = 3-5 for P10 Nogo-A^−/−^; n = 8-11 for P60 Nogo-A^−/−^ animals; and in average three ROIs per animal and brain region were used). *P < 0.05, **P < 0.01, ***P < 0.001.

**Supplementary Figure 5.**
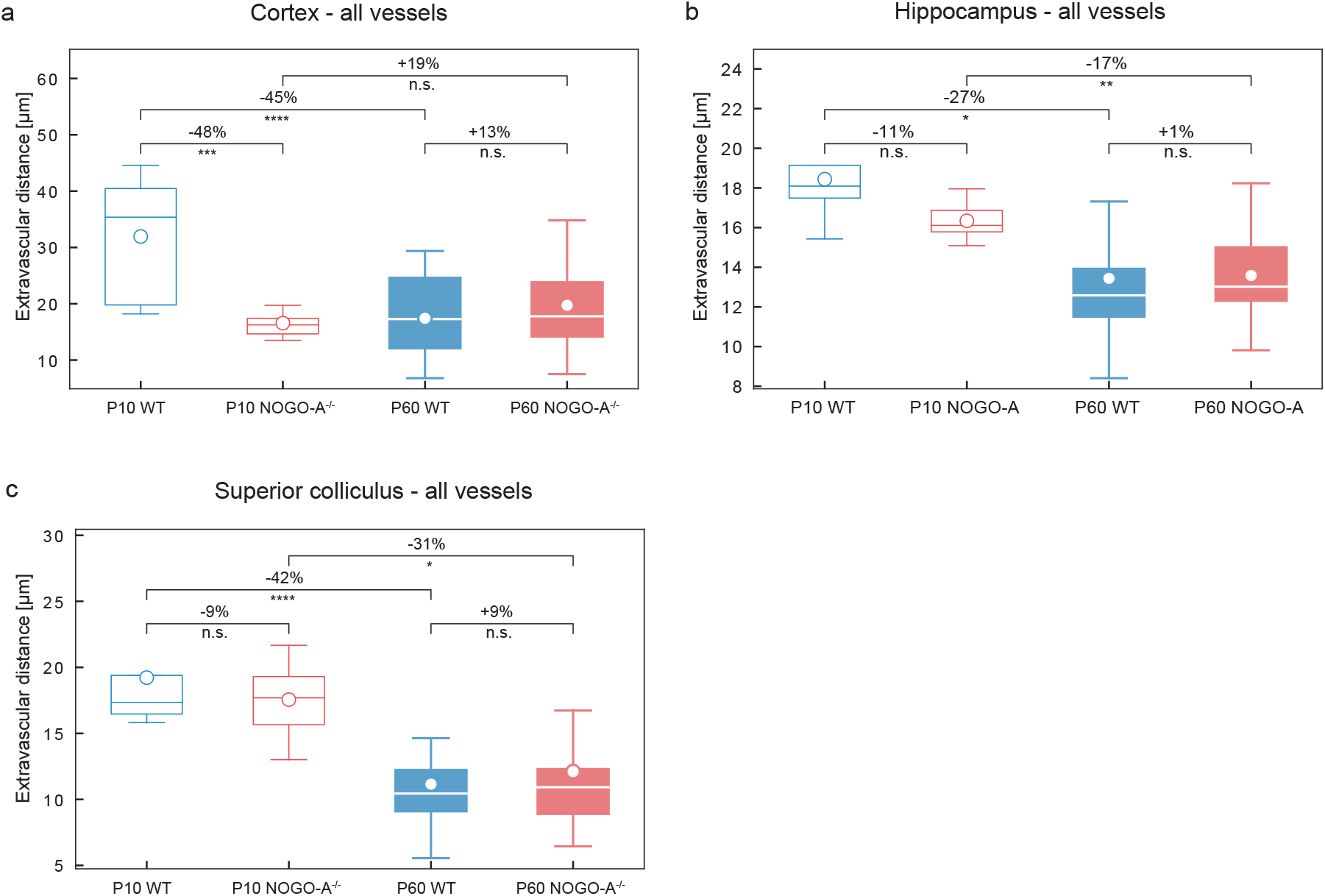
Overview of the effects of physiological ageing and of genetic deletion of Nogo-A on the extravascular distance in various regions of the adult (P60) and postnatal (P10) mouse brain. **a-c** Quantification of the extravascular distance in P10 and P60 in the three brain regions calculated by global morphometry analysis. The extravascular distance in the cortices (**a**), hippocampi (**b**) and superior colliculi (**c**) was lower in the P10 Nogo-A deficient animals than that in the WT control animals, whereas this effect was less obvious in the P60 Nogo-A versus WT (n = 4-6 for P10 WT; n = 8-12 for P60 WT; n = 3-5 for P10 Nogo-A^−/−^; n = 8-11 for P60 Nogo-A^−/−^ animals; and in average three ROIs per animal and brain region were used). All data are shown as mean distributions where the open dot represents the mean. Boxplots indicate the 25% to 75% quartiles of the data. The shaded blue and red areas indicate the SD. *P<0.05, **P<0.01, ***P<0.001.

**Supplementary Figure 6.**
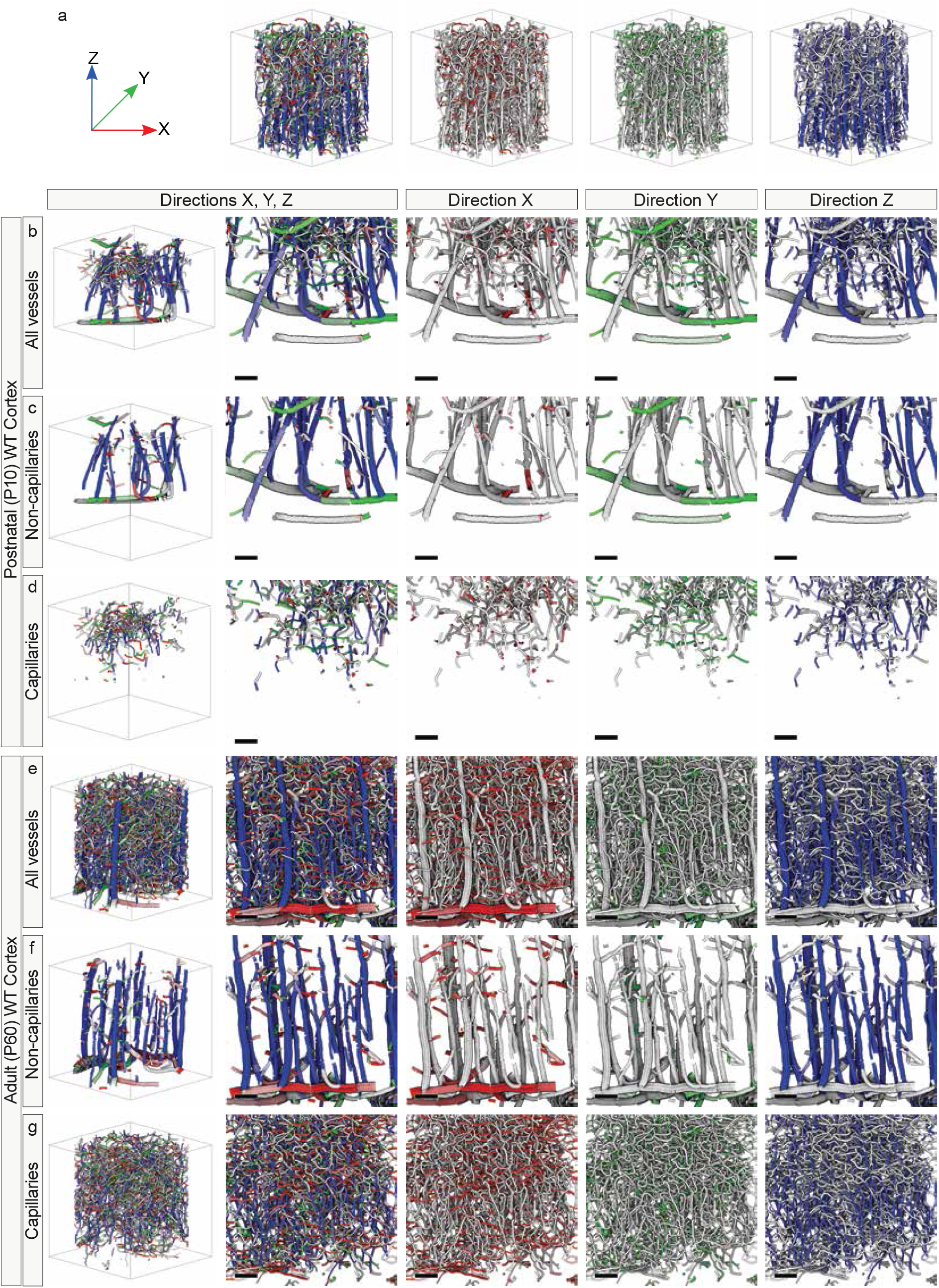
Global vascular network morphology – visualization of distinct patterns of vessel directionality in the cortex of the adult (P60) WT and postnatal (P10) WT mouse brain with a division into capillaries and non-capillaries. **a** Computational 3D reconstruction of µCT scans of vascular networks of P60 WT displayed with color-coded directionality (see figure 10 for details). **b-g** Computational 3D reconstruction of µCT scans color-coded for vessel directionality depicting the P10 WT (**b-d**) and P60 WT (**e-g**) cortex. A division was made between all vessels (**b,e**), non-capillaries (**c,f**), and capillaries (**d,g**), defined by an inner diameter of ≥ 7 µm for non-capillaries for < 7µm for capillaries. Cortical renderings clearly show that the superficial perineural vascular plexus (PNVP) extended in the x- and y-directions, whereas the interneural vascular plexus (INVP) exhibited a radial sprouting pattern into the brain parenchyma along the z-axis, perpendicular to the PNVP. This pattern is most pronounced at the level of non-capillaries (**c,f**). Scale bars: 100 µm (**b-g**).

**Supplementary Figure 7.**
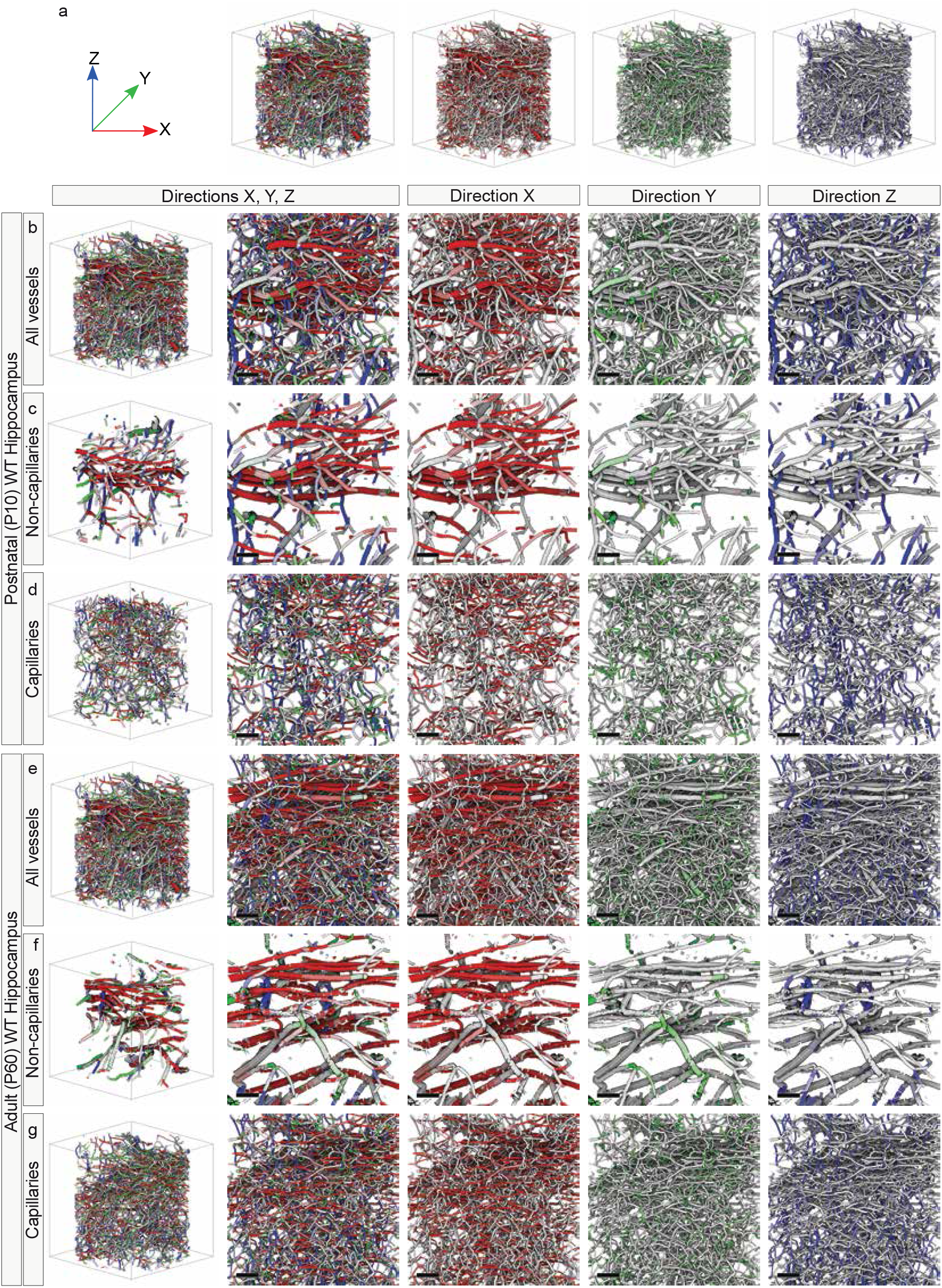
Global vascular network morphology – visualization of distinct patterns of vessel directionality in the hippocampus of the adult (P60) WT and postnatal (P10) WT mouse brain with a division into capillaries and non-capillaries. **a** Computational 3D reconstruction of µCT scans of vascular networks of P60 WT displayed with color-coded directionality (see figure 10 for details). **b-g** Computational 3D reconstruction of µCT scans color-coded for vessel directionality depicting the P10 WT (**b-d**) and P60 WT (**e-g**) hippocampus. A division was made between all vessels (**b,e**), non-cap illaries (**c,f**), and capillaries (**d,g**), defined by an inner diameter of ≥ 7 µm for non-capillaries for < 7µm for capillaries. In the hippocampus, horizontally orientated main vessel branches in the x-axis were recognizable, most likely following the distinct curved shape of this anatomical structure. This pattern is most pronounced at the level of non-capillaries (**c,f**). Scale bars: 100 µm (**b-g**).

**Supplementary Figure 8.**
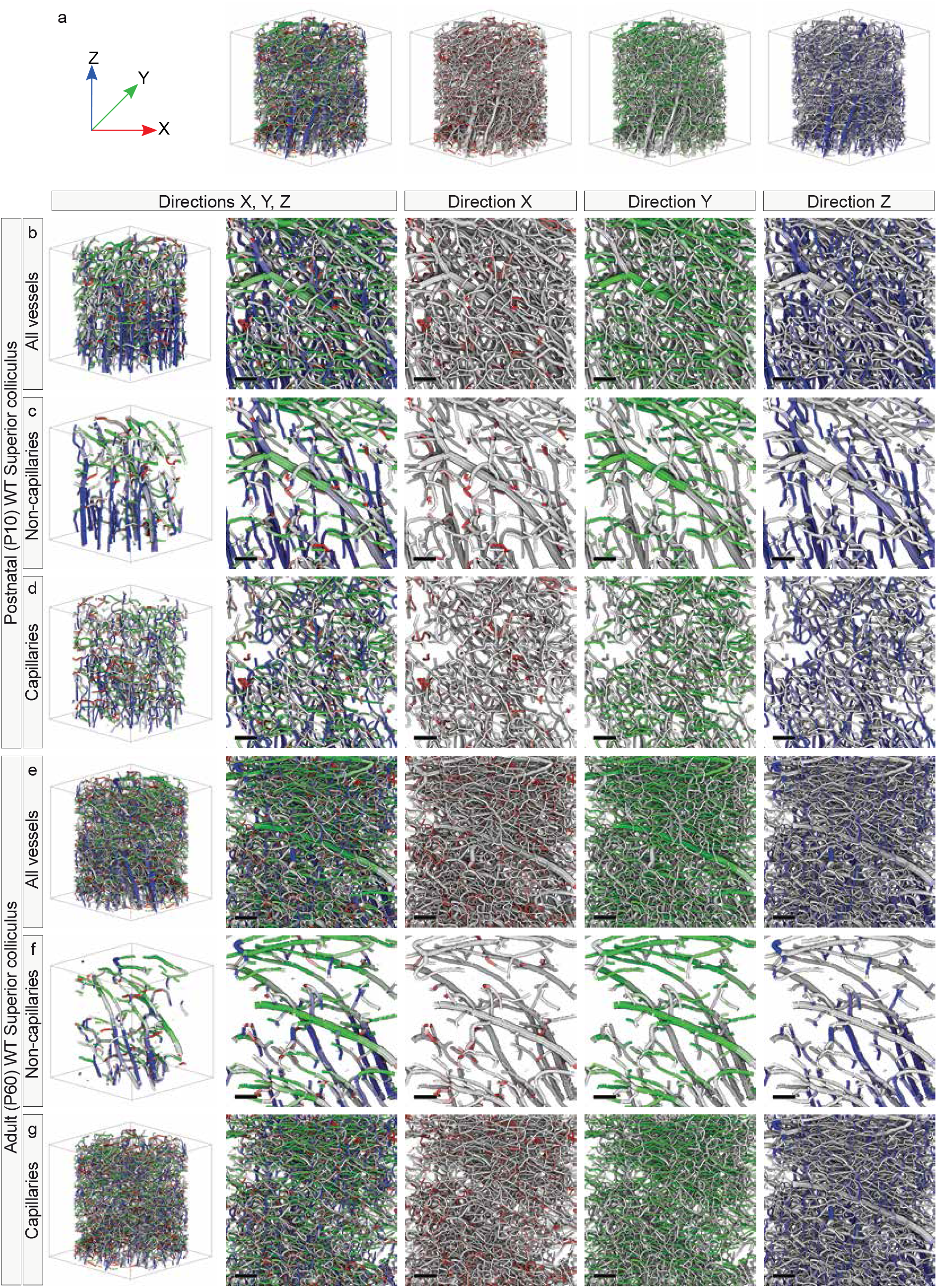
Global vascular network morphology – visualization of distinct patterns of vessel directionality in the superior colliculus of the adult (P60) WT and postnatal (P10) WT mouse brain with a division into capillaries and non-capillaries. **a** Computational 3D reconstruction of µCT scans of vascular networks of P60 WT displayed with color-coded directionality (see figure 10 for details). **b-g** Computational 3D reconstruction of µCT scans color-coded for vessel directionality depicting the P10 WT (**b-d**) and P60 WT (**e-g**) cortex. A division was made between all vessels (**b,e**), non-capillaries (**c,f**), and capillaries (**d,g**), defined by an inner diameter of ≥ 7 µm for non-capillaries for < 7µm for capillaries. In the superior colliculus, a comparable pattern of directionality as in the cortex was observed with recognizable INVP vessels radially sprouting into the brain parenchyma along the z-axis. This pattern is most pronounced at the level of non-capillaries (**c,f**). Scale bars: 100 µm (**b-g**).

**Supplementary Figure 9.**
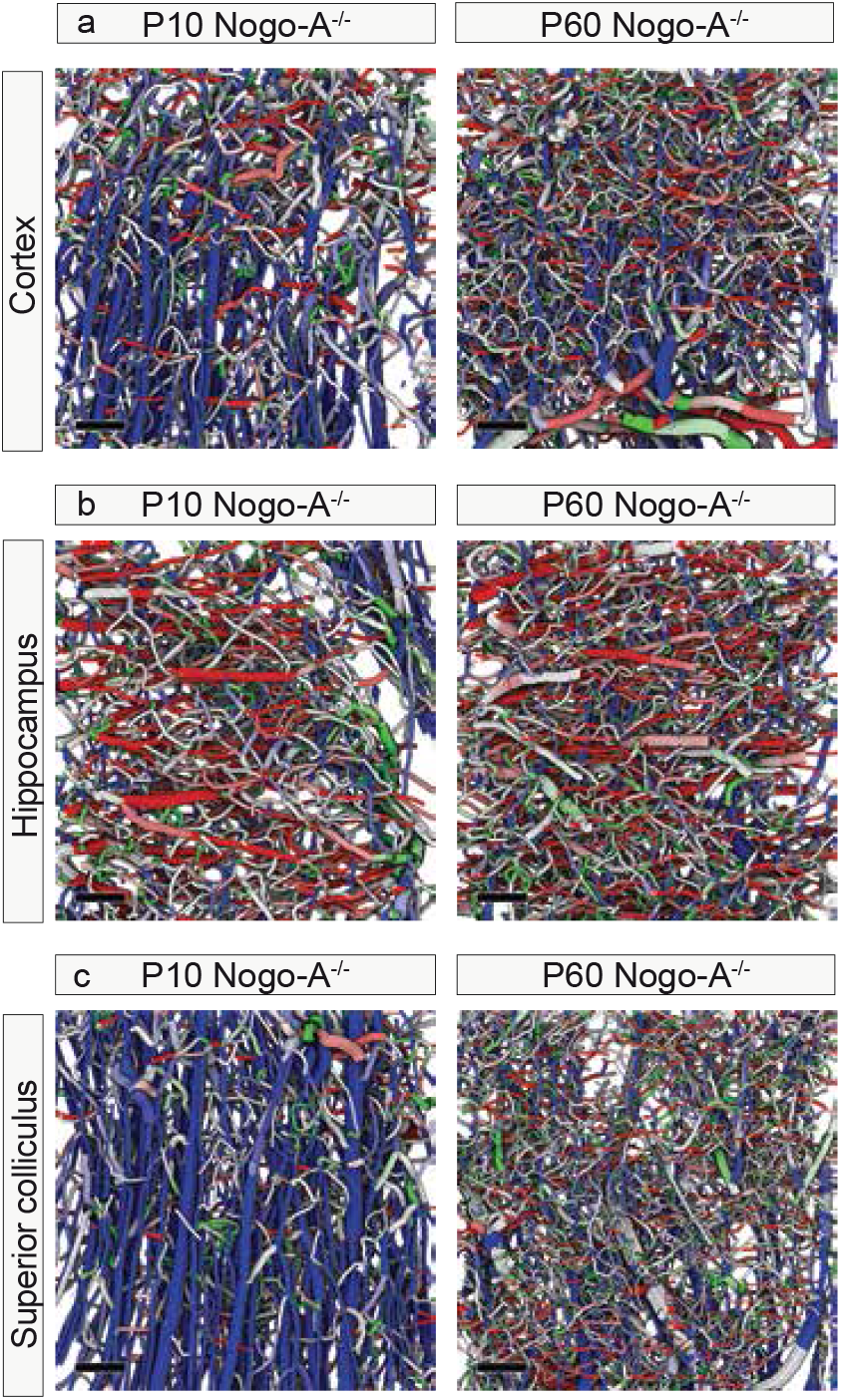
Global vascular network morphology – visualization of distinct patterns of vessel directionality in various brain regions of the adult (P60) Nogo-A^−/−^ and postnatal (P10) Nogo-A^−/−^ mouse brain. **a** Computational 3D reconstruction of µCT scans color-coded for vessel directionality depicting the P10 Nogo-A^−/−^ and P60 Nogo-A^−/−^ cortex (see figure 10 for details). Cortical renderings clearly show that the superficial perineural vascular plexus (PNVP) extended in the x- and y-directions, whereas the interneural vascular plexus (INVP) exhibited a radial sprouting pattern into the brain parenchyma along the z-axis, perpendicular to the PNVP. **b** In the hippocampus, horizontally orientated main vessel branches in the x-axis were recognizable, most likely following the distinct curved shape of this anatomical structure. **c** In the superior colliculus, a comparable pattern of directionality as in the cortex was observed with recognizable INVP vessels radially sprouting into the brain parenchyma along the z-axis. Scale bars: 100 µm (**b-g**).

**Supplementary Figure 10.**
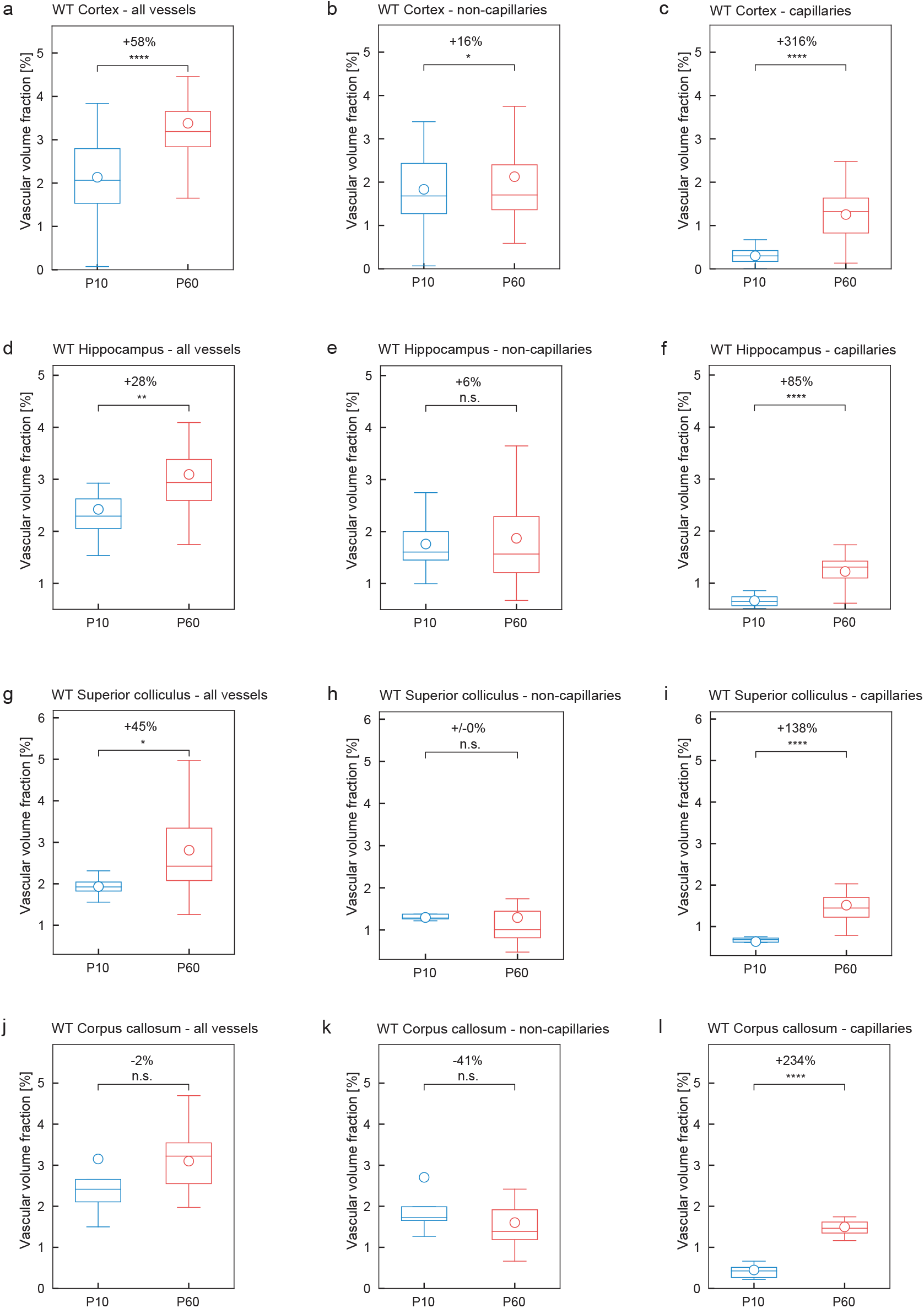
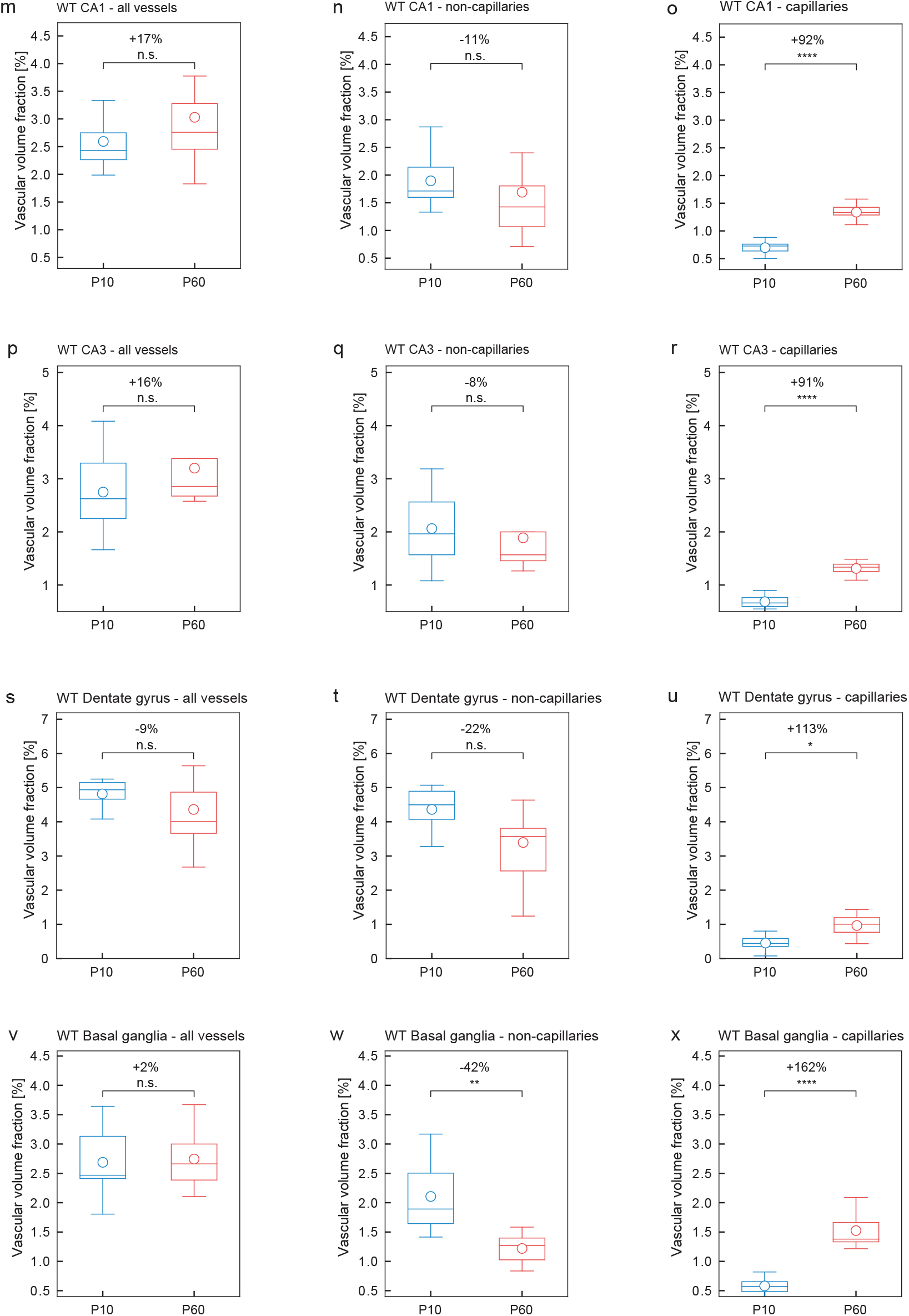
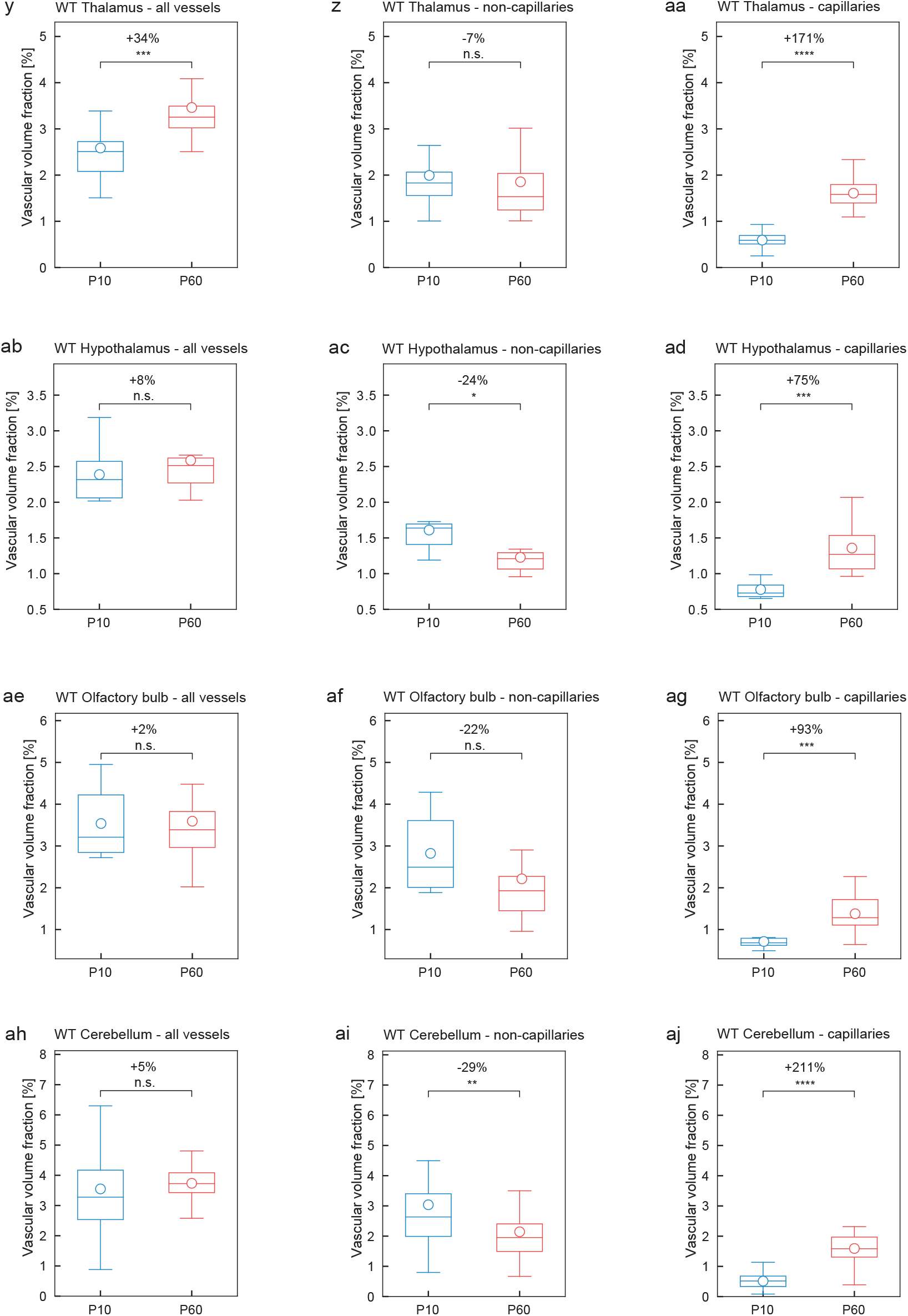
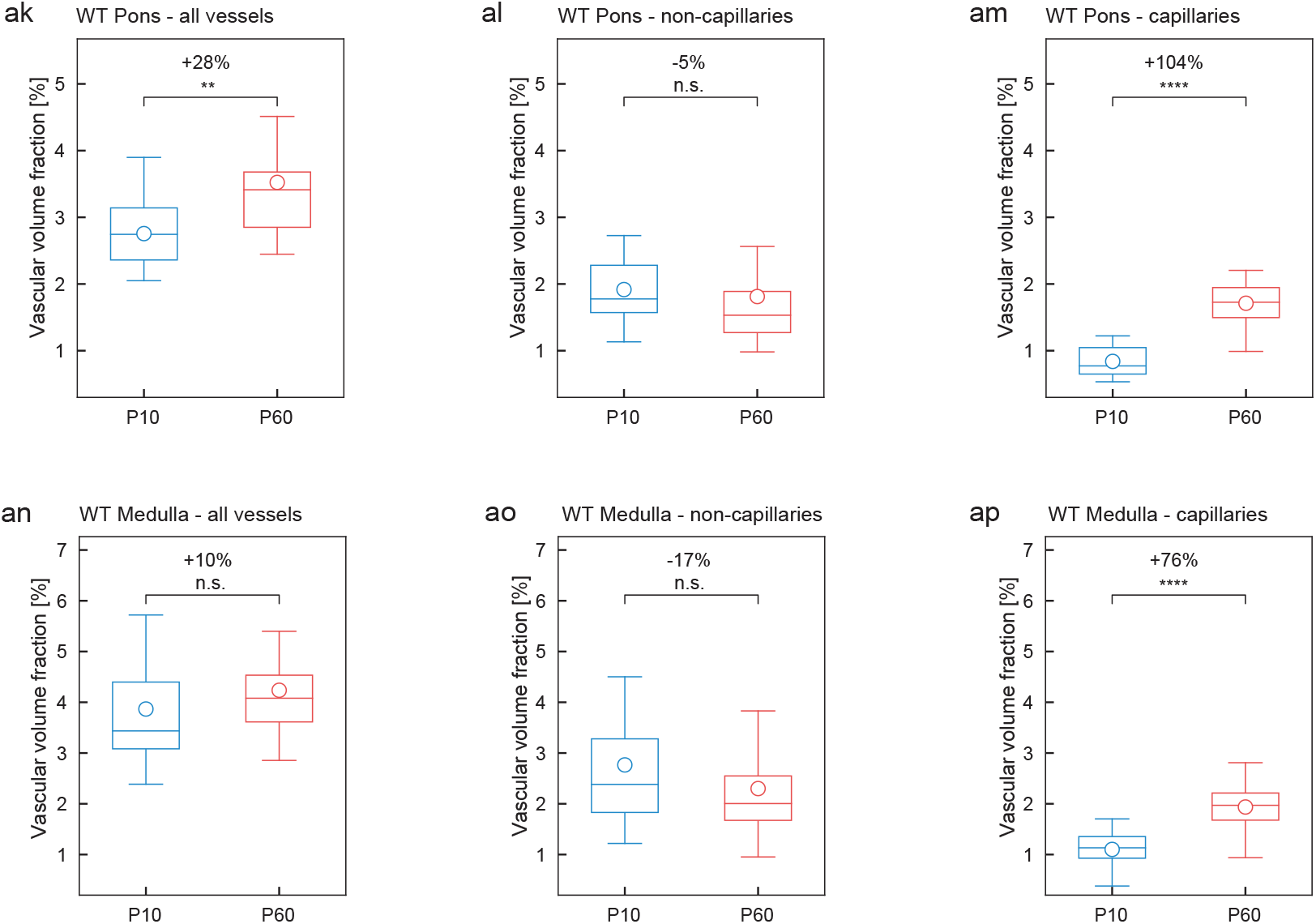
Whole-brain scan – local vascular network topology – Increased vascular volume fraction of the adult (P60) WT versus postnatal (P10) WT mouse brain mainly found at the capillary level. **a-ap** Quantification of the 3D vascular volume fraction for all vessels (**a,d,g,j,m,p,s,v,y,ab,ae,ah,ak,an**), non-capillaries (**b,e,h,k,n.q.t.w.z,ac,af,ai,al,ao**), and capillaries (**c,f,i,l,o,r,u,x,aa,ad,ag,aj,am,ap**) in P10 WT and adult (P60) WT mouse brain calculated by local topology analysis in the whole-brain scan. The increase of the vascular volume fraction for all vessels in the P60 WT animals (**a,d,g,m,p,v,y,ab,ae,ah,ak,an**) was mainly due to a significant increase at the level of capillaries (**c,f,i,l,o,r,u,x,aa,ad,ag,aj,am,ap**) in all brain regions (the anatomical regions were divided into ROI-sized section for analysis: between n = 5, dentate gyrus, and n = 126, cortex ROI-sized sections for P10 WT; n = 4, CA3, and n = 76, cortex, ROI-sized sections for P60 WT ROIs were analyzed, highly dependent on the size of the anatomical region). All data are shown as mean distributions where the open dot represents the mean. Boxplots indicate the 25% to 75% quartiles of the data. *P < 0.05, **P < 0.01, ***P < 0.001.

**Supplementary Figure 11.**
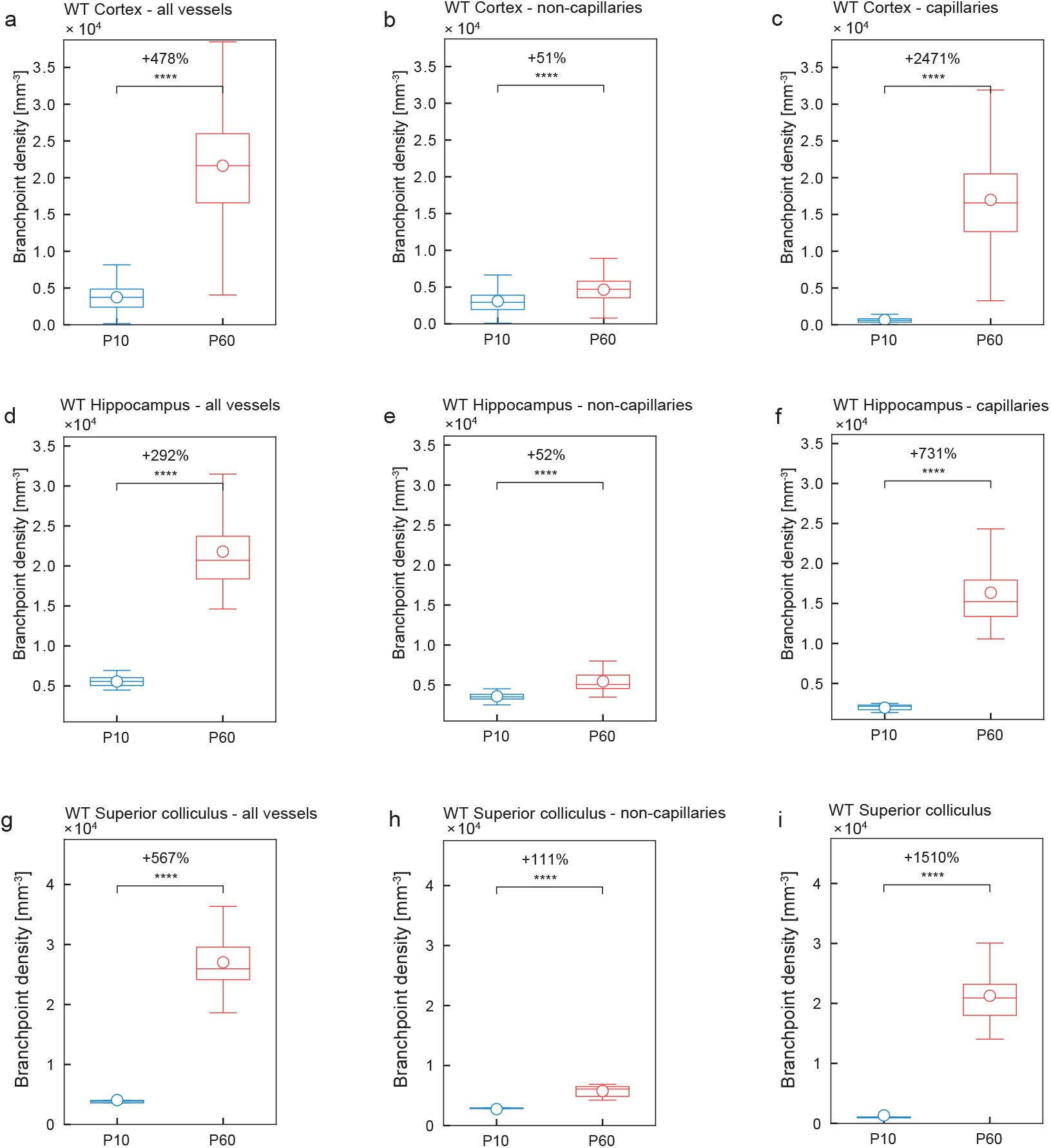
Whole-brain scan – local vascular network topology –Increased vascular branch point density in various regions of the adult (P60) WT versus the postnatal (P10) WT mouse brain. **a-i** Quantitative analysis of the branch point density for all vessels (**a,d,g**), non-capillaries (**b,e,h**), and capillaries (**c,f,i**) in the P10 WT and P60 WT mouse brain calculated by local topology analysis in the whole-brain scan. The significant increase of the branch point density for all vessels in the P60 WT animals (**a,d,g**) was mainly due to a significant increase at the level of capillaries (**c,f,i**) and in part due to a significant increase at the level of non-capillaries (**b,e,h**) (between n = 5, dentate gyrus, and n = 126, cortex ROI-sized sections for P10 WT; and n = 4, CA3, and n = 76, cortex, ROI-sized sections for P60 WT ROIs were analyzed, highly dependent on the size of the anatomical region). All data are shown as mean distributions where the open dot represents the mean. Boxplots indicate the 25% to 75% quartiles of the data. *P<0.05, **P<0.01, ***P<0.001.

**Supplementary Figure 12.**
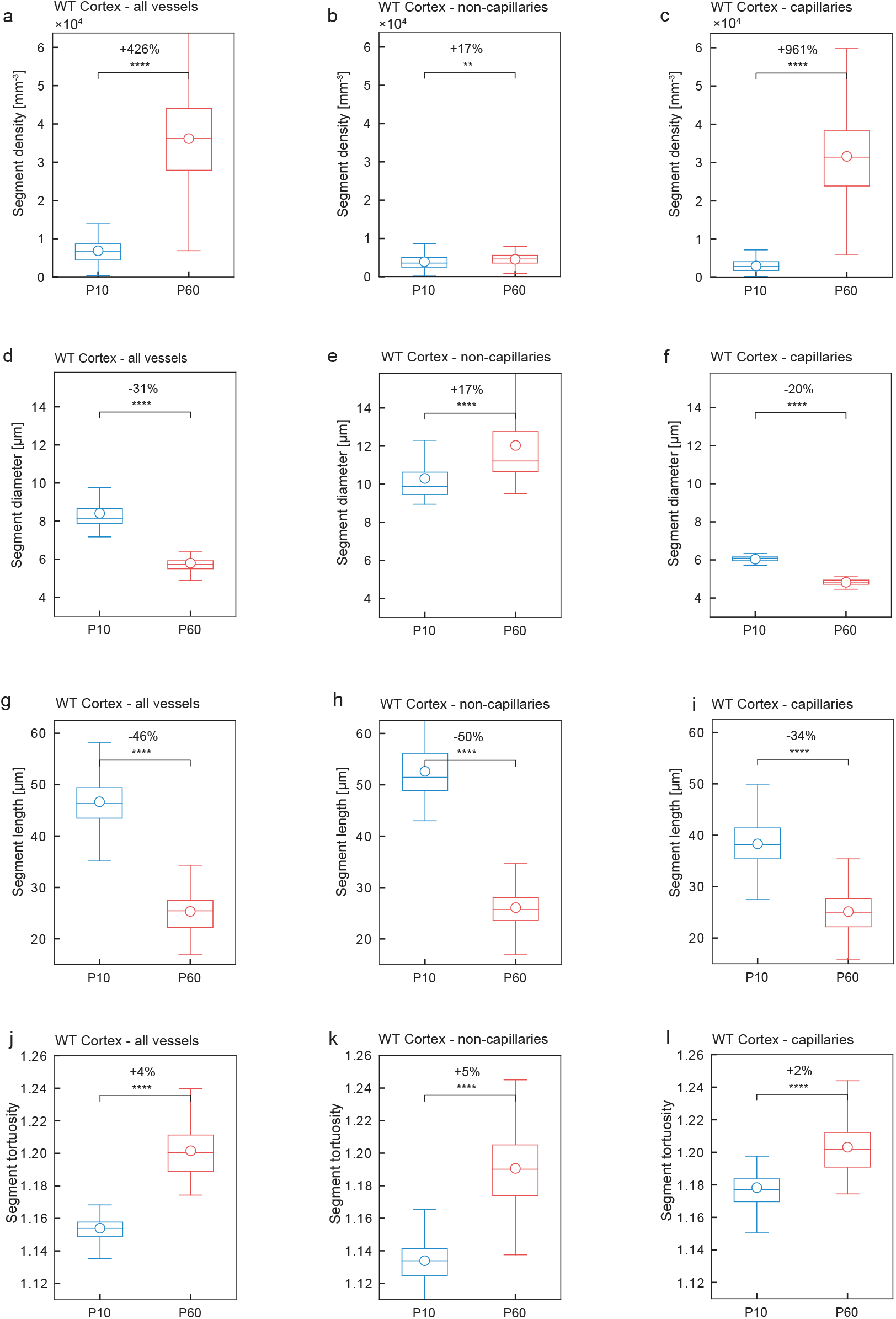
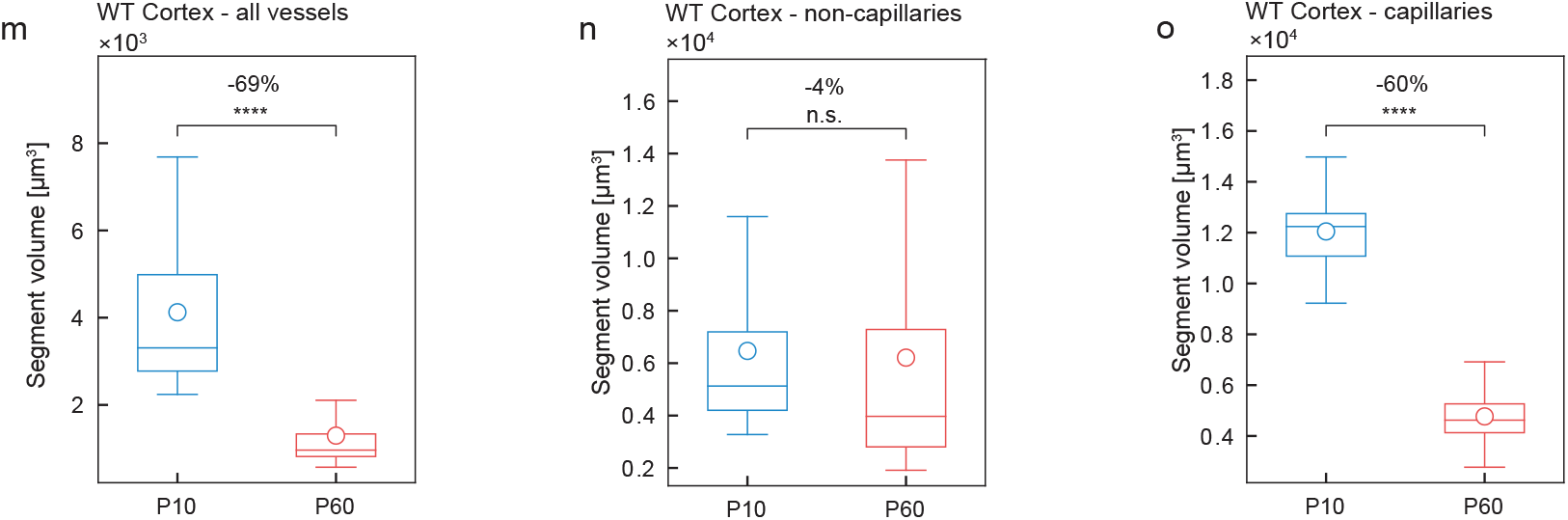
Whole-brain scan – local vascular network topology – segment density, -diameter, -length, - tortuosity, and -volume of the adult (P60) WT versus the postnatal (P10) WT mouse brain. **a-l** Quantification of the 3D vessel network parameters segment density (**a-c**), segment diameter (**d-f**), segment length (**g-i**), and segment tortuosity (**j-l**) for all vessels, non-capillaries, and capillaries in P10 WT and P60 WT cortices calculated by local morphometry analysis of the whole-brain scan. The segment density (**a**) and segment tortuosity (**j**) were significantly increased for all vessels in the cortices of P60 WT mice as compared to the P10 WT mice, whereas the segment diameter (**d**) and segment length (**g**) were significantly decreased for all vessels in the P60 WT mice as compared to the P10 WT mice. In line with the other parameters, these differences were mainly due to a highly significant increase respectively decrease at the level of capillaries (**c,f,i,l**) and in part due to an increase respectively decrease at the level of non-capillaries (**b,e,h,k**) (n = 126 ROI-sized sections for the P10 WT cortex; n = 76 for P60 WT cortex were analyzed). All data are shown as mean distributions where the open dot represents the mean. Boxplots indicate the 25% to 75% quartiles of the data. *P < 0.05, **P < 0.01, ***P < 0.001.

**Supplementary Figure 13.**
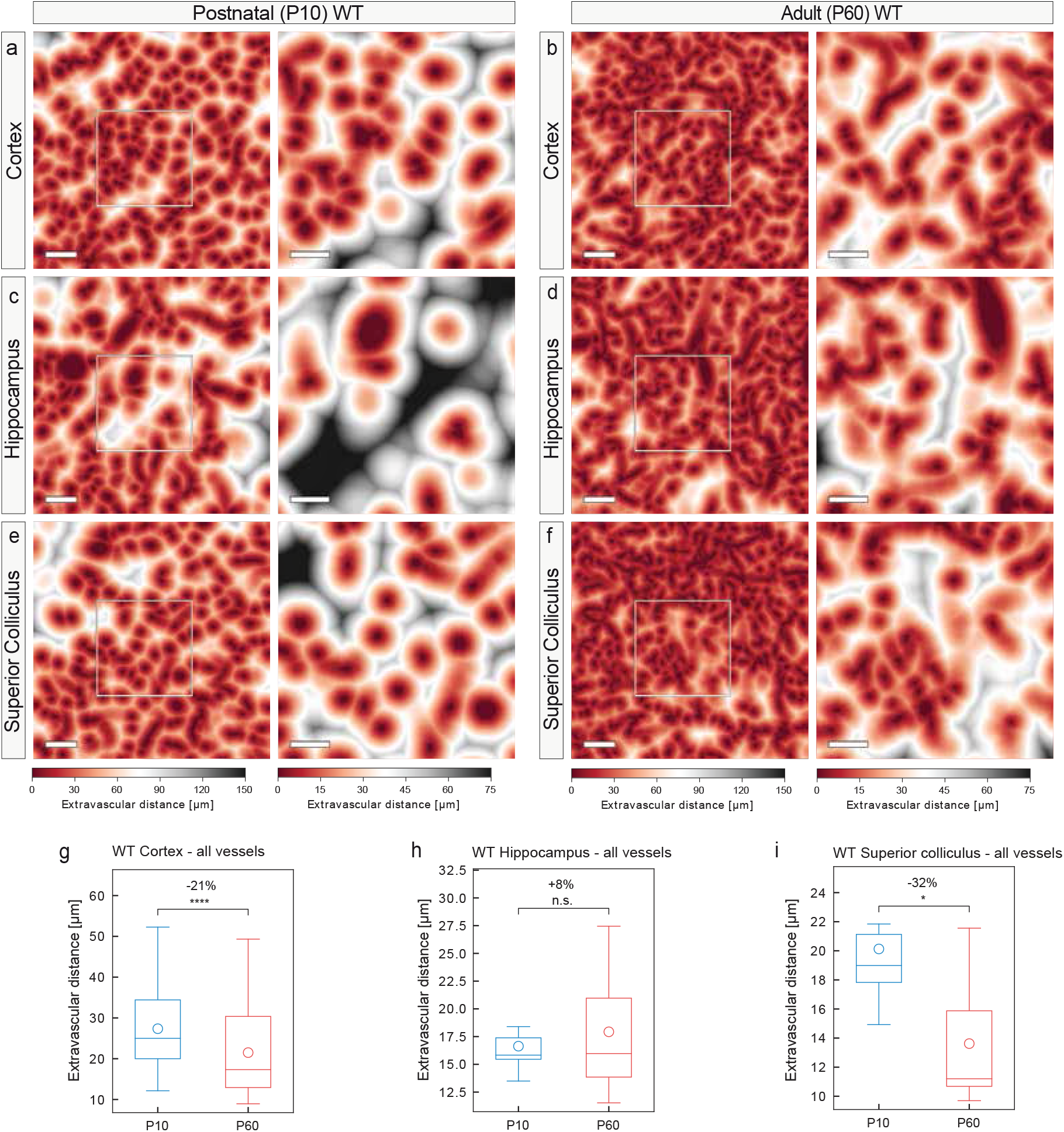
Whole-brain scan – local vascular network topology – extravascular distance is decreased in various regions of the adult (P60) WT versus the postnatal (P10) WT mouse brain. **a-f** Color map indicating the extravascular distance in the cortices, hippocampi and superior colliculi of P10 WT and P60 WT mice. Each voxel outside a vessel structure is assigned a color to depict its shortest distance to the nearest vessel structure. The reduced extravascular distance in P10 WT animals as compared to the P60 WT animals is obvious. The color bar indicates the shortest distance to the next vessel structure. **g-i** Quantification of the extravascular distance in P10 WT and P60 WT in the three brain regions calculated by global morphometry analysis. The extravascular distance in the cortices (**g**), and superior colliculi (**i**) of P60 WT animals was significantly decreased as compared to the P10 WT animals. For the hippocampi (**h**) this difference was not significant. (n = 1 for P10 WT; n = 1 for P60 WT animals were used; between n = 5, dentate gyrus, and n = 126, cortex ROI-sized sections for P10 WT; and between n = 4, CA3, and n = 76, cortex, ROI-sized sections for P60 WT ROIs were analyzed, highly dependent on the size of the anatomical region). All data are shown as mean distributions where the open dot represents the mean. Boxplots indicate the 25% to 75% quartiles of the data. *P<0.05, **P<0.01, ***P<0.001.

**Supplementary Figure 14.**
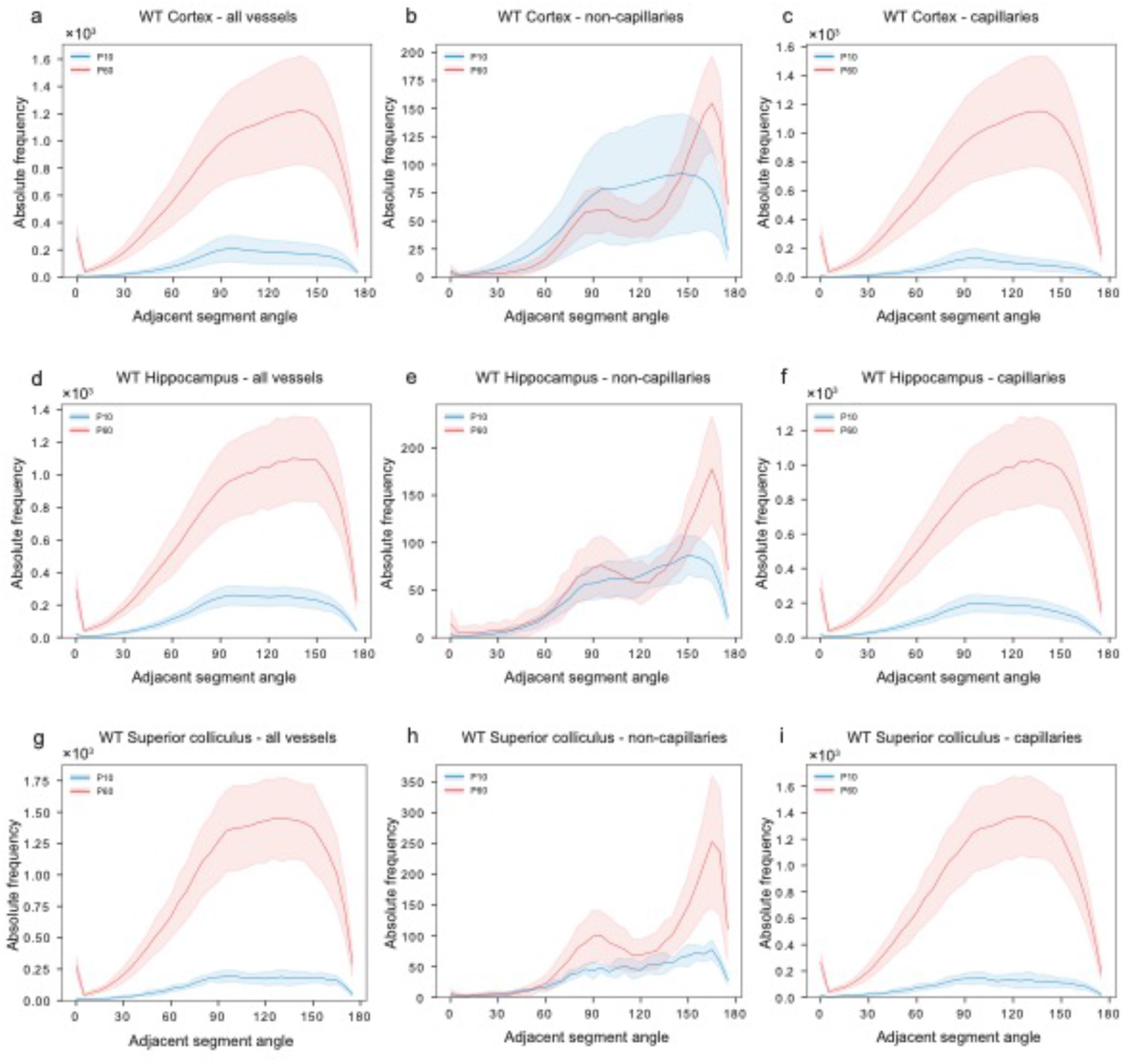
Whole-brain scan – Quantification of vessel directionality. Quantitative analysis showing the relative angles of the vascular segments emanating from a certain branch point in the cortex, hippocampus, and superior colliculus on the level of all vessels, non-capillaries and capillaries. **a-i** Analysis revealed that, comparable to the all vessel group, the capillaries in the cortex, hippocampus, and superior colliculus, derived at angles between around 75° and 150° at both the developing P10 and mature P60 stages (**c,f,i**). The non-capillaries, showed two main angles of orientation, namely around 90° and at around 170° in three brain regions examined (**b,e,h**). These two main angles of orientation were more pronounced at the P60 stage as compared to the P10 stage (**b,e,h**), and were very similar in the three brain regions, suggesting a more general underlying concept of vessel directionality/orientation. These findings, obtained from the whole-brain scans, show very similar findings to the ROI-based method (figure 10).

## SUPPLEMENTARY VIDEO

**Supplementary Video 1 Intracardial resin perfusion of a postnatal (P10) mouse**

This video shows the intracardial resin perfusion of a postnatal (P10) mouse pub. All steps are precisely explained in the “PROCEDURE” part of this manuscript.

## REFERENCES

1. Carmeliet, P. & Jain, R. K. Molecular mechanisms and clinical applications of angiogenesis. Nature 473, 298–307, doi:10.1038/nature10144 (2011).

2. Potente, M., Gerhardt, H. & Carmeliet, P. Basic and therapeutic aspects of angiogenesis. Cell 146, 873–887, doi:10.1016/j.cell.2011.08.039 (2011).

3. Walchli, T. et al. Wiring the Vascular Network with Neural Cues: A CNS Perspective. Neuron 87, 271–296, doi:10.1016/j.neuron.2015.06.038 (2015).

4. Jain, R. K. & Carmeliet, P. SnapShot: Tumor angiogenesis. Cell 149, 1408–1408.e1401, doi:10.1016/j.cell.2012.05.025 (2012).

5. Herbert, S. P. & Stainier, D. Y. Molecular control of endothelial cell behaviour during blood vessel morphogenesis. Nat Rev Mol Cell Biol 12, 551–564, doi:10.1038/nrm3176 (2011).

6. Quaegebeur, A., Lange, C. & Carmeliet, P. The neurovascular link in health and disease: molecular mechanisms and therapeutic implications. Neuron 71, 406–424, doi:10.1016/j.neuron.2011.07.013 (2011).

7. Burri, P. H., Hlushchuk, R. & Djonov, V. Intussusceptive angiogenesis: Its emergence, its characteristics, and its significance. Developmental Dynamics 231, 474–488, doi:10.1002/dvdy.20184 (2004).

8. Makanya, A. N., Hlushchuk, R. & Djonov, V. G. Intussusceptive angiogenesis and its role in vascular morphogenesis, patterning, and remodeling. Angiogenesis 12, 113–123, doi:10.1007/s10456-009-9129-5 (2009).

9. Garcia-Gomez, P. & Valiente, M. Vascular co-option in brain metastasis. Angiogenesis, doi:10.1007/s10456-019-09693-x (2019).

10. Kuczynski, E. A., Vermeulen, P. B., Pezzella, F., Kerbel, R. S. & Reynolds, A. R. Vessel co-option in cancer. Nat Rev Clin Oncol 16, 469–493, doi:10.1038/s41571-019-0181-9 (2019).

11. Angara, K., Borin, T. F. & Arbab, A. S. Vascular Mimicry: A Novel Neovascularization Mechanism Driving Anti-Angiogenic Therapy (AAT) Resistance in Glioblastoma. Transl Oncol 10, 650–660, doi:10.1016/j.tranon.2017.04.007 (2017).

12. Fernández-Cortés, M., Delgado-Bellido, D. & Oliver, F. J. Vasculogenic Mimicry: Become an Endothelial Cell “But Not So Much”. Front Oncol 9, 803–803, doi:10.3389/fonc.2019.00803 (2019).

13. Ricci-Vitiani, L. et al. Tumour vascularization via endothelial differentiation of glioblastoma stem-like cells. Nature 468, 824–828, doi:10.1038/nature09557 (2010).

14. Wang, R. et al. Glioblastoma stem-like cells give rise to tumour endothelium. Nature 468, 829–833, doi:10.1038/nature09624 (2010).

15. Cheng, L. et al. Glioblastoma stem cells generate vascular pericytes to support vessel function and tumor growth. Cell 153, 139–152, doi:10.1016/j.cell.2013.02.021 (2013).

16. Zhou, W. et al. Targeting Glioma Stem Cell-Derived Pericytes Disrupts the Blood-Tumor Barrier and Improves Chemotherapeutic Efficacy. Cell stem cell 21, 591–603.e594, doi:10.1016/j.stem.2017.10.002 (2017).

17. Coelho-Santos, V. & Shih, A. Y. Postnatal development of cerebrovascular structure and the neurogliovascular unit. Wiley Interdiscip Rev Dev Biol 9, e363–e363, doi:10.1002/wdev.363 (2020).

18. Mink, J. W., Blumenschine, R. J. & Adams, D. B. Ratio of central nervous system to body metabolism in vertebrates: its constancy and functional basis. *American Journal of Physiology-Regulatory*, Integrative and Comparative Physiology 241, R203–R212, doi:10.1152/ajpregu.1981.241.3.R203 (1981).

19. Begley, D. J. & Brightman, M. W. Structural and functional aspects of the blood-brain barrier. Prog Drug Res 61, 39–78 (2003).

20. Zlokovic, B. V. & Apuzzo, M. L. Strategies to circumvent vascular barriers of the central nervous system. Neurosurgery 43, 877–878, doi:10.1097/00006123-199810000-00089 (1998).

21. Tsai, P. S. et al. Correlations of neuronal and microvascular densities in murine cortex revealed by direct counting and colocalization of nuclei and vessels. J Neurosci 29, 14553–14570, doi:10.1523/jneurosci.3287-09.2009 (2009).

22. Blinder, P., Shih, A. Y., Rafie, C. & Kleinfeld, D. Topological basis for the robust distribution of blood to rodent neocortex. Proc Natl Acad Sci U S A 107, 12670–12675, doi:10.1073/pnas.1007239107 (2010).

23. Shih, A. Y. et al. Robust and fragile aspects of cortical blood flow in relation to the underlying angioarchitecture. *Microcirculation (New York*, N.Y. : 1994) 22, 204–218, doi:10.1111/micc.12195 (2015).

24. Harb, R., Whiteus, C., Freitas, C. & Grutzendler, J. In vivo imaging of cerebral microvascular plasticity from birth to death. J Cereb Blood Flow Metab 33, 146–156, doi:10.1038/jcbfm.2012.152 (2013).

25. Iadecola, C. The Neurovascular Unit Coming of Age: A Journey through Neurovascular Coupling in Health and Disease. Neuron 96, 17–42, doi:10.1016/j.neuron.2017.07.030 (2017).

26. Vallon, M., Chang, J., Zhang, H. & Kuo, C. J. Developmental and pathological angiogenesis in the central nervous system. Cellular and molecular life sciences : CMLS 71, 3489–3506, doi:10.1007/s00018-014-1625-0 (2014).

27. Walchli, T. et al. Quantitative assessment of angiogenesis, perfused blood vessels and endothelial tip cells in the postnatal mouse brain. Nature protocols 10, 53–74, doi:10.1038/nprot.2015.002 (2015).

28. Weller, R. O., Sharp, M. M., Christodoulides, M., Carare, R. O. & Møllgård, K. The meninges as barriers and facilitators for the movement of fluid, cells and pathogens related to the rodent and human CNS. Acta Neuropathol 135, 363–385, doi:10.1007/s00401-018-1809-z (2018).

29. Hogan, K. A., Ambler, C. A., Chapman, D. L. & Bautch, V. L. The neural tube patterns vessels developmentally using the VEGF signaling pathway. Development 131, 1503–1513, doi:10.1242/dev.01039 (2004).

30. Paredes, I., Himmels, P. & Ruiz de Almodovar, C. Neurovascular Communication during CNS Development. Dev Cell 45, 10–32, doi:10.1016/j.devcel.2018.01.023 (2018).

31. Mancuso, M. R., Kuhnert, F. & Kuo, C. J. Developmental angiogenesis of the central nervous system. Lymphat Res Biol 6, 173–180, doi:10.1089/lrb.2008.1014 (2008).

32. Fantin, A., Vieira, J. M., Plein, A., Maden, C. H. & Ruhrberg, C. The embryonic mouse hindbrain as a qualitative and quantitative model for studying the molecular and cellular mechanisms of angiogenesis. Nature protocols 8, 418–429 (2013).

33. Vasudevan, A., Long, J. E., Crandall, J. E., Rubenstein, J. L. & Bhide, P. G. Compartment-specific transcription factors orchestrate angiogenesis gradients in the embryonic brain. Nat Neurosci 11, 429–439, doi:10.1038/nn2074 (2008).

34. Tata, M., Ruhrberg, C. & Fantin, A. Vascularisation of the central nervous system. Mech Dev 138 Pt 1, 26–36, doi:10.1016/j.mod.2015.07.001 (2015).

35. Kurz, H. Cell lineages and early patterns of embryonic CNS vascularization. Cell Adh Migr 3, 205–210, doi:10.4161/cam.3.2.7855 (2009).

36. Fantin, A. et al. NRP1 acts cell autonomously in endothelium to promote tip cell function during sprouting angiogenesis. Blood 121, 2352–2362, doi:10.1182/blood-2012-05-424713 (2013).

37. Daneman, R., Zhou, L., Kebede, A. A. & Barres, B. A. Pericytes are required for blood-brain barrier integrity during embryogenesis. Nature 468, 562–566, doi:10.1038/nature09513 (2010).

38. Zeller, K., Vogel, J. & Kuschinsky, W. Postnatal distribution of Glut1 glucose transporter and relative capillary density in blood-brain barrier structures and circumventricular organs during development. Brain research. Developmental brain research 91, 200–208 (1996).

39. Walchli, T. et al. Nogo-A is a negative regulator of CNS angiogenesis. Proc Natl Acad Sci U S A 110, E1943–1952, doi:10.1073/pnas.1216203110 (2013).

40. Walchli, T. et al. Nogo-A regulates vascular network architecture in the postnatal brain. J Cereb Blood Flow Metab 37, 614–631, doi:10.1177/0271678x16675182 (2017).

41. Whiteus, C., Freitas, C. & Grutzendler, J. Perturbed neural activity disrupts cerebral angiogenesis during a postnatal critical period. Nature 505, 407–411, doi:10.1038/nature12821 (2014).

42. Sweeney, M. D., Ayyadurai, S. & Zlokovic, B. V. Pericytes of the neurovascular unit: key functions and signaling pathways. Nat Neurosci 19, 771–783, doi:10.1038/nn.4288 (2016).

43. Muoio, V., Persson, P. B. & Sendeski, M. M. The neurovascular unit - concept review. Acta Physiol (Oxf) 210, 790–798, doi:10.1111/apha.12250 (2014).

44. Eichmann, A. & Thomas, J. L. Molecular parallels between neural and vascular development. Cold Spring Harb Perspect Med 3, a006551, doi:10.1101/cshperspect.a006551 (2013).

45. Sweeney, M. D., Ayyadurai, S. & Zlokovic, B. V. Pericytes of the neurovascular unit: key functions and signaling pathways. Nature neuroscience 19, 771–783, doi:10.1038/nn.4288 (2016).

46. Yu, P. et al. FGF-dependent metabolic control of vascular development. Nature 545, 224–228, doi:10.1038/nature22322 (2017).

47. Li, X., Sun, X. & Carmeliet, P. Hallmarks of Endothelial Cell Metabolism in Health and Disease. Cell metabolism 30, 414–433, doi:10.1016/j.cmet.2019.08.011 (2019).

48. Andrae, J., Gallini, R. & Betsholtz, C. Role of platelet-derived growth factors in physiology and medicine. Genes Dev 22, 1276–1312, doi:10.1101/gad.1653708 (2008).

49. Jih, Y. J. et al. Distinct regulation of genes by bFGF and VEGF-A in endothelial cells. Angiogenesis 4, 313–321 (2001).

50. Dong, J. et al. Glioma stem cells involved in tumor tissue remodeling in a xenograft model. J Neurosurg 113, 249–260, doi:10.3171/2010.2.Jns09335 (2010).

51. Lawler, J. Thrombospondin-1 as an endogenous inhibitor of angiogenesis and tumor growth. J Cell Mol Med 6, 1–12, doi:10.1111/j.1582-4934.2002.tb00307.x (2002).

52. Lawler, P. R. & Lawler, J. Molecular basis for the regulation of angiogenesis by thrombospondin-1 and -2. Cold Spring Harb Perspect Med 2, a006627, doi:10.1101/cshperspect.a006627 (2012).

53. O’Reilly, M. S. et al. Endostatin: an endogenous inhibitor of angiogenesis and tumor growth. Cell 88, 277–285, doi:10.1016/s0092-8674(00)81848-6 (1997).

54. Dhanabal, M. et al. Angioarrestin: an antiangiogenic protein with tumor-inhibiting properties. Cancer Res 62, 3834–3841 (2002).

55. Lu, X. et al. The netrin receptor UNC5B mediates guidance events controlling morphogenesis of the vascular system. Nature 432, 179–186, doi:10.1038/nature03080 (2004).

56. Sakurai, A., Doçi, C. L. & Gutkind, J. S. Semaphorin signaling in angiogenesis, lymphangiogenesis and cancer. Cell Res 22, 23–32, doi:10.1038/cr.2011.198 (2012).

57. Eilken, H. M. & Adams, R. H. Dynamics of endothelial cell behavior in sprouting angiogenesis. Curr Opin Cell Biol 22, 617–625, doi:10.1016/j.ceb.2010.08.010 (2010).

58. Strilic, B. et al. The molecular basis of vascular lumen formation in the developing mouse aorta. Dev Cell 17, 505–515, doi:10.1016/j.devcel.2009.08.011 (2009).

59. Pitulescu, M. E. et al. Dll4 and Notch signalling couples sprouting angiogenesis and artery formation. Nat Cell Biol 19, 915–927, doi:10.1038/ncb3555 (2017).

60. Lammert, E. & Axnick, J. Vascular lumen formation. Cold Spring Harb Perspect Med 2, a006619, doi:10.1101/cshperspect.a006619 (2012).

61. Tung, J. J., Tattersall, I. W. & Kitajewski, J. Tips, Stalks, Tubes: Notch-Mediated Cell Fate Determination and Mechanisms of Tubulogenesis during Angiogenesis. Cold Spring Harb Perspect Med 2, a006601, doi:10.1101/cshperspect.a006601 (2012).

62. Wacker, A. & Gerhardt, H. Endothelial development taking shape. Curr Opin Cell Biol 23, 676–685, doi:10.1016/j.ceb.2011.10.002 (2011).

63. Mazzone, M. et al. Heterozygous deficiency of PHD2 restores tumor oxygenation and inhibits metastasis via endothelial normalization. Cell 136, 839–851, doi:10.1016/j.cell.2009.01.020 (2009).

64. Dela Paz, N. G., Melchior, B. & Frangos, J. A. Shear stress induces Gα(q/11) activation independently of G protein-coupled receptor activation in endothelial cells. American journal of physiology. Cell physiology 312, C428–C437, doi:10.1152/ajpcell.00148.2016 (2017).

65. Takuwa, N. et al. Tumor-suppressive sphingosine-1-phosphate receptor-2 counteracting tumor-promoting sphingosine-1-phosphate receptor-1 and sphingosine kinase 1 - Jekyll Hidden behind Hyde. Am J Cancer Res 1, 460–481 (2011).

66. Mendelson, K., Evans, T. & Hla, T. Sphingosine 1-phosphate signalling. Development 141, 5–9, doi:10.1242/dev.094805 (2014).

67. Gaengel, K. et al. The sphingosine-1-phosphate receptor S1PR1 restricts sprouting angiogenesis by regulating the interplay between VE-cadherin and VEGFR2. Dev Cell 23, 587–599, doi:10.1016/j.devcel.2012.08.005 (2012).

68. Jin, Z. G. et al. Ligand-independent activation of vascular endothelial growth factor receptor 2 by fluid shear stress regulates activation of endothelial nitric oxide synthase. Circ Res 93, 354–363, doi:10.1161/01.Res.0000089257.94002.96 (2003).

69. Nigro, P., Abe, J. & Berk, B. C. Flow shear stress and atherosclerosis: a matter of site specificity. Antioxid Redox Signal 15, 1405–1414, doi:10.1089/ars.2010.3679 (2011).

70. Baeyens, N. et al. Defective fluid shear stress mechanotransduction mediates hereditary hemorrhagic telangiectasia. Journal of Cell Biology 214, 807–816, doi:10.1083/jcb.201603106 (2016).

71. Franco, C. A. et al. Non-canonical Wnt signalling modulates the endothelial shear stress flow sensor in vascular remodelling. eLife 5, e07727–e07727, doi:10.7554/eLife.07727 (2016).

72. Pries, A. R. & Secomb, T. W. Modeling structural adaptation of microcirculation. Microcirculation 15, 753–764, doi:10.1080/10739680802229076 (2008).

73. Pries, A. R., Secomb, T. W. & Gaehtgens, P. Structural adaptation and stability of microvascular networks: theory and simulations. Am J Physiol 275, H349–360, doi:10.1152/ajpheart.1998.275.2.H349 (1998).

74. Hjelmeland, A. B., Lathia, J. D., Sathornsumetee, S. & Rich, J. N. Twisted tango: brain tumor neurovascular interactions. Nat Neurosci 14, 1375–1381, doi:10.1038/nn.2955 (2011).

75. Carmeliet, P., De Smet, F., Loges, S. & Mazzone, M. Branching morphogenesis and antiangiogenesis candidates: tip cells lead the way. Nat Rev Clin Oncol 6, 315–326, doi:10.1038/nrclinonc.2009.64 (2009).

76. Frahm, K. A., Nash, C. P. & Tobet, S. A. Endocan immunoreactivity in the mouse brain: method for identifying nonfunctional blood vessels. Journal of immunological methods 398-399, 27–32, doi:10.1016/j.jim.2013.09.005 (2013).

77. Geudens, I. & Gerhardt, H. Coordinating cell behaviour during blood vessel formation. Development 138, 4569–4583, doi:10.1242/dev.062323 (2011).

78. Blanco, R. & Gerhardt, H. VEGF and Notch in tip and stalk cell selection. Cold Spring Harb Perspect Med 3, a006569, doi:10.1101/cshperspect.a006569 (2013).

79. Phng, L. K. & Gerhardt, H. Angiogenesis: a team effort coordinated by notch. Dev Cell 16, 196–208, doi:10.1016/j.devcel.2009.01.015 (2009).

80. Noguera-Troise, I. et al. Blockade of Dll4 inhibits tumour growth by promoting non-productive angiogenesis. Nature 444, 1032–1037, doi:10.1038/nature05355 (2006).

81. Thurston, G., Noguera-Troise, I. & Yancopoulos, G. D. The Delta paradox: DLL4 blockade leads to more tumour vessels but less tumour growth. Nat Rev Cancer 7, 327–331, doi:10.1038/nrc2130 (2007).

82. Yan, M. et al. Chronic DLL4 blockade induces vascular neoplasms. Nature 463, E6–7, doi:10.1038/nature08751 (2010).

83. Lobov, I. B. et al. Delta-like ligand 4 (Dll4) is induced by VEGF as a negative regulator of angiogenic sprouting. Proc Natl Acad Sci U S A 104, 3219–3224, doi:10.1073/pnas.0611206104 (2007).

84. Ridgway, J. et al. Inhibition of Dll4 signalling inhibits tumour growth by deregulating angiogenesis. Nature 444, 1083–1087, doi:10.1038/nature05313 (2006).

85. Hiruma, T., Nakajima, Y. & Nakamura, H. Development of pharyngeal arch arteries in early mouse embryo. Journal of Anatomy 201, 15–29, doi:10.1046/j.1469-7580.2002.00071.x (2002).

86. Hiruma, T. & Hirakow, R. Formation of the pharyngeal arch arteries in the chick embryo. Observations of corrosion casts by scanning electron microscopy. Anat Embryol (Berl) 191, 415–423, doi:10.1007/bf00304427 (1995).

87. Kim, B.-G. et al. CXCL12-CXCR4 signalling plays an essential role in proper patterning of aortic arch and pulmonary arteries. Cardiovasc Res 113, 1677–1687, doi:10.1093/cvr/cvx188 (2017).

88. Adamson, S. L. et al. Interactions between trophoblast cells and the maternal and fetal circulation in the mouse placenta. Dev Biol 250, 358–373, doi:10.1016/s0012-1606(02)90773-6 (2002).

89. Gest, T. R. & Carron, M. A. Embryonic origin of the caudal mesenteric artery in the mouse. Anat Rec A Discov Mol Cell Evol Biol 271, 192–201, doi:10.1002/ar.a.10022 (2003).

90. Iwagaki, T., Suzuki, T. & Nakashima, T. Development and regression of cochlear blood vessels in fetal and newborn mice. Hear Res 145, 75–81, doi:10.1016/s0378-5955(00)00075-7 (2000).

91. Carraro, M. & Harrison, R. V. Degeneration of stria vascularis in age-related hearing loss; a corrosion cast study in a mouse model. Acta Otolaryngol 136, 385–390, doi:10.3109/00016489.2015.1123291 (2016).

92. Carraro, M., Park, A. H. & Harrison, R. V. Partial corrosion casting to assess cochlear vasculature in mouse models of presbycusis and CMV infection. Hear Res 332, 95–103, doi:10.1016/j.heares.2015.11.010 (2016).

93. Carraro, M. et al. Cytomegalovirus (CMV) Infection Causes Degeneration of Cochlear Vasculature and Hearing Loss in a Mouse Model. J Assoc Res Otolaryngol 18, 263–273, doi:10.1007/s10162-016-0606-4 (2017).

94. Hossler, F. E., Lametschwandtner, A., Kao, R. & Finsterbusch, F. Microvascular architecture of mouse urinary bladder described with vascular corrosion casting, light microscopy, SEM, and TEM. Microsc Microanal 19, 1428–1435, doi:10.1017/S143192761301341X (2013).

95. Peão, M. N., Aguas, A. P., de Sá, C. M. & Grande, N. R. Neoformation of blood vessels in association with rat lung fibrosis induced by bleomycin. Anat Rec 238, 57–67, doi:10.1002/ar.1092380108 (1994).

96. Peáo, M. N., Aguas, A. P., de Sá, C. M. & Grande, N. R. Identification of vascular sphincters at the junction between alveolar capillaries and pulmonary venules of the mouse lung. Anat Rec 241, 383–390, doi:10.1002/ar.1092410313 (1995).

97. Ackermann, M. et al. Effects of nintedanib on the microvascular architecture in a lung fibrosis model. Angiogenesis 20, 359–372, doi:10.1007/s10456-017-9543-z (2017).

98. Gibney, B. C. et al. Structural and functional evidence for the scaffolding effect of alveolar blood vessels. Exp Lung Res 43, 337–346, doi:10.1080/01902148.2017.1368739 (2017).

99. Dahl, E. The fine structure of intracerebral vessels. Z Zellforsch Mikrosk Anat 145, 577–586, doi:10.1007/bf00306725 (1973).

100. Levesque, M. J., Cornhill, J. F. & Nerem, R. M. Vascular casting. A new method for the study of the arterial endothelium. Atherosclerosis 34, 457–467, doi:10.1016/0021-9150(79)90070-4 (1979).

101. Duvernoy, H. M., Delon, S. & Vannson, J. L. Cortical blood vessels of the human brain. Brain Res Bull 7, 519–579, doi:10.1016/0361-9230(81)90007-1 (1981).

102. Reina-De La Torre, F., Rodriguez-Baeza, A. & Sahuquillo-Barris, J. Morphological characteristics and distribution pattern of the arterial vessels in human cerebral cortex: a scanning electron microscope study. Anat Rec 251, 87–96, doi:10.1002/(SICI)1097-0185(199805)251:1<87::AID-AR14>3.0.CO;2-7 (1998).

103. Zagórska-Swiezy, K., Litwin, J. A., Gorczyca, J., Pityński, K. & Miodoński, A. J. Arterial supply and venous drainage of the choroid plexus of the human lateral ventricle in the prenatal period as revealed by vascular corrosion casts and SEM. Folia Morphol (Warsz) 67, 209–213 (2008).

104. Heinzer, S. et al. Hierarchical microimaging for multiscale analysis of large vascular networks. Neuroimage 32, 626–636, doi:10.1016/j.neuroimage.2006.03.043 (2006).

105. Krucker, T., Lang, A. & Meyer, E. P. New polyurethane-based material for vascular corrosion casting with improved physical and imaging characteristics. Microsc Res Tech 69, 138–147, doi:10.1002/jemt.20263 (2006).

106. Sangiorgi, S. et al. Arterial and microvascular supply of cerebral hemispheres in the nude mouse revealed using corrosion casting and scanning electron microscopy. Journal of anatomy 232, 739–746, doi:10.1111/joa.12791 (2018).

107. Quintana, D. D. et al. The cerebral angiome: High resolution MicroCT imaging of the whole brain cerebrovasculature in female and male mice. NeuroImage 202, 116109, doi:10.1016/j.neuroimage.2019.116109 (2019).

108. Heinzer, S. et al. Novel three-dimensional analysis tool for vascular trees indicates complete micro-networks, not single capillaries, as the angiogenic endpoint in mice overexpressing human VEGF(165) in the brain. Neuroimage 39, 1549–1558, doi:10.1016/j.neuroimage.2007.10.054 (2008).

109. Krucker, T., Schuler, A., Meyer, E. P., Staufenbiel, M. & Beckmann, N. Magnetic resonance angiography and vascular corrosion casting as tools in biomedical research: application to transgenic mice modeling Alzheimer’s disease. Neurol Res 26, 507–516, doi:10.1179/016164104225016281 (2004).

110. Meyer, E. P., Ulmann-Schuler, A., Staufenbiel, M. & Krucker, T. Altered morphology and 3D architecture of brain vasculature in a mouse model for Alzheimer’s disease. Proc Natl Acad Sci U S A 105, 3587–3592, doi:10.1073/pnas.0709788105 (2008).

111. Walker, E. J., Shen, F., Young, W. L. & Su, H. Cerebrovascular casting of the adult mouse for 3D imaging and morphological analysis. Journal of visualized experiments : JoVE, e2958–e2958, doi:10.3791/2958 (2011).

112. Walker, E. J. et al. Arteriovenous malformation in the adult mouse brain resembling the human disease. Ann Neurol 69, 954–962, doi:10.1002/ana.22348 (2011).

113. Risser, L. et al. From homogeneous to fractal normal and tumorous microvascular networks in the brain. Journal of cerebral blood flow and metabolism : official journal of the International Society of Cerebral Blood Flow and Metabolism 27, 293–303, doi:10.1038/sj.jcbfm.9600332 (2007).

114. Sangiorgi, S. et al. Early-stage microvascular alterations of a new model of controlled cortical traumatic brain injury: 3D morphological analysis using scanning electron microscopy and corrosion casting. 118, 763, doi:10.3171/2012.11.Jns12627 (2013).

115. Ohtake, M., Morino, S., Kaidoh, T. & Inoué, T. Three-dimensional structural changes in cerebral microvessels after transient focal cerebral ischemia in rats: scanning electron microscopic study of corrosion casts. Neuropathology 24, 219–227, doi:10.1111/j.1440-1789.2004.00560.x (2004).

116. Rodríguez-Baeza, A., Reina-de la Torre, F., Poca, A., Martí, M. & Garnacho, A. Morphological features in human cortical brain microvessels after head injury: a three-dimensional and immunocytochemical study. Anat Rec A Discov Mol Cell Evol Biol 273, 583–593, doi:10.1002/ar.a.10069 (2003).

117. Zhang, F., Wen, Y. & Guo, X. CRISPR/Cas9 for genome editing: progress, implications and challenges. Hum Mol Genet 23, R40–46, doi:10.1093/hmg/ddu125 (2014).

118. Gerhardt, H. et al. Neuropilin-1 is required for endothelial tip cell guidance in the developing central nervous system. Dev Dyn 231, 503–509, doi:10.1002/dvdy.20148 (2004).

119. Gerhardt, H. et al. VEGF guides angiogenic sprouting utilizing endothelial tip cell filopodia. J Cell Biol 161, 1163–1177, doi:10.1083/jcb.200302047 (2003).

120. Beckmann, N. et al. Age-dependent cerebrovascular abnormalities and blood flow disturbances in APP23 mice modeling Alzheimer’s disease. J Neurosci 23, 8453–8459 (2003).

121. Pitulescu, M. E., Schmidt, I., Benedito, R. & Adams, R. H. Inducible gene targeting in the neonatal vasculature and analysis of retinal angiogenesis in mice. Nature protocols 5, 1518–1534, doi:10.1038/nprot.2010.113 (2010).

122. O’Connor, J. P. et al. Quantifying antivascular effects of monoclonal antibodies to vascular endothelial growth factor: insights from imaging. Clin Cancer Res 15, 6674–6682, doi:10.1158/1078-0432.Ccr-09-0731 (2009).

123. Paganin, D., Mayo, S. C., Gureyev, T. E., Miller, P. R. & Wilkins, S. W. Simultaneous phase and amplitude extraction from a single defocused image of a homogeneous object. Journal of Microscopy 206, 33–40, doi:10.1046/j.1365-2818.2002.01010.x (2002).

124. Marone, F. & Stampanoni, M. Regridding reconstruction algorithm for real-time tomographic imaging. J Synchrotron Radiat 19, 1029–1037, doi:10.1107/s0909049512032864 (2012).

125. Miettinen, A., Oikonomidis, I. V., Bonnin, A. & Stampanoni, M. NRStitcher: non-rigid stitching of terapixel-scale volumetric images. Bioinformatics 35, 5290–5297, doi:10.1093/bioinformatics/btz423 (2019).

126. Otsu, N. A Threshold Selection Method from Gray-Level Histograms. *IEEE Transactions on Systems*, Man, and Cybernetics 9, 62–66, doi:10.1109/TSMC.1979.4310076 (1979).

127. Huang, L.-K. & Wang, M.-J. J. Image thresholding by minimizing the measures of fuzziness. Pattern Recognition 28, 41–51, doi:https://doi.org/10.1016/0031-3203(94)E0043-K (1995).

128. Prewitt, J. M. S. & Mendelsohn, M. L. THE ANALYSIS OF CELL IMAGES*. Annals of the New York Academy of Sciences 128, 1035–1053, doi:10.1111/j.1749-6632.1965.tb11715.x (1966).

129. Li, C. H. & Tam, P. K. S. An iterative algorithm for minimum cross entropy thresholding. Pattern Recogn. Lett. 19, 771–776, doi:10.1016/s0167-8655(98)00057-9 (1998).

130. Lee, T. C., Kashyap, R. L. & Chu, C. N. Building Skeleton Models via 3-D Medial Surface Axis Thinning Algorithms. CVGIP: Graphical Models and Image Processing 56, 462–478, doi:https://doi.org/10.1006/cgip.1994.1042 (1994).

131. Calvin R. Maurer, J., Qi, R. & Raghavan, V. A Linear Time Algorithm for Computing Exact Euclidean Distance Transforms of Binary Images in Arbitrary Dimensions. IEEE Trans. Pattern Anal. Mach. Intell. 25, 265–270, doi:10.1109/tpami.2003.1177156 (2003).

132. Suhadolnik, A., Petrišič, J. & Kosel, F. An anchored discrete convolution algorithm for measuring length in digital images. Measurement 42, 1112–1117, doi:https://doi.org/10.1016/j.measurement.2009.04.005 (2009).

133. Tukey, J. W. Exploratory data analysis. (Addison-Wesley Pub. Co., 1977).

134. Provost, J. et al. 3-D ultrafast Doppler imaging applied to the noninvasive mapping of blood vessels in vivo. IEEE Trans Ultrason Ferroelectr Freq Control 62, 1467–1472, doi:10.1109/TUFFC.2015.007032 (2015).

135. Demené, C. et al. 4D microvascular imaging based on ultrafast Doppler tomography. Neuroimage 127, 472–483, doi:10.1016/j.neuroimage.2015.11.014 (2016).

136. Pathak, A. P., Kim, E., Zhang, J. & Jones, M. V. Three-dimensional imaging of the mouse neurovasculature with magnetic resonance microscopy. PLoS One 6, e22643–e22643, doi:10.1371/journal.pone.0022643 (2011).

137. Hartung, M. P., Grist, T. M. & François, C. J. Magnetic resonance angiography: current status and future directions. J Cardiovasc Magn Reson 13, 19–19, doi:10.1186/1532-429X-13-19 (2011).

138. Kelch, I. D. et al. Organ-wide 3D-imaging and topological analysis of the continuous microvascular network in a murine lymph node. Sci Rep 5, 16534, doi:10.1038/srep16534 (2015).

139. Marien, K. M. et al. Development and Validation of a Histological Method to Measure Microvessel Density in Whole-Slide Images of Cancer Tissue. PLoS One 11, e0161496–e0161496, doi:10.1371/journal.pone.0161496 (2016).

140. Bukenya, F., Nerissa, C., Serres, S., Pardon, M. C. & Bai, L. An automated method for segmentation and quantification of blood vessels in histology images. Microvasc Res 128, 103928, doi:10.1016/j.mvr.2019.103928 (2020).

141. Oren, R. et al. Whole Organ Blood and Lymphatic Vessels Imaging (WOBLI). Scientific reports 8, 1412–1412, doi:10.1038/s41598-018-19663-w (2018).

142. Kennel, P., Teyssedre, L., Colombelli, J. & Plouraboué, F. Toward quantitative three-dimensional microvascular networks segmentation with multiview light-sheet fluorescence microscopy. J Biomed Opt 23, 1–14, doi:10.1117/1.Jbo.23.8.086002 (2018).

143. Di Giovanna, A. P. et al. Tailored Sample Mounting for Light-Sheet Fluorescence Microscopy of Clarified Specimens by Polydimethylsiloxane Casting. Frontiers in neuroanatomy 13, 35–35, doi:10.3389/fnana.2019.00035 (2019).

144. Lugo-Hernandez, E. et al. 3D visualization and quantification of microvessels in the whole ischemic mouse brain using solvent-based clearing and light sheet microscopy. J Cereb Blood Flow Metab 37, 3355–3367, doi:10.1177/0271678x17698970 (2017).

145. Todorov, M. I. et al. Machine learning analysis of whole mouse brain vasculature. Nat Methods 17, 442–449, doi:10.1038/s41592-020-0792-1 (2020).

146. Kirst, C. et al. Mapping the Fine-Scale Organization and Plasticity of the Brain Vasculature. Cell 180, 780–795.e725, doi:10.1016/j.cell.2020.01.028 (2020).

147. Susaki, E. A. & Ueda, H. R. Whole-body and Whole-Organ Clearing and Imaging Techniques with Single-Cell Resolution: Toward Organism-Level Systems Biology in Mammals. Cell Chem Biol 23, 137–157, doi:10.1016/j.chembiol.2015.11.009 (2016).

148. Zagorchev, L. et al. Micro computed tomography for vascular exploration. J Angiogenes Res 2, 7, doi:10.1186/2040-2384-2-7 (2010).

149. Blery, P. et al. Vascular imaging with contrast agent in hard and soft tissues using microcomputed-tomography. Journal of Microscopy 262, 40–49, doi:10.1111/jmi.12339 (2016).

150. Starosolski, Z. et al. Ultra High-Resolution In vivo Computed Tomography Imaging of Mouse Cerebrovasculature Using a Long Circulating Blood Pool Contrast Agent. Scientific Reports 5, 10178, doi:10.1038/srep10178 (2015).

151. Gore, A. V., Monzo, K., Cha, Y. R., Pan, W. & Weinstein, B. M. Vascular development in the zebrafish. Cold Spring Harb Perspect Med 2, a006684, doi:10.1101/cshperspect.a006684 (2012).

152. Dorr, A., Sled, J. G. & Kabani, N. Three-dimensional cerebral vasculature of the CBA mouse brain: a magnetic resonance imaging and micro computed tomography study. Neuroimage 35, 1409–1423, doi:10.1016/j.neuroimage.2006.12.040 (2007).

153. Figueiredo, G., Boll, H., Kramer, M., Groden, C. & Brockmann, M. A. In Vivo X-Ray Digital Subtraction and CT Angiography of the Murine Cerebrovasculature Using an Intra-Arterial Route of Contrast Injection. American Journal of Neuroradiology 33, 1702, doi:10.3174/ajnr.A3071 (2012).

154. Ben-Zvi, A. et al. Mfsd2a is critical for the formation and function of the blood-brain barrier. Nature 509, 507–511, doi:10.1038/nature13324 (2014).

155. Vanlandewijck, M. et al. A molecular atlas of cell types and zonation in the brain vasculature. Nature 554, 475–480, doi:10.1038/nature25739 (2018).

156. Kalucka, J. et al. Single-Cell Transcriptome Atlas of Murine Endothelial Cells. Cell 180, 764–779.e720, doi:10.1016/j.cell.2020.01.015 (2020).

157. Goveia, J. et al. An Integrated Gene Expression Landscape Profiling Approach to Identify Lung Tumor Endothelial Cell Heterogeneity and Angiogenic Candidates. Cancer cell 37, 21–36.e13, doi:10.1016/j.ccell.2019.12.001 (2020).

158. Sawamiphak, S., Ritter, M. & Acker-Palmer, A. Preparation of retinal explant cultures to study ex vivo tip endothelial cell responses. Nature protocols 5, 1659–1665, doi:10.1038/nprot.2010.130 (2010).

159. Moyon, D., Pardanaud, L., Yuan, L., Bréant, C. & Eichmann, A. Plasticity of endothelial cells during arterial-venous differentiation in the avian embryo. Development 128, 3359–3370 (2001).

160. Villa, N. et al. Vascular expression of Notch pathway receptors and ligands is restricted to arterial vessels. Mech Dev 108, 161–164, doi:10.1016/s0925-4773(01)00469-5 (2001).

161. Seki, T., Yun, J. & Oh, S. P. Arterial endothelium-specific activin receptor-like kinase 1 expression suggests its role in arterialization and vascular remodeling. Circ Res 93, 682–689, doi:10.1161/01.Res.0000095246.40391.3b (2003).

162. Cui, X. et al. Venous Endothelial Marker COUP-TFII Regulates the Distinct Pathologic Potentials of Adult Arteries and Veins. Scientific Reports 5, 16193, doi:10.1038/srep16193 (2015).

163. You, L. R. et al. Suppression of Notch signalling by the COUP-TFII transcription factor regulates vein identity. Nature 435, 98–104, doi:10.1038/nature03511 (2005).

164. Wang, H. U., Chen, Z. F. & Anderson, D. J. Molecular distinction and angiogenic interaction between embryonic arteries and veins revealed by ephrin-B2 and its receptor Eph-B4. Cell 93, 741–753 (1998).

165. Malcontenti-Wilson, C., Chan, L., Nikfarjam, M., Muralidharan, V. & Christophi, C. Vascular targeting agent Oxi4503 inhibits tumor growth in a colorectal liver metastases model. J Gastroenterol Hepatol 23, e96–e104, doi:10.1111/j.1440-1746.2007.04899.x (2008).

166. Kaidoh, T., Yasugi, T. & Uehara, Y. The microvasculature of the 7,12-dimethylbenz(a)anthracene (DMBA)-induced rat mammary tumour. Virchows Archiv A 418, 111–117, doi:10.1007/BF01600286 (1991).

167. Chakhoyan, A. et al. Validation of vessel size imaging (VSI) in high-grade human gliomas using magnetic resonance imaging, image-guided biopsies, and quantitative immunohistochemistry. Scientific Reports 9, 2846, doi:10.1038/s41598-018-37564-w (2019).

168. Burrell, J. S. et al. MRI measurements of vessel calibre in tumour xenografts: Comparison with vascular corrosion casting. Microvascular Research 84, 323–329, doi:https://doi.org/10.1016/j.mvr.2012.08.001 (2012).

169. ACSF for LSPS. Cold Spring Harbor Protocols 2012, pdb.rec071944, doi:10.1101/pdb.rec071944 (2012).

170. Rust, R. et al. Nogo-A targeted therapy promotes vascular repair and functional recovery following stroke. Proceedings of the National Academy of Sciences 116, 14270, doi:10.1073/pnas.1905309116 (2019).

171. Joly, S., Dejda, A., Rodriguez, L., Sapieha, P. & Pernet, V. Nogo-A inhibits vascular regeneration in ischemic retinopathy. Glia 66, 2079–2093, doi:10.1002/glia.23462 (2018).

172. Seiler, S., Di Santo, S. & Widmer, H. R. Nogo-A Neutralization Improves Graft Function in a Rat Model of Parkinson’s Disease. Front Cell Neurosci 10, 87, doi:10.3389/fncel.2016.00087 (2016).

173. Weber, B., Keller, A. L., Reichold, J. & Logothetis, N. K. The microvascular system of the striate and extrastriate visual cortex of the macaque. Cerebral cortex 18, 2318–2330, doi:10.1093/cercor/bhm259 (2008).

174. Cassot, F., Lauwers, F., Lorthois, S., Puwanarajah, P. & Duvernoy, H. Scaling laws for branching vessels of human cerebral cortex. Microcirculation 16, 331–344, 332 p following 344, doi:10.1080/10739680802662607 (2009).

175. Fenrich, K. K., Zhao, E. Y., Wei, Y., Garg, A. & Rose, P. K. Isolating specific cell and tissue compartments from 3D images for quantitative regional distribution analysis using novel computer algorithms. J Neurosci Methods 226, 42–56, doi:10.1016/j.jneumeth.2014.01.011 (2014).

176. Simonen, M. et al. Systemic deletion of the myelin-associated outgrowth inhibitor Nogo-A improves regenerative and plastic responses after spinal cord injury. Neuron 38, 201–211 (2003).

177. Tatu, L. & Vuillier, F. Structure and vascularization of the human hippocampus. Front Neurol Neurosci 34, 18–25, doi:10.1159/000356440 (2014).

178. Marinković, S., Milisavljević, M. & Puskas, L. Microvascular anatomy of the hippocampal formation. Surg Neurol 37, 339–349, doi:10.1016/0090-3019(92)90001-4 (1992).

